# The Core Human Fecal Metabolome

**DOI:** 10.1101/2021.05.08.442269

**Authors:** Jacob J. Haffner, Mitchelle Katemauswa, Thérèse S. Kagone, Ekram Hossain, David Jacobson, Karina Flores, Adwaita R. Parab, Alexandra J. Obregon-Tito, Raul Y. Tito, Luis Marin Reyes, Luzmila Troncoso-Corzo, Emilio Guija-Poma, Nicolas Meda, Hélène Carabin, Tanvi P. Honap, Krithivasan Sankaranarayanan, Cecil M. Lewis, Laura-Isobel McCall

## Abstract

Among the biomolecules at the center of human health and molecular biology is a system of molecules that defines the human phenotype known as the metabolome. Through an untargeted metabolomic analysis of fecal samples from human individuals from Africa and the Americas—the birthplace and the last continental expansion of our species, respectively—we present the characterization of the core human fecal metabolome. The majority of detected metabolite features were ubiquitous across populations, despite any geographic, dietary, or behavioral differences. Such shared metabolite features included hyocholic acid and cholesterol. However, any characterization of the core human fecal metabolome is insufficient without exploring the influence of industrialization. Here, we show chemical differences along an industrialization gradient, where the degree of industrialization correlates with metabolomic changes. We identified differential metabolite features like leucyl-leucine dipeptides and urobilin as major metabolic correlates of these behavioral shifts. Our results indicate that industrialization significantly influences the human fecal metabolome, but diverse human lifestyles and behavior still maintain a core human fecal metabolome. This study represents the first characterization of the core human fecal metabolome through untargeted analyses of populations along an industrialization gradient.

## Manuscript

Metabolites fit as the final stage of biology’s central dogma: DNA transcribed into RNA translated into proteins which enzymatically interact, form, and shed into small molecules as part of the biochemical pathways of metabolism^1–3^. For this study, we define a metabolite as any small molecule (<1,500 Da) involved in biochemical pathways and the metabolome as the collection of these small molecules within a biological system^3–5^. Using the definition from the Human Metabolome Database, these endogenous metabolites (synthesized by the host) are supplemented by exogenous small molecules (acquired from external sources, such as cosmetics, medication, dietary sources, and pollution)^6^. The human metabolome thus contains both endogenous and exogenous metabolites, representing the nexus of genetic and environmental influences^5, 7, 8^.

Characterizing the fecal metabolome requires an understanding of how it is influenced by different factors, such as industrialization^9, 10^. Broadly, industrialization is a series of economic and technological changes relating to the processing and distribution of resources that ultimately cause a shift from agrarian to industrial societies^11, 12^. Such changes generally involve an increase in manufactured products compared to agriculture/hunting and other raw products, a greater percentage of workers employed in industrial workplaces over agriculture, and changes in the physical landscape such as increased construction of built environments^13^. Industrialization is often linked with urbanization, which refers to social and demographic shifts increasing population size and density within a settlement^14^. These processes lead to industrialized-urban populations exhibiting denser populations^14^, reduced environmental exposures^15, 16^, an indirect relationship with food sources^17, 18^, and dietary shifts^18, 19^ compared to non-industrial rural populations. Moreover, industrialization leads to significant biological changes; industrialization reduces microbial diversity^16, 20–22^, increases allergic diseases^23, 24^ and asthma^25^, and heightens susceptibility to illnesses such as inflammatory bowel disease^26, 27^. Investigations into industrially-caused metabolomic shifts have identified differences based in amino acids, amines, sphingolipids, and hexoses, among others^19, 21, 28, 29^. Some studies detailed human fecal metabolomes by comparing rural and urban populations and found differences in levels of acylcarnitines, amino acids, and short-chain fatty acids^28–30^. However, such studies employed targeted/semi-targeted metabolomic approaches and/or sampled a single human population^19, 21, 28–30^. As a result, these studies do not represent ranges of human diversity and behavior, highlighting the need for broader investigations of the human fecal metabolome in terms of geographic range and chemical space.

Here, we performed untargeted liquid chromatography mass spectrometry (LC-MS)-based metabolomics on fecal samples obtained from six human populations from diverse geographic regions (Figure 1a; Table 1; Supplementary Table 1). These populations included male and female children and adults. Our sampled populations were given an industrialization score corresponding with their degree of industrialization from a scale of one (most urban-industrialized) to four (least industrialized; see Materials and Methods for details on score calculation). Importantly, we included two populations with similar degrees of industrialization but from distinct continents, to control for any geographic confounders - this key aspect has not been considered in prior industrialization-focused metabolomics research. Our populations include: Norman (USA; industrialization score 1); Guayabo (Peru; industrialization score 2); Tambo de Mora (Peru; industrialization score 2); Boulkiemdé (Burkina Faso; industrialization score 3); Tunapuco (Peru; industrialization score 3); and Matses (Peru; industrialization score 4).

**Figure 1.**
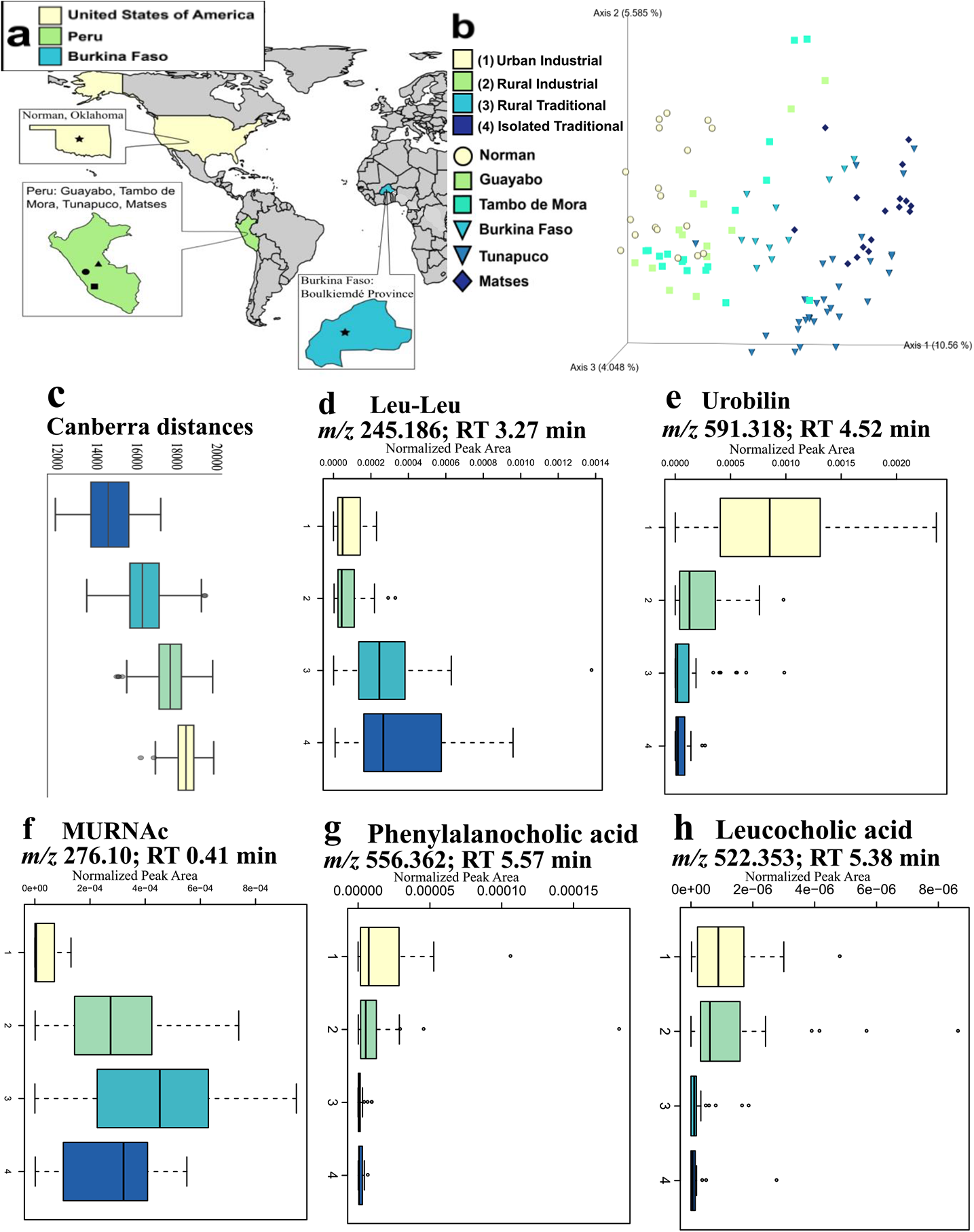
Fecal metabolomic profiles follow an industrialization gradient. **a**, Sampling sites. Tan star: Norman; Green circle: Guayabo; Green square: Tambo de Mora; green triangle: Tunapuco; Blue star: Boulkiemdé.Matses left unmarked due to privacy concerns. **b** Principal coordinate analysis (Canberra distance metric) depicts industrialization gradient. Colored by industrialization score and shape-coded by population. **c**, Calculated Canberra distances follow an industrialization gradient. Colored by industrialization score. Color key from **b** applies to **c-i. d-f**, Normalized abundances of features identified by random forest differing by industrialization score: **d**, Leucyl-leucine (leu-leu), associated with non-industrialized populations. *m/z* 245.186; RT 3.27 min. **e**, Urobilin, associated with industrialized populations. *m/z* 591.318; RT 4.16 min. **f**, Feature structurally similar to *N*-acetylmuramic acid (MURNAc) associated with semi-industrialized and non-industrialized populations, *m/z* 276.108; RT 0.41 min. **g-h**, Normalized abundances of novel amino acid-conjugated bile acids depict an industrialization gradient: **g**, Phenylalanocholic acid. *m/z* 556.36; RT 5.57 min. **h**, Leucocholic acid. *m/z* 522.353; RT 5.38 min.

**Table 1.** Sampled population metadata.

| Population | Abbreviation | Geographic Origin | Industrialization Score | Sample size (n) | Age distribution |  |  | Sex distribution |  |
| --- | --- | --- | --- | --- | --- | --- | --- | --- | --- |
|  |  |  |  |  | 1-17 years | 18-44 years | 45+ years | Female | Male |
| Total |  |  | 1-4 | 105 | 39 | 48 | 17 | 50 | 34 |
| Norman | NO | Norman, Oklahoma, United States | 1 | 18 | 0 | 18 | 0 | 7 | 11 |
| Guayabo | GU | Guayabo, Peru, South America | 2 | 13 | 5 | 7 | 5 | 11 | 0 |
| Tambo de Mora | TM | Tambo de Mora District, Peru, South America | 2 | 17 | 8 | 5 | 3 | 8 | 1 |
| Boulkiemdé | BF | Boulkiemdé Province, Burkina Faso, Africa | 3 | 11 | 0 | 6 | 5 | 6 | 5 |
| Tunapuco | HCO | Andean Highlands, Peru, South America | 3 | 30 | 15 | 9 | 2 | 17 | 9 |
| Matses | SM | Peruvian Amazon, South America | 4 | 16 | 11 | 3 | 2 | 8 | 8 |

Fecal metabolomes of these populations followed an industrialization gradient, where populations exhibited similar metabolomes based on the degree of industrialization. Principal Coordinate Analysis (PCoA) demonstrated significant differences in overall metabolome composition based on industrialization score (Figure 1b-c; Permutational Multivariate Analysis of Variance (PERMANOVA) p=0.001, R^2^=0.140; Canberra distance). Moreover, the gradient seen in our data indicates industrialization had a stronger influence on metabolic similarity between populations than geographic origin (Figure 1c; ANOVA industrialization score p=0.046, effect size (eta2)=0.08; ANOVA geographic origin p=0.245, eta2=0.01). This overshadowing of the influence of geography demonstrates the profound influence industrialization has on human molecular biology. Our findings concur with prior studies demonstrating industrialization’s role in shaping the human microbiome^31–34^, the built environment microbiome^16, 35^, the built environment metabolome^35^, and the plasma metabolome^21^. Additionally, the observation of industrialization outweighing the effects of geographic origin is novel for human fecal metabolomics analyses, but concurs with findings from human fecal microbiome studies^31–34^. To the best of our knowledge, this is the first study to illustrate the industrialization gradient in the human fecal metabolome—the intuitive path for revealing the key chemistry of the distal gut.

To determine the factors driving this clustering of metabolite profiles by industrialization degree, we employed a random forest machine learning algorithm. This random forest analysis analyzed the top 1,000 most abundant metabolites features to identify the 30 most differential metabolite features by degree of industrialization (Table 2; Supplementary Figure 1). Only two of the most abundant differential features could be annotated: leucyl-leucine (mass-to-charge ratio (*m/z*): 245.186; retention time (RT): 3.27 min; Kruskal-Wallis p=8.73e-09) and urobilin (*m/z*: 591.318; RT: 4.52 min; Kruskal-Wallis p=4.45e-07). Leucyl-leucine (leu-leu) abundance was most associated with non-industrial populations, while urobilin abundance was strongly associated with industrialized populations (Figure 1d-e). Leu-leu is a leucine dipeptide previously recognized as a human metabolite in a study comparing fecal metabolomes of individuals with and without colorectal cancer, where leu-leu showed 99% prevalence across both control and colorectal cancer groups^36^. While leu-leu has not been mentioned in previous industrialization-focused studies of human fecal metabolomes, increased abundance of leucine was noted in fecal metabolomes of urban Nigerian adults as compared to rural adults^28^, contrasting with the non-industrial association of leu-leu in our data. The second annotated differential metabolite feature, urobilin, is formed from the metabolic breakdown of hemoglobin^37^. While previous industrialization-focused fecal metabolomics studies did not report this metabolite, urobilin has been identified as a common metabolite in human urine and fecal metabolomes^38, 39^. Importantly, urobilin abundance is affected by host diet and behavior^40^, with increased abundance seen in populations consuming diets rich in animal fat, proteins, and carbohydrates^41^, such as those seen in industrialized populations. Urobilin’s association with industrialized human fecal metabolomes highlights the relationship between diet and industrialization, and reinforces the industrialization gradient seen in our results. While only two of the 30 differential metabolite features could be directly annotated, two other features were structurally related to *N*-acetylmuramic acid (MURNAc), as determined by molecular networking^42^. These two features were elevated in semi-industrialized and non-industrialized populations (Figure 1f). MURNAc is a biopolymer component comprising the peptidoglycan layers of bacterial cell walls and a prior study identified reduced abundance of MURNAc in human fecal metabolomes of individuals with lupus^43^. Lupus is an autoimmune disease whose susceptibility is associated with increased environmental exposures that are common to industrialized populations^44–46^. Since MURNAc is a component of all bacterial cell walls, its association with differential industrialization metabolites also suggests these unannotated metabolite features are bacteria-derived or related. All in all, our results identified several metabolite features that are heavily influenced by industrialization.

**Table 2.**
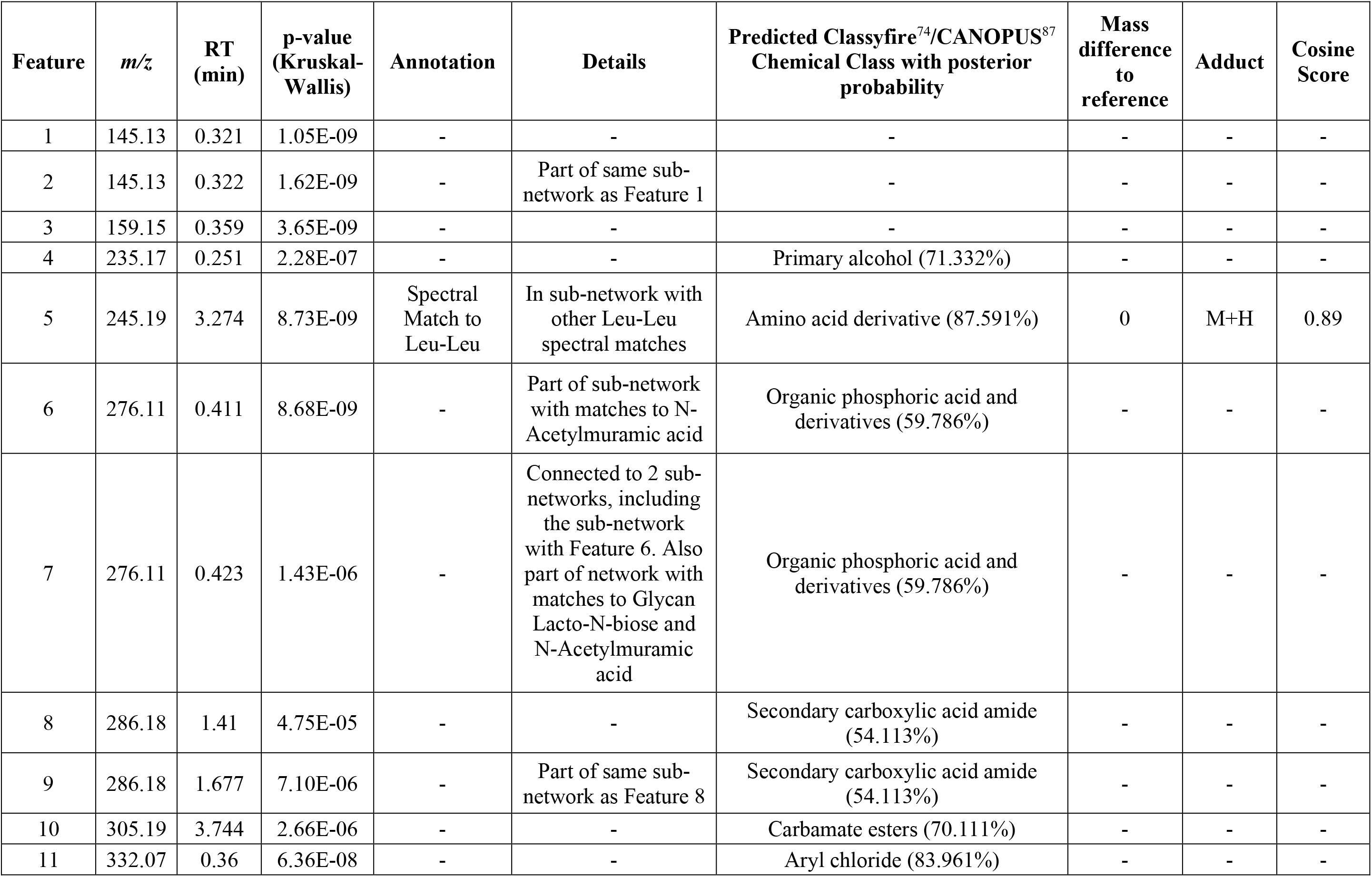
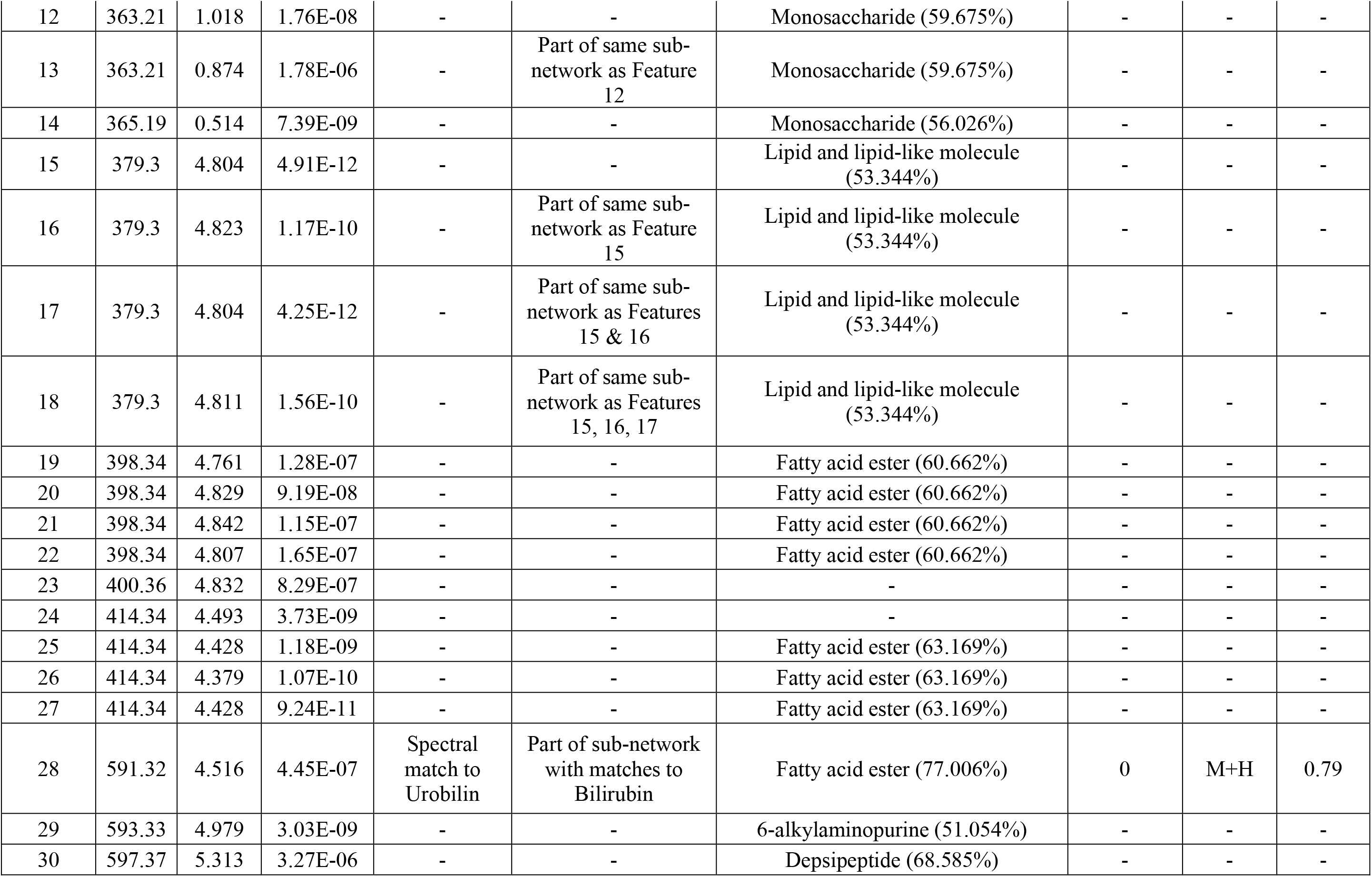
Top 30 most differential metabolite features as determined by random forest classifier.

Recent research has revealed novel amino acid-conjugated bile acids that are produced by the microbiota^47–49^. Given their enrichment in patients with inflammatory bowel disease^48^, which is associated with industrialization processes^26, 27^, we investigated their distribution across our industrialization gradient. Overall, ten of the 12 total identified amino acid-conjugated bile acids demonstrated a striking increase with industrialization. Such differential amino acid-conjugated bile acids include phenylalanocholic acid (Kruskal-Wallis p=1.9e-6), leucocholic acid (Kruskal-Wallis p=1.69e-7), leucine-conjugated chenodeoxycholic acid (CHDCA) (Kruskal-Wallis p=0.04), tyrosocholic acid (Kruskal-Wallis p=7.71e-3), tyrosine-conjugated deoxycholic acid (Kruskal-Wallis p=1.61e-5), glutamate-conjugated CHDCA (Kruskal-Wallis p=1.69e-7), tryptophan-conjugated CHDCA (Kruskal-Wallis p=4.9e-7), aspartate-conjugated CHDCA (Kruskal-Wallis p=1.13e-5), histidine-conjugated CHDCA (Kruskal-Wallis p=6.41e-3), and histidine-conjugated cholic acid (Kruskal-Wallis p=0.04) (Figure 1g-h, Supplementary Figure 2). However, two amino acid-conjugate bile acids, aspartate-conjugated cholic acid (Kruskal-Wallis p=0.05) and threonine-conjugated CHDCA (Kruskal-Wallis p=0.4), were not enriched in industrialized populations and did not display any statistically significant differences based on industrialization score. The functional role of these amino acid-conjugated bile acids in health is currently unknown, though our results further support a link to industrialization-associated inflammatory diseases.

These differences notwithstanding, our study identified many similarities across populations. A total of 36,324 metabolite features were detected in our samples with 28,288 features being shared across our populations (Figure 2a-b). Our sampled populations are considerably different from each other with strong dietary, behavioral, and geographic differences and, together, represent distinct realms of human experience and diversity. Thus, metabolite features common to these markedly separate populations likely constitute a core human metabolome shared by all humans, even if their abundances vary. To identify these common metabolite features, we filtered our data using three different levels of stringency, limiting metabolite features to those found in at least six individuals from each population, half our samples, or all our samples. The six-sample filtering retained 16,609 total metabolite features across all populations, the half-sample filtering retained 6,205 total metabolite features with 6,008 shared by all populations, and the all-sample filtering retained 1,080 total shared metabolite features. To validate that this high number of shared features was not an artefact of our data processing pipeline, we further filtered our data to only include features shared between two different processing methods: gap-filled and non-gap-filled. Additionally, features annotated to researcher-derived molecules such as DEET were excluded from our list of the core fecal metabolome. These retained common metabolite features included chemical groups like indoles, steroids, lactones, and fatty acyls (Supplementary Table 2; Supplementary Figure 3). Dipeptides included threonylphenylalanine (*m/z* 267.134; RT 0.48 min), valylvaline (*m/z* 217.155; RT 0.45 min), and isoleucylproline (*m/z* 229.155; RT 0.55 min). Shared bile acids include hyocholic acid (*m/z* 158.154; RT 4.78 min; primary bile acid involved with absorbing and transporting diet fats and drugs to the liver^50^) and lithocholic acid (*m/z* 323.273; RT 6.84 min; secondary bile acid commonly found in feces^51^ and associated with irritable bowel syndrome^52^). Fatty acid examples include 3-hydroxydodecanoic acid (*m/z* 199.169; RT 7.10 min; medium chain fatty acid associated with fatty acid metabolic disorders, potentially acquired from the microbial genera *Pseudomonas, Moraxella,* and *Acinetobacter*^53, 54^) and palmitoleic acid (*m/z* 237.001; RT 6.42 min; fatty acid commonly found in human adipose tissue and associated with obesity^55^; also acquired in diet from human breast milk^56^). Additional metabolites include cholesterol (*m/z* 369.352; RT 10.5 min; essential sterol found in animals^6^), methionine (*m/z* 105.058; RT 0.33 min; amino acid), and leucine enkephalin (*m/z* 336.192; RT 3.21 min; peptide naturally produced in animal brains, including humans^6, 57^). While a number of these shared metabolite features listed above provide key biological functions, some metabolites appear to be derived from dietary sources. An example of a metabolite possibly acquired from food products includes conjugated linoleic acid (*m/z* 263.24; RT 6.68 min; commonly found in meat and dairy products, also recognized for anti-inflammatory capabilities^6, 58^).

**Figure 2.**
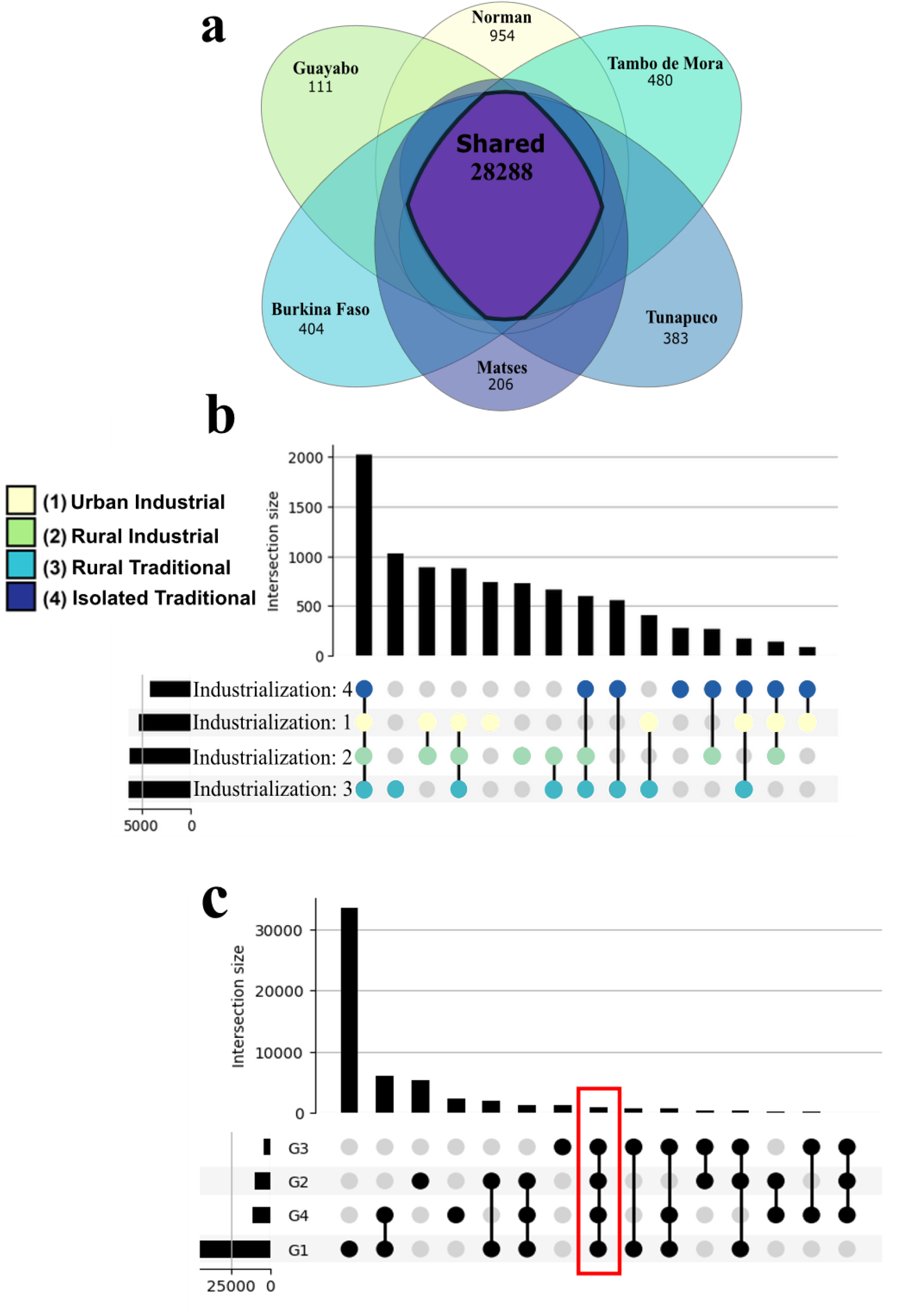
The core human metabolome. **a**, Metabolic feature overlap across study populations. **b**, UpSet plot of industrialization score sets indicate strong similarity of metabolomic profiles. **c**, UpSet plot of ReDU co-analysis datasets sorted by MS instrument. G1 is ThermoFisher Scientific Q Exactive (*n*=696); G2 is Bruker Impact (*n*=447); G3 is Bruker maXis (*n*=143); G4 is the dataset from this study (*n*=105). The co-analysis illustrates overlap across the datasets, despite instrumental differences. Colored box highlights intersection of all datasets (855 total metabolite feature).

To explore associations between the core human fecal metabolome and gut microbiome profiles, Spearman’s rho correlation coefficients were calculated for the core metabolites and identified microbial operational taxonomic units (OTUs) derived from clustering sequences. Moderate to strong correlations were noted between 604 core human fecal metabolites and gut microbe pairs (Figure 3; Table 3; Supplementary Table 3; Supplementary Files 1-2), though no metabolite-microbe pair reported a correlation coefficient exceeding ±0.6. Most of these metabolites had both positive and negative correlations with different microbes. Many microbes were correlated with multiple metabolites, on average seven. Likewise, on average, each of these 604 metabolites was correlated with 12 microbes, indicating high connectivity between fecal microbiome and metabolome. Extreme examples include methyl-oxindole and bilirubin, which each reported over 40 correlations to microbial OTUs, while Val-Met or Thr-Pro each had only one correlated microbe. Methyl-oxindole is a tryptophan derivative; metabolism of tryptophan by the gut bacteria has been extensively studied, though methyl-oxindole is less well characterized^59^. Most correlated microbes were categorized as Clostridia (48% of total nodes) or Bacteroidia classes (16% of total nodes) (Figure 3d), which respectively, have reduced and increased abundance in industrialized populations^19, 32^. Indeed, urobilin had higher abundance in industrialized than in non-industrialized populations in our analysis, and most of its strong correlations were with Clostridia microbes while most of its negative correlations were with Bacteroidia. Bilirubin, which was enriched in industrialized populations, was negatively correlated with Bacteroidia. This pattern highlights interactions between the core human fecal metabolome and the gut microbiome, especially as they are influenced by processes like industrialization.

**Figure 3.**
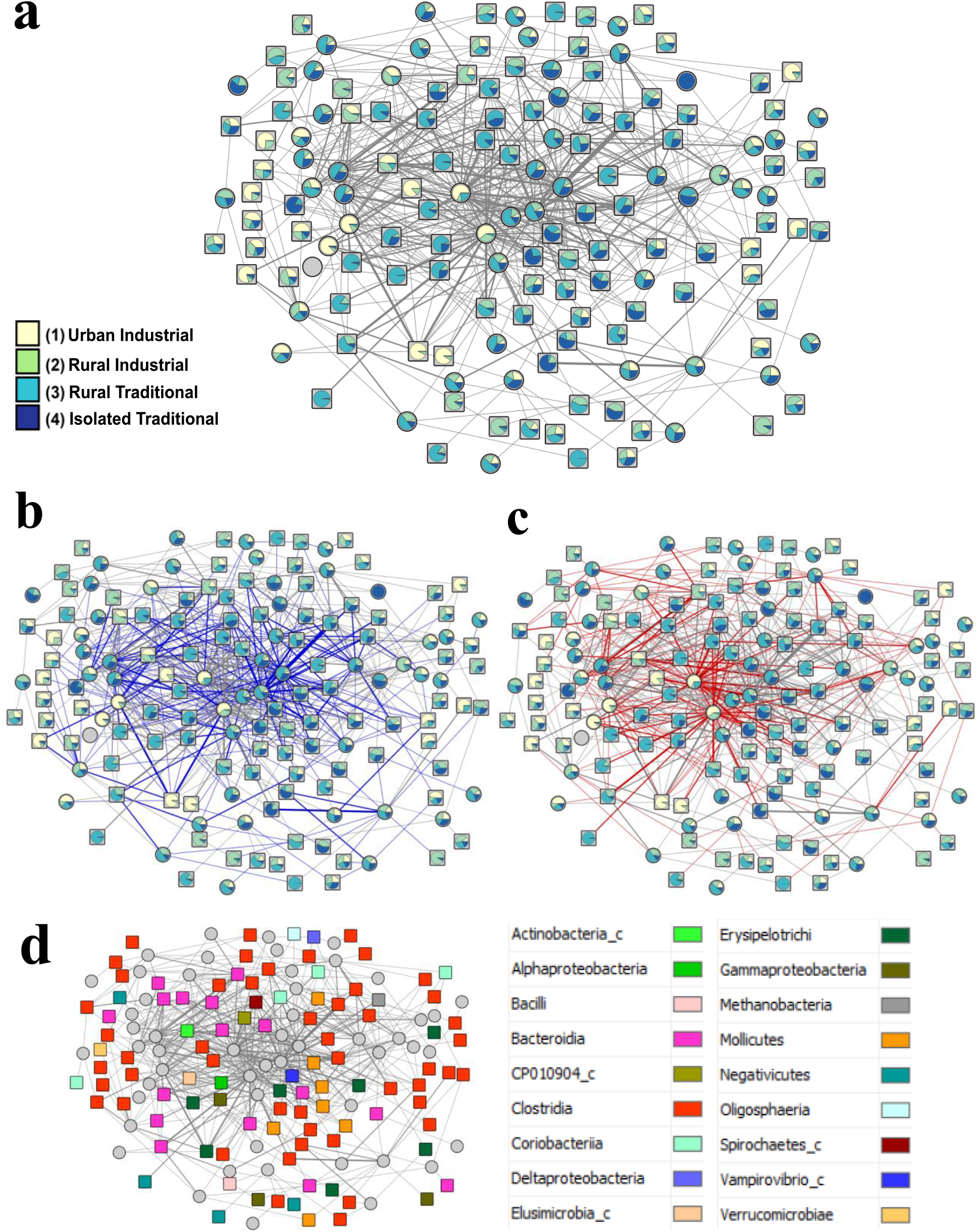
Correlation network of metabolite-microbe associations across an urbanization gradient. **a,** Correlation network. **a-d**, Metabolites are circles and microbial OTUs are squares. **a-c**, Thin edges represent weak correlations (±0.3 - ±0.399), medium edges are moderate correlations (±0.4 - ±0.499), and thick edges are moderately strong correlations (>±0.5). Pie charts represent node abundance across industrialization groups. Color key from **a** applies to **b-c**. **b,** Positive correlations are blue. **c,** Negative correlations are red. **d**, Microbes are color-coded by microbial class. Clostridia is the predominant microbial class represented in the correlation network, followed by Bacteroidia and Mollicutes.

**Table 3.** Metabolite-Microbe correlations. Only metabolite-microbe pairs greater than ±0.3 were included. Major microbial class was determined by calculating the percentage of microbial nodes connected to the respective metabolite node, focused on classes with greater than 10% for each respective metabolite.

| *m/z* | RT (min) | Metabolite Name | Number of Edges | Major Taxa Class (%) |
| --- | --- | --- | --- | --- |
| 137.046 | 0.38 | Hypoxanthine | 25 | Bacteroidia (24%);<br>Clostridia (24%);<br>Mollicutes (20%) |
| 139.05 | 0.31 | Nicotinamide N-oxide | 1 | Clostridia (100%) |
| 148.076 | 3.22 | 3-methyl-2-oxindole | 41 | Clostridia (43.9%) |
| 158.154 | 4.78 | Hyocholic acid | 12 | Clostridia (50%) |
| 177.055 | 4.13 | 3-Hydroxy-4-methoxycinnamic acid | 11 | Clostridia (72.7%) |
| 177.164 | 5.91 | 2-Butanone, 4-(2,6,6-trimethyl-2-cyclohexen-1-yl)- | 3 | Clostridia (66.6%);<br>Bacteroidia (33.3%) |
| 181.072 | 0.81 | Paraxanthine | 39 | Clostridia (25.6%);<br>Bacteroidia (23.1%) |
| 195.065 | 3.01 | trans-Ferulic acid | 9 | Clostridia (66.7%) |
| 197.117 | 3.10 | Loliolide | 15 | Clostridia (53.3%);<br>Bacteroidia (26.7%) |
| 199.169 | 5.35 | 3-Hydroxydodecanoic acid | 7 | Clostridia (28.6%) |
| 204.087 | 3.53 | N-Acetyl-D-mannosamine | 8 | Bacteroidia (37.5%);<br>Clostridia (37.5%) |
| 208.097 | 2.84 | L-Phenylalanine, N-acetyl- | 24 | Clostridia (62.5%) |
| 217.122 | 0.462 | Thr-Pro | 1 | Bacteroidia (100%) |
| 217.155 | 0.45 | Val-Val | 10 | Bacteroidia (40%);<br>Clostridia (20%) |
| 220.118 | 0.56 | Pantothenic acid | 4 | Bacteroidia (75%);<br>Clostridia (25%) |
| 227.201 | 5.94 | Myristoleic acid | 9 | Clostridia (44.4%);<br>Mollicutes (33.3%) |
| 229.155 | 0.78 | Ile-Pro | 4 | Clostridia (50%);<br>Bacteroidia (25%);<br>Coriobacteriia (25%) |
| 231.171 | 2.82 | Val-Ile | 37 | Clostridia (35.1%);<br>Bacteroidia (18.9%) |
| 237.221 | 6.49 | cis-9-Hexadecenoic acid | 7 | Clostridia (42.9%);<br>Negativicutes (28.6%) |
| 239.102 | 0.44 | Gly-Tyr | 5 | Erysipelotrichi (40%) |
| 245.098 | 2.39 | Biotin | 3 | Bacteroidia (33.3%);<br>Clostridia (33.3%);<br>Deltaproteobacteria<br>(33.3%) |
| 245.186 | 2.40 | Leu-Leu | 23 | Bacteroidia (43.5%);<br>Clostridia (21.7%) |
| 247.145 | 1.22 | Lenticin | 14 | Bacteroidia (50%) |
| 249.126 | 0.56 | Val-Met | 1 | Clostridia (100%) |
| 263.237 | 6.68 | Conjugated linoleic acid (9E,11E) | 14 | Clostridia (78.6%) |
| 263.237 | 7.52 | Conjugated linoleic acid (10E,12Z) | 1 | Clostridia (100%) |
| 267.134 | 0.48 | Thr-Phe | 2 | Clostridia (50%);<br>Deltaproteobacteria (50%) |
| 272.171 | 0.32 | Pro-Arg | 12 | Clostridia (83.3%) |
| 275.201 | 4.42 | 9-OxoOTrE | 4 | Clostridia (75%) |
| 279.171 | 3.47 | Leu-Phe | 22 | Clostridia (36.4%) |
| 282.279 | 6.09 | N-Tetracosenoyl-4-sphingenine | 7 | Clostridia (71.4%) |
| 285.083 | 0.40 | Xanthosine | 23 | Bacteroidia (26%);<br>Clostridia (26%) |
| 288.203 | 0.37 | Arg-Ile | 25 | Bacteroidia (32%);<br>Clostridia (28%) |
| 291.268 | 5.54 | cis-11,14-Eicosadienoic acid | 3 | Clostridia (66%) |
| 294.119 | 0.38 | N-Acetylmuramic acid | 5 | Bacteroidia (60%) |
| 297.126 | 3.14 | Phe-Met | 3 | Actinobacteria (33.3%);<br>Clostridia (33.3%);<br>Coriobacteriia (33.3%) |
| 302.205 | 3.05 | Ile-Gly-Ile | 1 | Clostridia (100%) |
| 304.166 | 3.02 | Val-Trp | 23 | Clostridia (52.2%);<br>Mollicutes (17.4%) |
| 309.164 | 0.313 | Fructoselysine | 3 | Bacteroidia (66.7%);<br>Clostridia (33.3%) |
| 318.167 | 2.27 | Ile-Trp | 4 | Bacteroidia (50%);<br>Clostridia (50%) |
| 323.273 | 6.84 | Lithocholic acid | 14 | Clostridia (57.1%) |
| 336.192 | 3.22 | Leucine Enkephalin | 4 | Bacteroidia (50%);<br>Clostridia (50%) |
| 338.342 | 9.19 | 13-Docosenamide, (Z)- | 4 | Bacteroidia (50%);<br>Clostridia (25%);<br>Spirochaetes (25%) |
| 359.266 | 0.60 | Ile-Val-Lys | 2 | Clostridia (100%) |
| 369.352 | 10.51 | Cholesterol | 2 | Bacteroidia (50%);<br>Erysipelotrichi (50%) |
| 405.264 | 4.82 | (R)-4-<br>((3R,5S,8R,9S,10S,13R,14S,17R)-<br>3-hydroxy-10,13-dimethyl-7,12-dioxohexadecahydro-1H-cyclopenta[a]phenanthren-17-yl)pentanoic acid | 2 | Bacteroidia (50%);<br>Erysipelotrichi (50%) |
| 439.359 | 7.62 | Oleanolic acid | 12 | Bacteroidia (33.3%) |
| 471.347 | 6.92 | Enoxolone | 23 | Clostridia (39.1%) |
| 585.272 | 8.90 | Bilirubin | 41 | Bacteroidia (29.3%);<br>Clostridia (26.8%);<br>Mollicutes (12.2%) |
| 591.318 | 4.08 | Urobilin | 14 | Bacteroidia (42.9%);<br>Clostridia (28.6%) |
| 595.349 | 4.05 | Stercobilin | 5 | Clostridia (80%) |
| 839.565 | 5.12 | Cholic acid | 15 | Clostridia (40%);<br>Mollicutes (26.7%) |

Our novel data thus represent a core human fecal metabolome from populations of diverse behaviors and lifestyles, yet we do not presume to have captured the range of diversity of industrial lifestyles or age groups seen in international metabolome initiatives. To broaden our analysis, we co-analyzed our data with a total of 1,286 samples from ten public fecal metabolome datasets^47, 49, 60–64^ (Supplementary Table 4), using the Re-Analysis of Data User Interface (ReDU)^39^. These datasets contained samples from male and female children and adults. Eight of the datasets consisted of samples collected from the United States, one contained samples from Venezuela, and one dataset did not report samples’ geographic origin. Furthermore, the datasets included different MS platforms and different metabolite extraction methods, enabling us to assess the commonality of these metabolites across experimental methods. Indeed, every annotated core metabolite (Supplementary Table 2) was detected in this co-analysis, but only 31% were identified in every selected dataset. Such shared annotated molecules include palmitelaidic acid, urobilin, lithocholic acid, and cholesterol. Furthermore, we also examined the human fecal metabolome database (HFMDB)^65^, which contains 6,810 metabolites identified across multiple datasets, for our annotated core metabolite features. 65% of our annotated core metabolite features were present in the HFMDB (Supplementary Table 2); examples of identified metabolites also found in the HFMDB include palmitoleic acid, hypoxanthine, and xanthosine. However, it should be noted that the HFMDB is comprised of data derived from various instrumental, analytical, and processing methods^65^. The absence of some of our core metabolites from the HFMDB can be attributed to these methodological differences.

While we were able to reveal the core human fecal metabolome, only 6.1% of our complete dataset had putative compound-level annotations (level 2 according to the metabolomics standards initiative^66^). Fifteen of these were validated using standards, enabling level 1 confidence^66^ (Supplementary Figure 4). 28.8% of the dataset had annotations based only on chemical class (level 3 of the metabolomics standards initiative^66^). This underscores the need for further annotation of human fecal metabolites, especially from human populations traditionally underrepresented in metabolomic databases. Lastly, it is important to note that samples used for this study were collected at different times and subjected to varying preservation treatments and lengths. However, our samples clustered based on industrialization score rather than storage conditions or geographic origin, indicating that any confounding influence from preservation was overshadowed by the effect of industrialization. Full data are freely available on the GNPS^42^ and ReDU^39^ “living data” infrastructure (see Data Availability statement below) so they can be of use to other researchers and annotations can continue to expand.

Overall, we demonstrate how industrialization profoundly shapes human biology regardless of age, sex, or geographic origin, highlighting the importance of further exploring the biological consequences of industrialization. We also highlight strong commonalities in the fecal metabolome across these distinct populations, representing a core human fecal metabolome of both endogenous and exogenous metabolites. Based on our definition, these chemical components are core to human groups or populations, but not necessarily found in every human individual or LC-MS analysis, given differences in metabolite extraction or instrumental conditions between studies. Further studies focused on untargeted analyses of a spectrum of industrial and non-industrial populations, including past and present humans, can help elucidate the core human fecal metabolome’s ubiquity, its relationship with the gut microbiome, and how processes such as industrialization drive human evolution.

## Materials and Methods

### Project Design

Fecal samples from six human populations were analyzed, representing ranges of industrialization. Populations were assigned industrialization scores to reflect varying degrees of industrialization, based on diet, access to pharmacies and public markets/stores, housing structure, and population density. Score values are: one—highly industrial urban population; two —industrialized rural population; three—a rural community with some industrialization; four— isolated rural community with little to no industrialization. The study populations include: Norman, Oklahoma, USA, a standard Western industrialization population located in the Oklahoma City metropolitan area; Guayabo, Peru, a large rural town influenced by industrialization; Tambo de Mora district, Peru, a large rural district influenced by industrialization; Boulkiemdé province, Burkina Faso, with some industrialization influence; Tunapuco, a traditional rural community located in the Andean Highlands with minimal industrialization influence; and the Matses, an isolated traditional hunter-gatherer community from the Peruvian Amazon (Figure 1; Table 1; Supplementary Table 1). All populations contained both males and females of varying age ranges.

### Populations

Fecal samples from Norman, Oklahoma, USA, were analyzed for this project (*n*=18), representing western industrial lifestyles and diets. Norman residents live in the Oklahoma City metropolitan area, exemplifying a highly industrialized environment. Self-reported diets generally consisted of regular dairy consumption plus processed and/or prepackaged foods like canned vegetables. Due to the strongly industrialized setting and diet, this population received an industrialization score of one.

We also selected fecal samples from the Guayabo (*n*=13) and Tambo de Mora (*n*=17) populations, which practice similar lifestyles. These populations exhibit rural lifestyles and diets but are still strongly influenced by industrialization. Both communities have regular access to public markets and pharmacies and live in densely packed areas. Their diets are generally reliant on foods obtained from these markets, as well as local produce and livestock. While the Guayabo diet commonly consists of maize with some meat and dairy consumption, the Tambo de Mora population relies more on fish, due to their proximity with the Peruvian coastline. Because the Guayabo and Tambo de Mora communities exhibit some characteristics of non-industrial and industrial lifestyles, these populations received an industrialization score of two.

The Boulkiemdé (*n=*11) and Tunapuco (*n=*30) communities represent the next degree of industrialization in our sampled populations. Although these populations are from Africa and South America, respectively, they practice similar traditional non-industrial, rural lifestyles and share some features of industrialized populations such as access to public markets. The Boulkiemdé samples were collected from the Boulkiemdé province of Burkina Faso. This Burkinabé community practices an agricultural lifestyle, usually growing their own crops, raising livestock, and with infrequent dairy consumption. Meanwhile, the Tunapuco population have similar traditional agricultural lifestyles, relying on local produce and livestock. Residing in the Peruvian Andes highlands, the Tunapuco people have diets largely consisting of root and stem tubers, bread, and rice. The Tunapuco people occasionally consume animal proteins and dairy products. Additionally, Tunapuco residents have access to lowland markets, which offer other dietary sources like fruit. Since both the Boulkiemdé and Tunapuco communities sampled for this project lived in largely rural yet partly industrial environments, these populations had an industrialization score of three.

Our last sampled population is the Matses (*n*=16). The Matses people practice traditional hunter-gatherer lifestyles, making them unique for this study. Their diet is based heavily on tubers, plantains, fish, and game meat. Dairy and processed foods are very rarely consumed by the Matses community. Due to their location in the Amazonian regions of Peru and unique lifestyles, the Matses are almost completely isolated from external sociocultural and economic influences like industrialization, so they received an industrialization score of four.

### Sample Collection

Fecal material was deposited into polypropylene containers and then put in ice. Samples were kept in ice while in the field until arriving at research facilities equipped with freezers. The Norman samples were kept in ice after collection and frozen at the laboratory within 24 hours.

The Peruvian samples were secured similarly to the Norman samples. After collection, samples were stored on ice for four days until arriving at Lima, Peru. Samples were frozen and sent to the laboratory in Norman, Oklahoma.

The Norman, Tunapuco, and Matses samples had previously been aliquoted and underwent 16S rRNA gene sequencing for an earlier study^20^, using the MoBio PowerSoil DNA Isolation Kit protocol (full details can be found in the original article^20^). The raw fecal samples were otherwise kept frozen at −80 °C until use for this project.

Boulkiemdé samples were collected similarly to Norman and Peruvian samples. After collection, Boulkiemdé samples were frozen at −20 °C and kept frozen overnight. Samples were thawed the following evening to extract DNA, refrozen at −20 °C, and kept frozen until shipped to the laboratory in Norman, Oklahoma. Upon arrival, 2 g of fecal material was extracted from each sample for anaerobic culturing. Following this 2 g aliquoting, samples were frozen at −80 °C until use for this project.

### Ethics Approval and Informed Consent

Ethical protocols for community engagement and sample collection were developed through collaboration with representatives and authorities from each sampled region and in accordance with institutional regulations. All Peruvian samples were obtained through community engagement with local and national authorities and informed consent with consultation from the Center for Intercultural Health of the Peruvian Institute of Health and Peruvian National Institute of Health ethics committee. This project was reviewed and approved by the research ethics committee of the Instituto Nacional de Salud del Peru (Projects PP-059-11, OEE-036-16).

Human fecal samples were collected with informed consent from resident volunteers in central Burkina Faso under the ethics review committee of Centre MURAZ, a national health research institute in Burkina Faso (IRB ID No. 31/2016/CE-CM). OU IRB deemed this project consistent with US policy 45 CRF 46.101(b) exempt category 4 (OU IRB 6976).

### LC-MS/MS Fecal Sample Preparation

The sample preparation protocol used for this project was adapted from a global metabolite extraction protocol with proven success^67^. Samples were thawed and 500 μl of chilled LC-MS grade water (Fisher Scientific) was added to 50 mg of fecal material. Next, a Tissuelyzer homogenized samples at 25 Hz for three minutes. Following homogenization, chilled LC-grade methanol (Fisher Scientific) spiked with 4 μM sulfachloropyridazine as the internal standard (IS) was added, bringing the total concentration to 50% methanol. The TissueLyzer homogenized samples again at 25 Hz for three minutes, followed by overnight incubation at 4 °C. The next day, samples were centrifuged at 16,000 x g at 4 °C for ten minutes. Aqueous supernatant was then removed and dried using a SpeedVac vacuum concentrator. Dried extracts were frozen at −80 °C until the day of MS analysis. Immediately prior to MS analysis, extracts were resuspended in 150 μl chilled LC-MS methanol:water (1:1) spiked with 1 μg/ml sulfadimethoxine as a second IS. After resuspension, samples were diluted to a 1:10 ratio. Diluted samples were sonicated using a Fisher Scientific Ultrasonic Cleaning Bath at maximum power for ten minutes.

Supernatants were spun briefly to remove any particulates, then loaded into a 96-well plate for MS analysis. One well contained only 150 μl of the resuspension solution to serve as a negative control.

### LC-MS/MS Analysis

LC was performed on a ThermoFisher Scientific Vanquish Flex Binary LC System with a Kinetex C18 core-shell column (50 x 2.1 mm, 1.7 μM particle size, 100 Å pore size). LC column was kept at 40 °C and the sample compartment was held at 10 °C. The LC System was coupled to a ThermoFisher Scientific Q Exactive Plus Hybrid Quadrupole-Orbitrap Mass Spectrometer for MS/MS analysis. For the LC mobile phase, Solvent A was LC-MS grade water (Fisher Scientific) with 0.1% formic acid and Solvent B was LC-MS grade acetonitrile (Fisher Scientific) with 0.1% formic acid. Elution gradient started at 5% Solvent B for one minute, increased to 100% Solvent B until minute nine, held at 100% Solvent B for two minutes, dropped to 5% Solvent B over 30 seconds, and 5% Solvent B for one minute as re-equilibration. Samples were injected in random order with an injection volume of 5 μl. After elution, electrospray ionization was conducted with spray voltage of 3.8 kV, auxiliary gas flow rate of 10 L/min, auxiliary gas temperature at 350 °C, sheath gas flow rate at 35 L/min, and sweep gas flow at 0 L/min. Capillary temperature was 320 °C and S-lens RF was 50 V.

MS1 scan range was 100-1,500 *m/z*, MS1 resolution was set to 35,000 and MS1 AGC target to 1e6. MS1 data were obtained in positive mode and MS2 data were obtained using data-dependent acquisition. In each cycle, 5 MS/MS scans of the most abundant ion were recorded. Both MS1 and MS2 injection times were set at 100 ms. MS2 resolutions were set to 17,500, MS2 AGC target was set to 5e5, and the inclusion window to 2 *m/z*. MS/MS was conducted at an apex trigger of 2-8 seconds and an exclusion window of 10 seconds. MS/MS collision energy gradually increased from 20-40%.

Authentic standards also underwent LC-MS/MS analysis to validate metabolite annotations. A total of 15 standards were purchased from AA Blocks (hyocholic acid, 13-docosenamide), AvaChem (lenticin), Biosynth (bilirubin, N-acetylmuramic acid, fructosyl-L-lysine), BLD Pharm (N-palmitoylglycine, trans-ferulic acid), ChemScene (leucine enkephalin), LGC Standards (L-saccharopine), Sigma-Aldrich (L-abrine, N-acetyl L-phenylalanine, enoxolone, octadecanamide, lithocholic acid, paraxanthine), and VWR (nicotinamide N-oxide). Each pure standard was diluted to 100 μM, 50 μM, 10 μM, 5 μM, and 1 μM concentrations to maximize standard detection. All standards (and their five dilutions) were analyzed according to the same LC-MS/MS parameters as the original samples. Additionally, fecal extracts with the highest abundance for each standard were re-analyzed as part of the same LC-MS/MS batch to ensure standard peaks were present in samples and to prevent confounding from retention time shifts caused by the gap between initial data acquisition and annotation validation.

### Data Analysis and Processing

MSConvert v3.0.19014^68^ converted raw data files to mzXML format in preparation for data processing via feature-based molecular networking (FBMN)^69^. MZmine v2.33^70^ was used to identify MS features for all samples (Supplementary Table 5). After feature filtering, only features with abundance three times greater than the abundance of blanks were retained in these analyses. Total ion current (TIC) normalization was conducted through R programming language v3.5.3^71^ in Jupyter Notebook^72^. FBMN and library spectral database searches were completed using the FBMN workflow on Global Natural Products Social Molecular Networking (GNPS)^42^. FBMN GNPS parameters for MS/MS analysis were as follows: precursor and fragment ion mass tolerance: 0.02 Da, minimum cosine score for networking and library matches: 0.7, minimum number of matched MS2 fragment ions for networking and library matches: 4, network topK: 50, maximum connected component size: 100, maximum shift between precursors: 500 Da, analog search: enabled, maximum analog mass difference: 100 Da, precursor window filtering: enabled, 50 Da peak window filtering: enabled, normalization per file: row sum normalization. Results were analyzed by visually evaluating mirror plot similarity, cosine score, and match likelihood. Molecular networking results were exported to Cytoscape v3.7.1^73^ to visualize and analyze networks. Predicted ClassyFire^74^ classifications for shared metabolites were derived using the MolNetEnhancer^75^ workflow in GNPS. In addition, select annotations were confirmed using authentic standards (Supplementary Figure 4).

MS filtering was performed in MZmine^70^. Three separate filtering workflows were done: 6 minimum peaks in a row (half the number of samples in a single population), 52 minimum peaks in a row (half our total samples), and 105 minimum peaks in a row (all samples). After each filtering step, gap-filling was performed using the previous parameters. For the six-sample filtering, additional processing was done in R^71^ to remove any features that were not found in at least six samples from each population. The resulting files were also analyzed in GNPS as described above.

For 16S rRNA gene sequencing data, we used AdapterRemoval v2^76^ to filter out sequences < 90 bp in length. QIIME1^77^ was used to perform closed-reference OTU picking using the EzTaxon database^78^ as a reference. For OTU picking, the maximum number of database hits per sequence was eight and the maximum number of rejects for a new OTU was 12. After creating biom files, each sample file was rarefied to a depth of 10,000. Generated taxa summaries were limited to genus-level identifications. Only taxa with >0.5% relative frequency were included for correlation analyses.

### Correlation and Statistical Analyses

Nonparametric Spearman correlation coefficients were calculated with false discovery rate correction (Supplementary File 1) using metabolite and OTU abundances per sample and per industrialization group. Normalized metabolite feature abundances were summed across each industrialization group using the feature tables derived from R processing via JupyterNotebook. OTU abundances per industrialization group were calculated by determining the relative abundance for each sample and summing the sample abundances according to industrialization score group assignments. Correlation networks with relative MS1 and OTU abundances were visualized using Cytoscape v3.7.1^73^. Weak correlations (correlation coefficient between −0.3 and 0.3) were excluded from subsequent analyses.

Principal coordinate analysis (PCoA) plots were created using Canberra distance metrics from Quantitative Insights Into Microbial Ecology 2 (QIIME2)^79^ and visualized using Emperor^80^. PERMANOVA via QIIME2 assessed statistical significance for beta diversity measures. Kruskal-Wallis p-values were calculated in R^71^ through Jupyter Notebook^72^. Boxplots (Figure 1c-h, Supplementary Figures 1-2) were also generated using R^71^ in JupyterNotebook^72^. For these boxplots, the center line represents the median, the upper and lower box lines reflect upper and lower quartiles, whiskers reflect the interquartile range multiplied by one-and-a-half, and outliers are dots. R packages ggplot2^81^ and rworldmap^82^ were used to create Figures 1a, 1c-h,. External visualization tools in GNPS v23^42^ were used to create UpSet plots^83^. R package effectsize^84^ provided p-values for ANOVA effect size.

To identify metabolite features unique to specific populations or lifestyles, a random forest machine learning algorithm from the R package “randomForest” was used in Jupyter Notebook^85^. The number of trees increased gradually from five until reaching a plateau from out-of-bag error at 200 trees. SIRIUS v4.4.26^86^ with ClassyFire^74^ classification and CANOPUS^87^ compound prediction were used to provide class-level annotations for features identified by random forest.

### Data Availability

LC-MS/MS data was uploaded to MassIVE (accession number: MSV000084794). GNPS FBMN jobs are available at https://gnps.ucsd.edu/ProteoSAFe/status.jsp?task=196ab44c10e44c1d898f15e7c046a591 (v21, original analysis) and https://gnps.ucsd.edu/ProteoSAFe/status.jsp?task=505b8b39810c48eb9f9b65fee7c6bc7b (v23, primarily used throughout data analysis). FBMN jobs for filtered data are available at: https://gnps.ucsd.edu/ProteoSAFe/status.jsp?task=db26beb51aff418585e6ad0b92f522b7 (six-sample per population filter, gap-filling), https://gnps.ucsd.edu/ProteoSAFe/status.jsp?task=4693e01a2af740ceb39bfb19720e798d (six-sample per population filter, no gap-filling), https://gnps.ucsd.edu/ProteoSAFe/status.jsp?task=220d1afd0a564ec1818601d3d928d27a (half-sample filter, gap-filling), https://gnps.ucsd.edu/ProteoSAFe/status.jsp?task=d9686d483e5b496299a02750d6a3ec23 (half-sample filter, no gap-filling), https://gnps.ucsd.edu/ProteoSAFe/status.jsp?task=45150c751a8e42eea51f3ea4936aee95 (all-sample filter, gap-filling), and https://gnps.ucsd.edu/ProteoSAFe/status.jsp?task=45150c751a8e42eea51f3ea4936aee95 (all-sample filter, no gap-filling). ReDU co-analysis is available at https://gnps.ucsd.edu/ProteoSAFe/status.jsp?task=bd266e2a3eba45fa9a1b3819e809a1b6 (co-analysis with fecal data from different MS instruments). Instructions for recreating data analyses in R are available as JupyterNotebook^72^ links at: https://github.com/jhaffner09/core_metabolome_2021. 16S data was uploaded to the Qiita database (study ID: 13802).

Public ReDU datasets used for co-analyses are available at: MSV000083559^47^ (doi: 10.25345/C5C032; dataset license: CC0 1.0 Universal); MSV000082433^47, 62, 63^ (dataset license: CC0 1.0 Universal); MSV000081351^47^ (dataset license: CC0 1.0 Universal); MSV000083756^64^ (doi: 10.25345/C53S6N; dataset license: CC0 1.0 Universal); MSV000083300^61^ (doi: 10.25345/C56C86; dataset license: CC0 1.0 Universal); MSV000081492^47^ (dataset license: CC0 1.0 Universal); MSV000082629^47^ (dataset license: CC0 1.0 Universal); MSV000082262 (dataset license: CC0 1.0 Universal); MSV000082221^60^ (dataset license: CC0 1.0 Universal); and MSV000082374^49^ (dataset license: CC0 1.0 Universal).

## Supplementary Tables and Figures

**Supplementary Figure 1.**
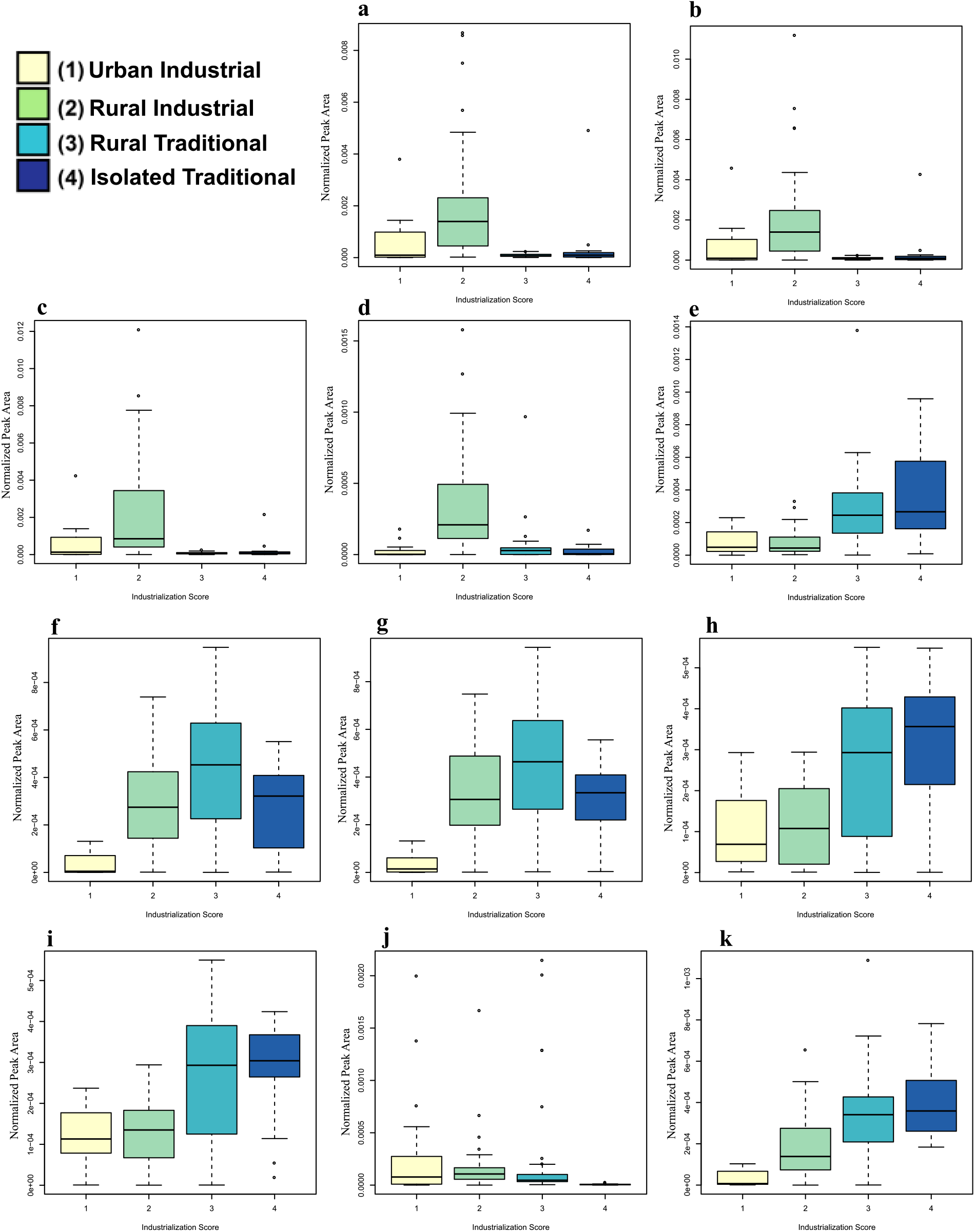
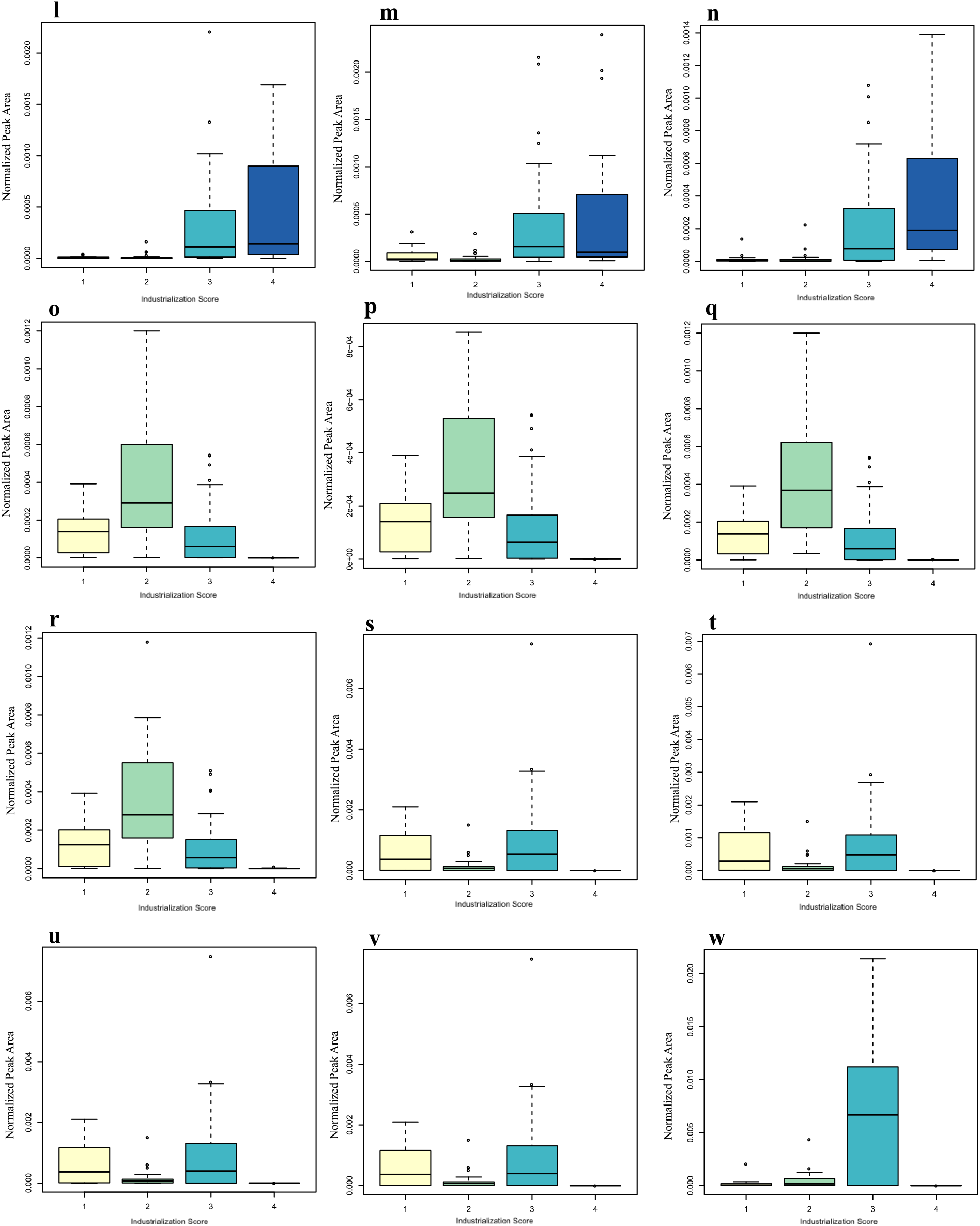
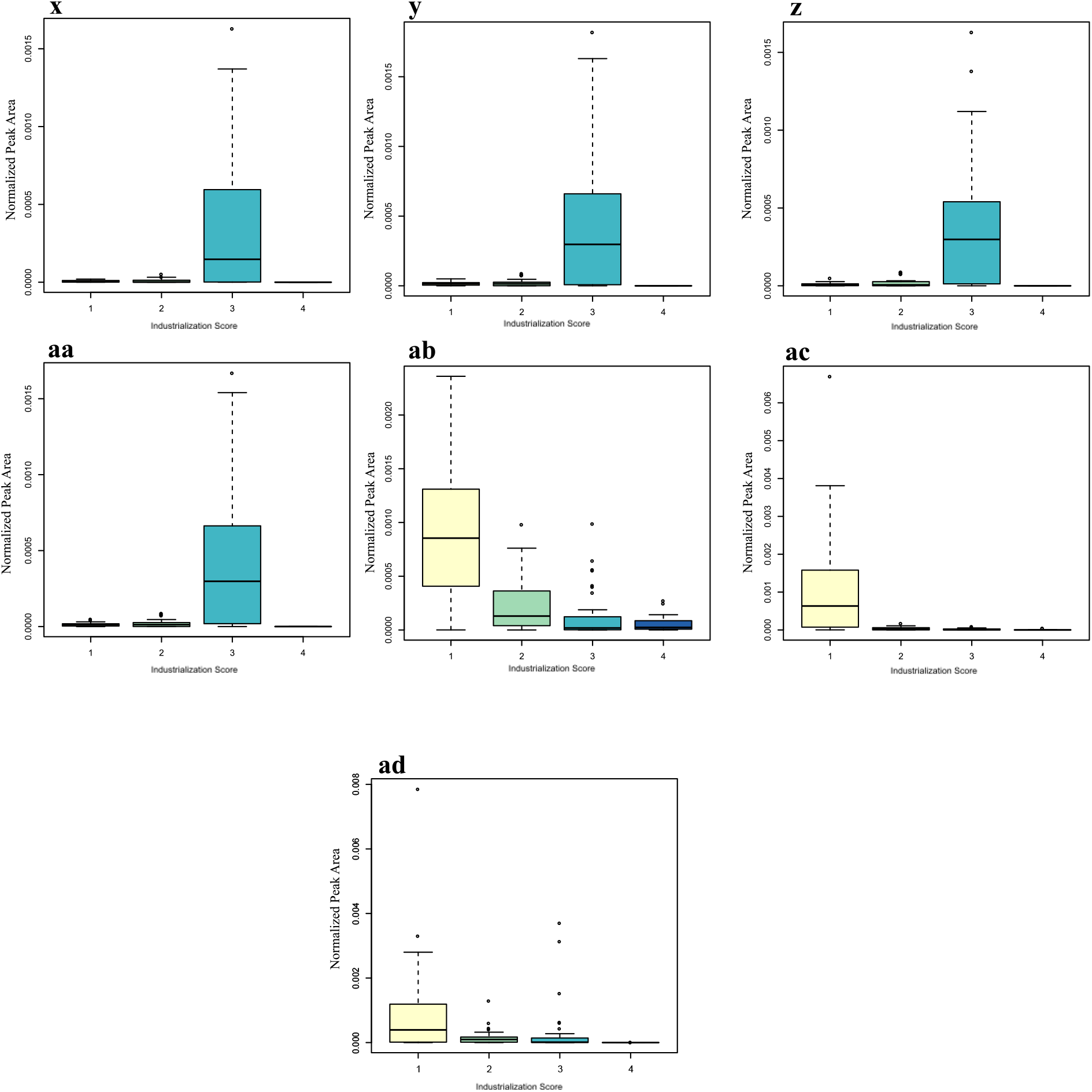
Abundances of top 30 differential metabolite features based on industrialization score identified by RandomForest. Color key applies for figures **a-ad**. **a,** *m/z* 145.134, RT 0.32 min; **b,** *m/z* 145.134, RT 0.32 min; **c,** *m/z* 145.1340, RT 0.36 min; **d**, *m/z* 235.166, RT 0.25 min; **e,** *m/z* 246.186, RT 3.27 min; **f,** 276.108, RT 4.41 min; **g,** *m/z* 276.108, 0.42; **h,** *m/z* 286.176, RT 1.41 min; **i,** *m/z* 286.176, RT 1.677 min; **j,** *m/z* 305.186, RT 3.74 min; **k,** *m/z* 332.074, RT 0.36 min; **l,** *m/z* 363.211, RT 1.02; **m,** *m/z* 363.213, RT 0.87 min; **n,** *m/z* 365.192, RT 0.51 min; **o,** *m/z* 379.295, RT 4.8 min; **p,** *m/z* 379.296, RT 4.82 min; **q,** *m/z* 379.296, RT 4.8 min; **r,** *m/z* 379.297, RT 4.81 min; **s,** *m/z* 398.342, RT 4.76 min; **t,** *m/z* 398.342, RT 4.82 min; **u,** *m/z* 398.342, RT 4.84 min; **v,** *m/z* 398.345, RT 4.81 min; **w,** *m/z* 400.358, RT 4.83 min; **x,** *m/z* 414.335, RT 4.49 min; **y,** *m/z* 414.337, RT 4.43 min; **z,** *m/z* 414.337, RT 4.38 min; **aa,** *m/z* 414.337, RT 4.43 min; **ab,** 591.318, RT 4.52; **ac,** *m/z* 593.333, RT 4.98 min; **ad,** *m/z* 597.37, RT 5.31 min.

**Supplementary Figure 2.**
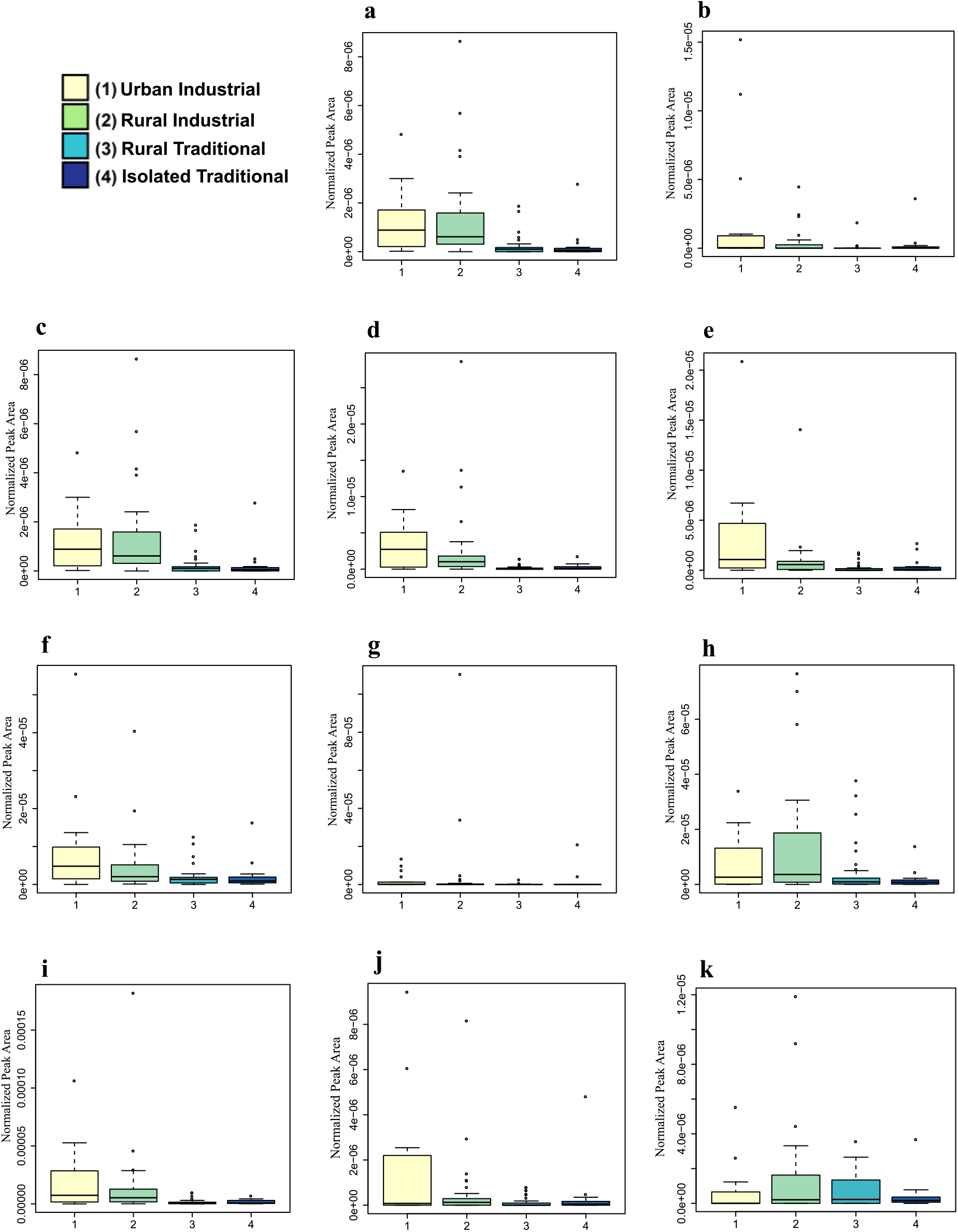
Normalized abundances of amino acid-conjugated bile acids by industrialization score. Same color key from **a** applies to figures **b**-**k. a,** leucocholic acid (Kruskal-Wallis p=1.69e-7); **b,** tyrosocholic acid (Kruskal-Wallis p=7.71e-3; **c,** glutamate-conjugated chenodeoxycholic acid (Kruskal-Wallis p=1.69e-7); **d,** tryptophan-conjugated chenodeoxycholic acid (Kruskal-Wallis p=4.9e-7); **e,** aspartate-conjugated chenodeoxycholic acid (Kruskal-Wallis p=1.13e-5); **f,** histidine-conjugated chenodeoxycholic acid (Kruskal-Wallis p=6.41e-3); **g,** histidine-conjugated cholic acid (Kruskal-Wallis p=0.04); **h,** leucine-conjugated chenodeoxycholic acid (Kruskal-Wallis p=0.04); **i,** tyrosine-conjugated deoxycholic acid (Kruskal-Wallis p=1.61e-5). **j**, aspartate-conjugated cholic acid (Kruskal-Wallis p=0.05); **k**, threonine-conjugated chenodeoxycholic acid (Kruskal-Wallis p=0.4

**Supplementary Figure 3.**
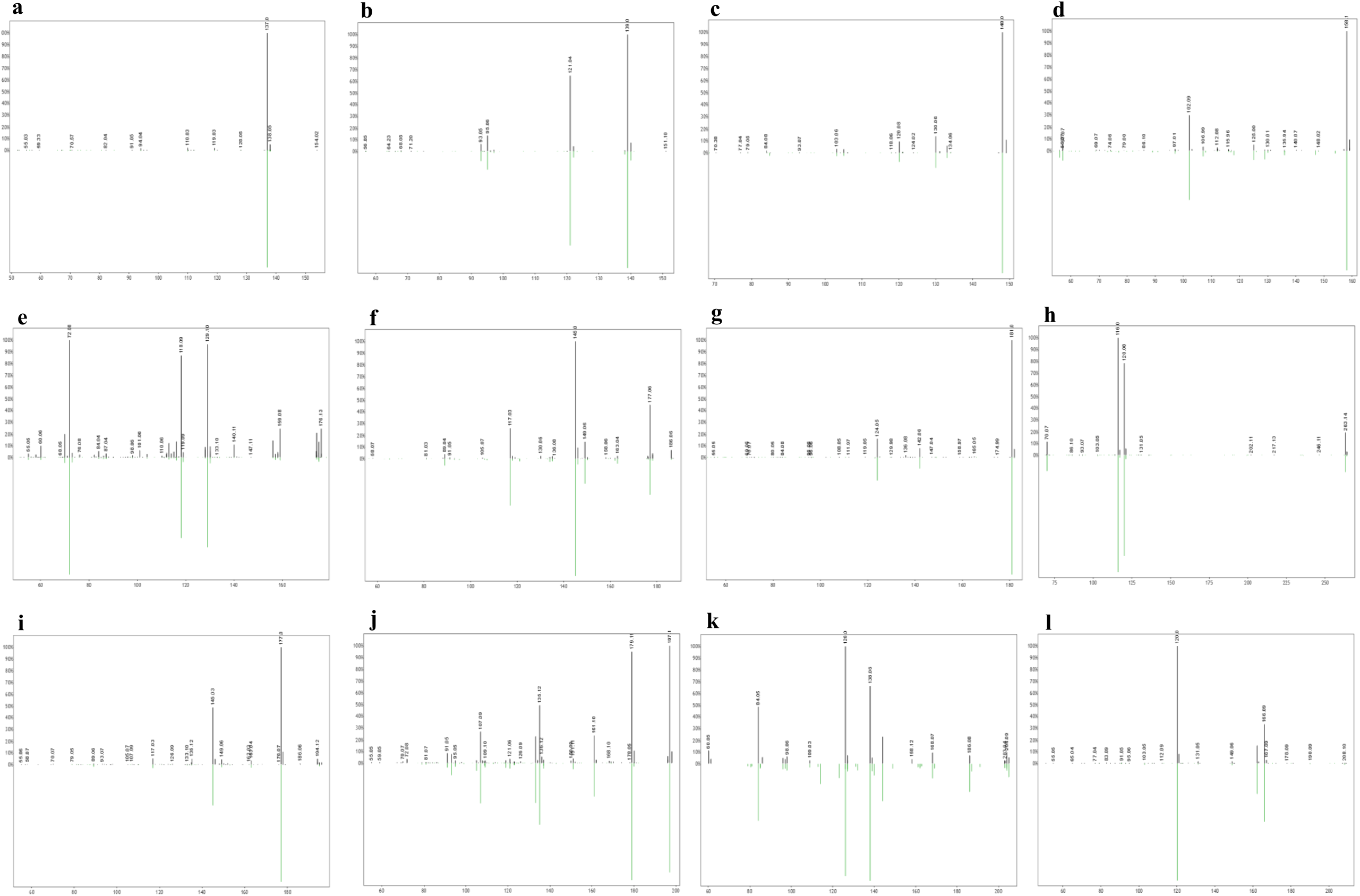
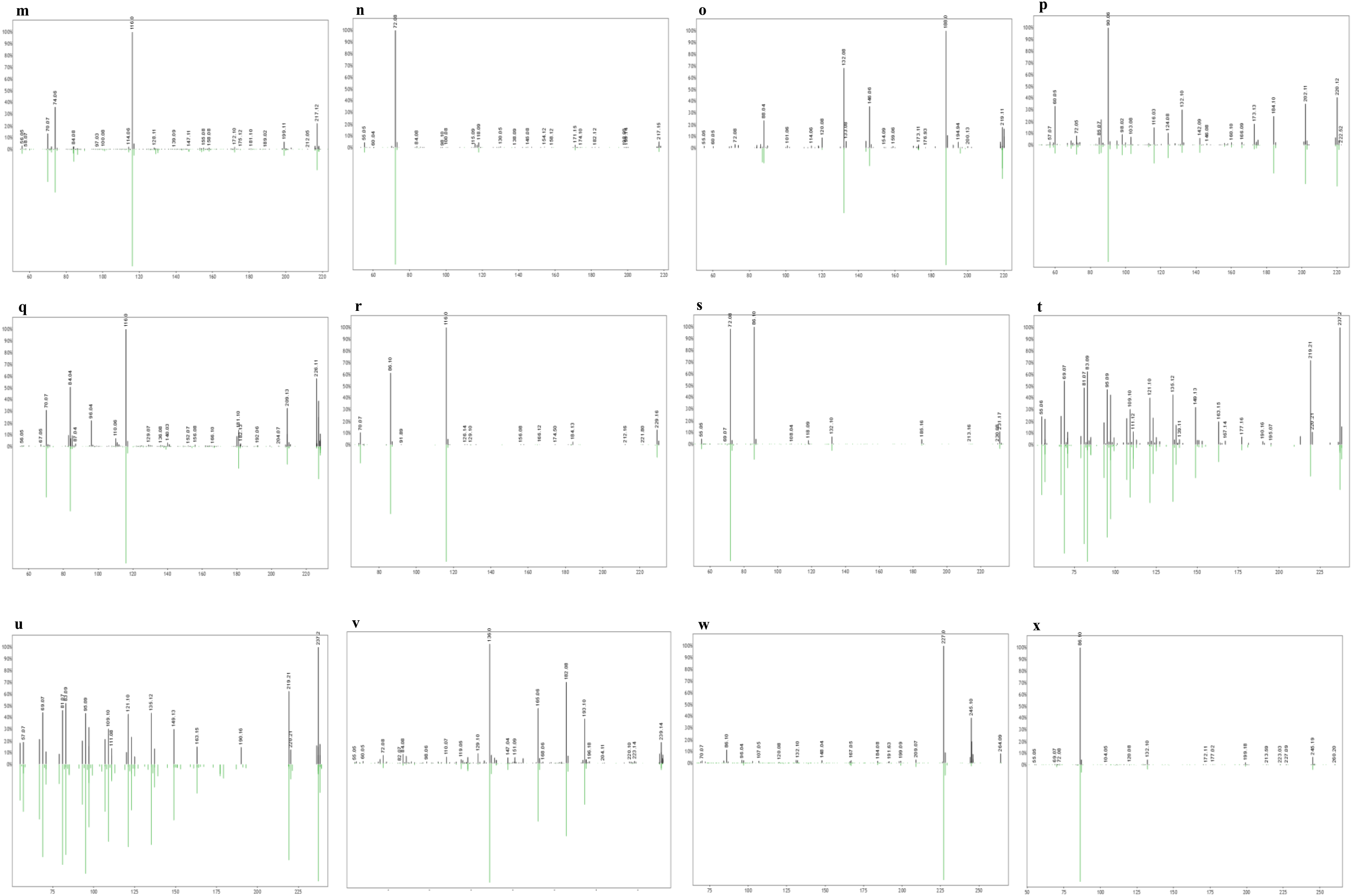
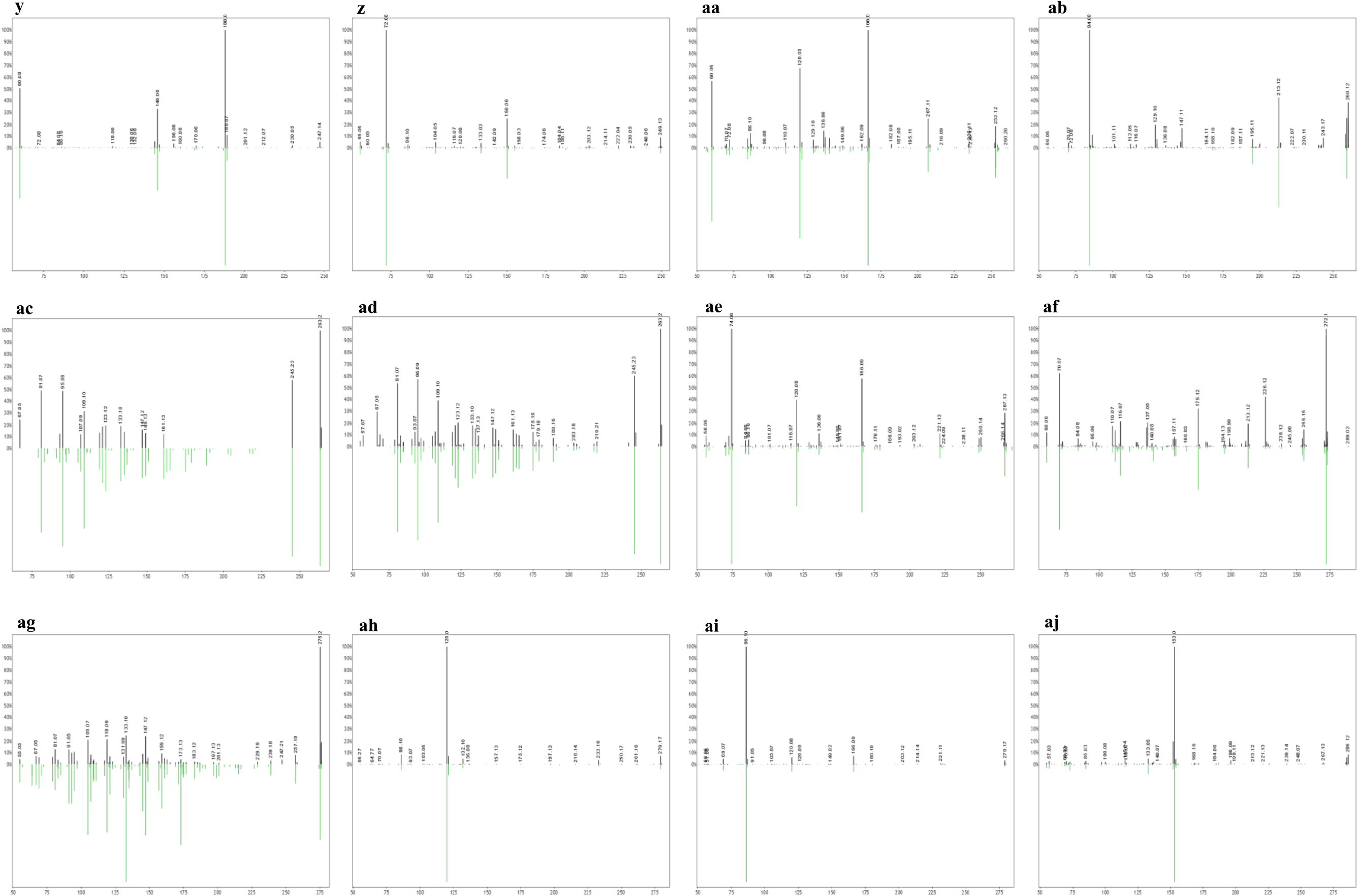
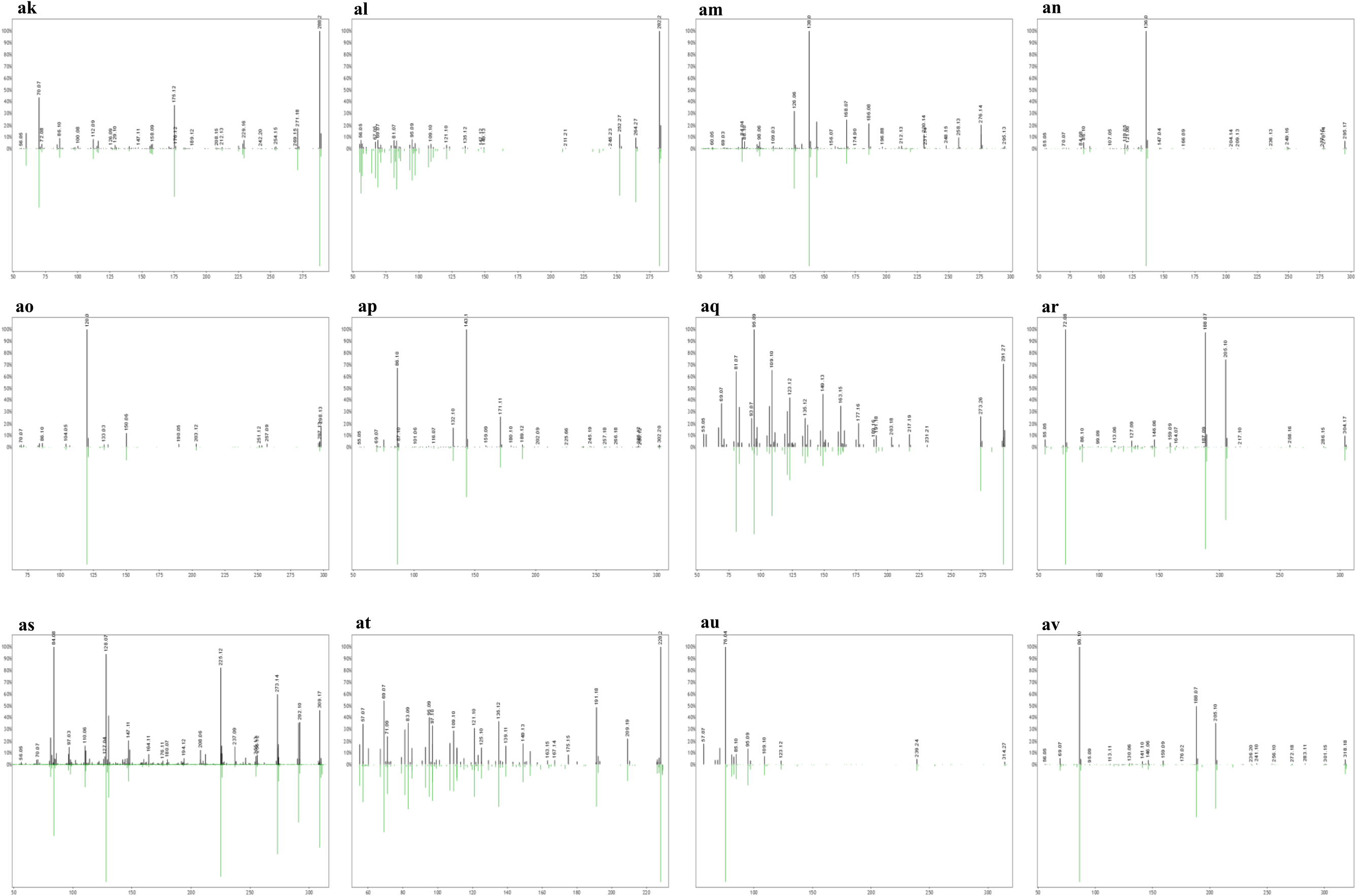
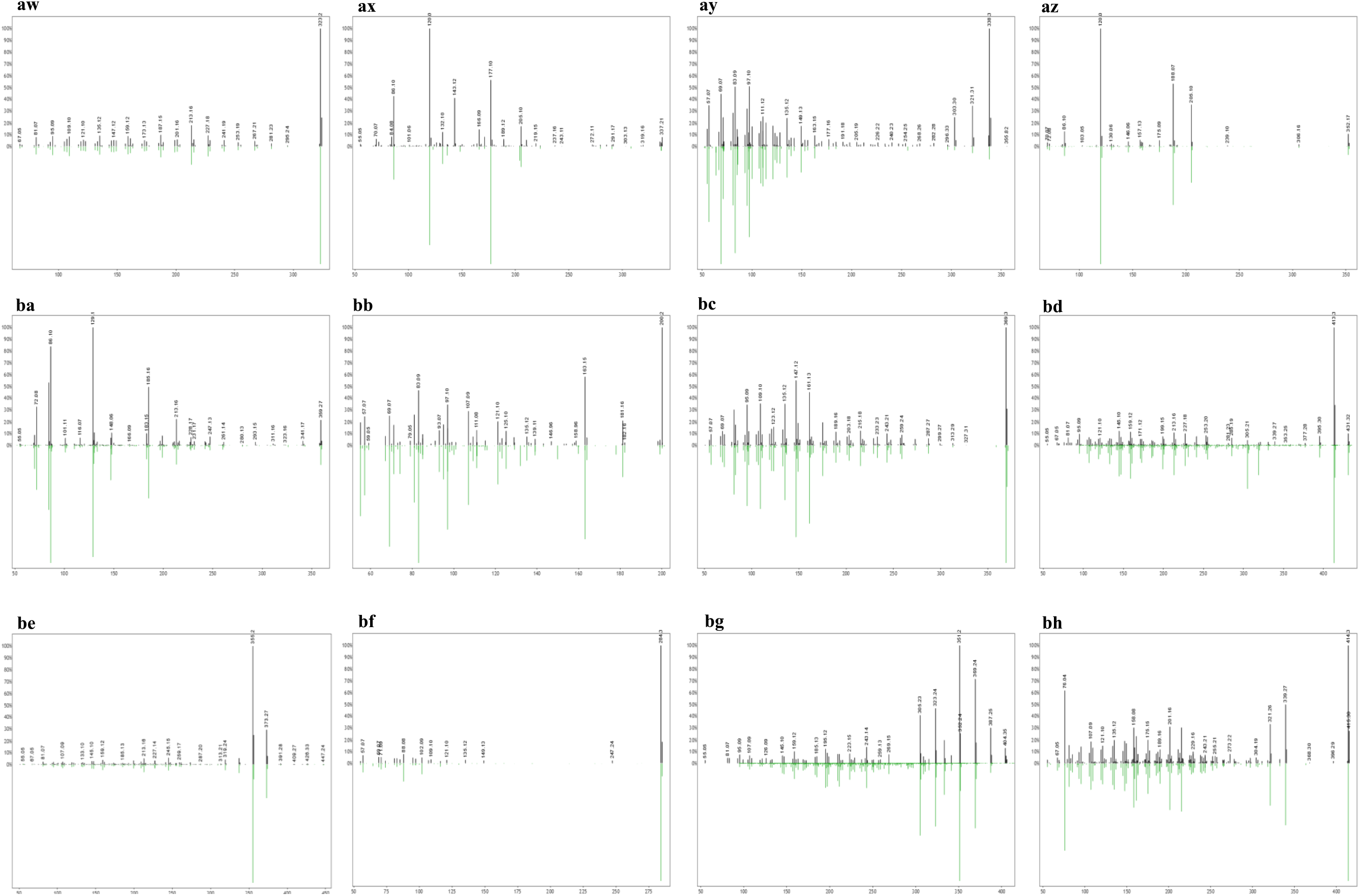
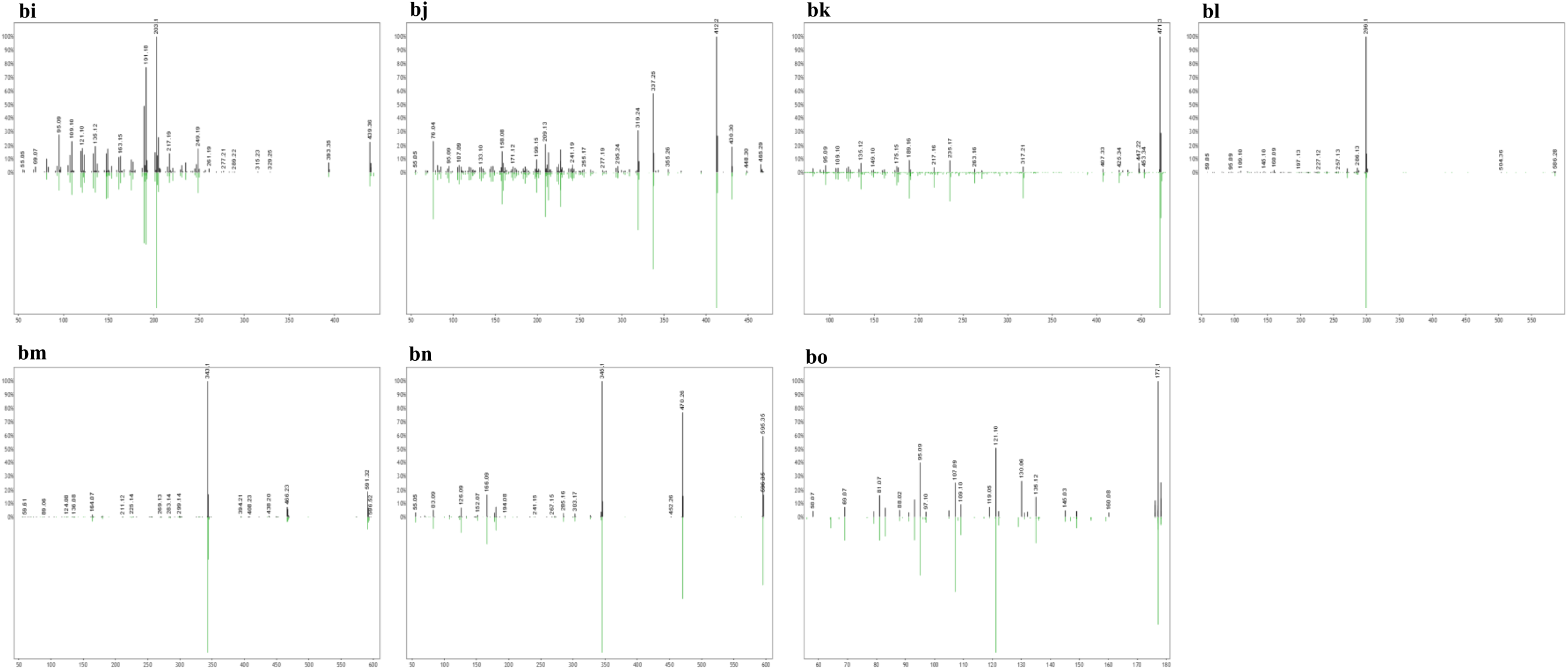
Mirror plots of core human fecal metabolites. Top black bars represent sample peaks for the respective metabolite, while bottom green bars represent library reference peaks. **a**, Hypoxanthine; **b**, Nicotinamide N-oxide; **c**, 3-methyl-2-oxindole (3-Methyloxindole); **d**, Hyocholic acid; **e**, Gly-Val (Glycylvaline); **f**, 3-Hydroxy-4-methoxycinnamic acid (Isoferulic acid); **g**, Paraxanthine; **h**, Phe-Pro (Phenylalanylproline); **i**, trans-Ferulic acid; **j**, Loliolide; **k**, N-Acetyl-D-mannosamine; **l**, N-acetyl-L-Phenylalanine; **m**, Thr-Pro (Threonylproline); **n**, Val-Val (Valylvaline); **o**, Abrine; **p**, Pantothenic acid; **q**, PyroGlu-Pro (Pyroglutamylproline); **r**, Ile-Pro (Isoleucylproline); **s**, Val-Ile (Valylisoleucine); **t**, cis-9-Hexadecenoic acid (Palmitoleic acid); **u**, Palmitelaidic acid; **v**, Gly-Tyr (Glycyltyrosine); **w**, Biotin; **x**, Leu-Leu (Leucylleucine); **y**, Lenticin; **z**, Val-Met (Valylmethionine); **aa**, Ser-Phe (Serylphenylalanine); **ab**, L-Saccharopine; **ac**, Conjugated linoleic Acid (10E,12Z); **ad**, Conjugated linoleic acid (9E,11E); **ae**, Thr-Phe (Threonylphenylalanine); **af**, Pro-Arg (Prolylarginine); **ag**, 9-OxoOTrE; **ah**, Phe-Leu (Phenylalanylleucine); **ai**, Leu-Phe (Leucylphenylalanine); **aj**, Xanthosine; **ak**, Arg-Ile (Arginylisoleucine); **al**, N-Tetracosenoyl-4-sphingenine; **am**, N-Acetylmuramic Acid; **an**, Tyr-Leu (Tyrosylleucine); **ao**, Phe-Met (Phenylalanylmethionine); **ap**, Ile-Gly-Ile (Isoleucylglycylisoleucine); **aq**, cis-11,14-Eicosadienoic acid; **ar**, Val-Trp (Valyltryptophan); **as**, Fructoselysine; **at**, Myristoleic acid; **au**, N-Palmitoylglycine; **av**, Ile-Trp (Iseoleucyltryptophan); **aw**, Lithocholic acid; **ax**, Leucine enkephalin; **ay**, 13-Docosenamide, (Z)-(Erucamide); **az**, Phe-Trp (Phenylalanyltryptophan); **ba**, Ile-Val-Lys (Isoleucylvalyllysine); **bb**, 3-Hydroxydodecanoic acid; **bc**, Cholesrol; **bd**, 6R)-2-(hydroxymethyl)-6-((3R,5R,7R,8R,9S,10S,12S,13R,14S,17R)-3,7,12-trihydroxy-10,13-dimethylhexadecahydro-1H-cyclopenta[a]phenanthren-17-yl)heptanoic acid; **be**, Cholic acid; **bf**, Octadecanamide; **bg**, (R)-4-((3R,5S,8R,9S,10S,13R,14S,17R)-3-hydroxy-10,13-dimethyl-7,12-dioxohexadecahydro-1H-cyclopenta[a]phenanthren-17-yl)pentanoic acid; **bh**, Glycoursodeoxycholic acid; **bi**, Oleanolic acid; **bj**, Glycocholic acid; **bk**, Enoxolone; **bl**, Bilirubin; **bm**, Urobilin; **bn**, Stercobilin; **bo**, 2-Butanone, 4-(2,6,6-trimethyl-2-cyclohexen-1-yl)

**Supplementary Figure 4.**
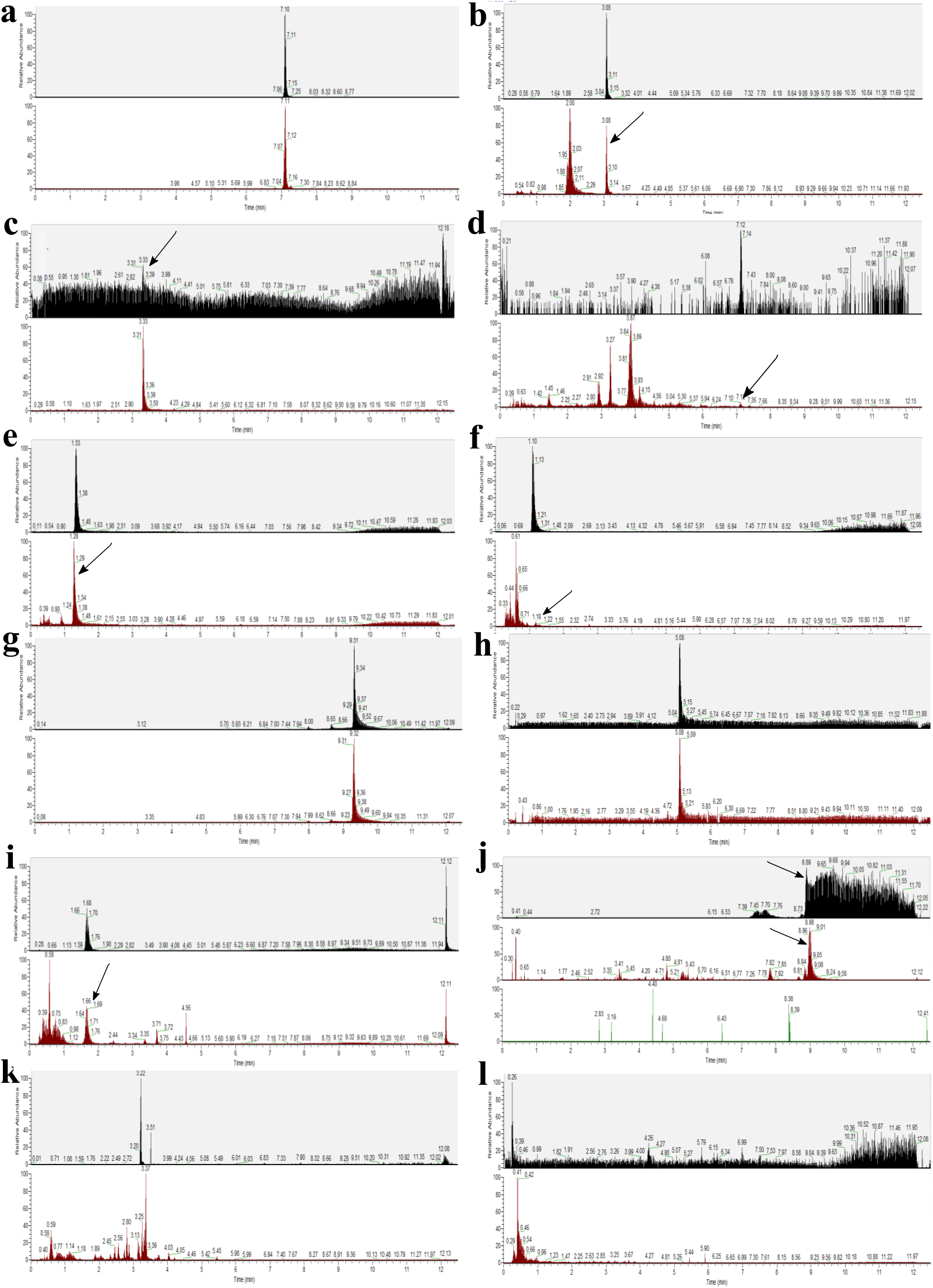
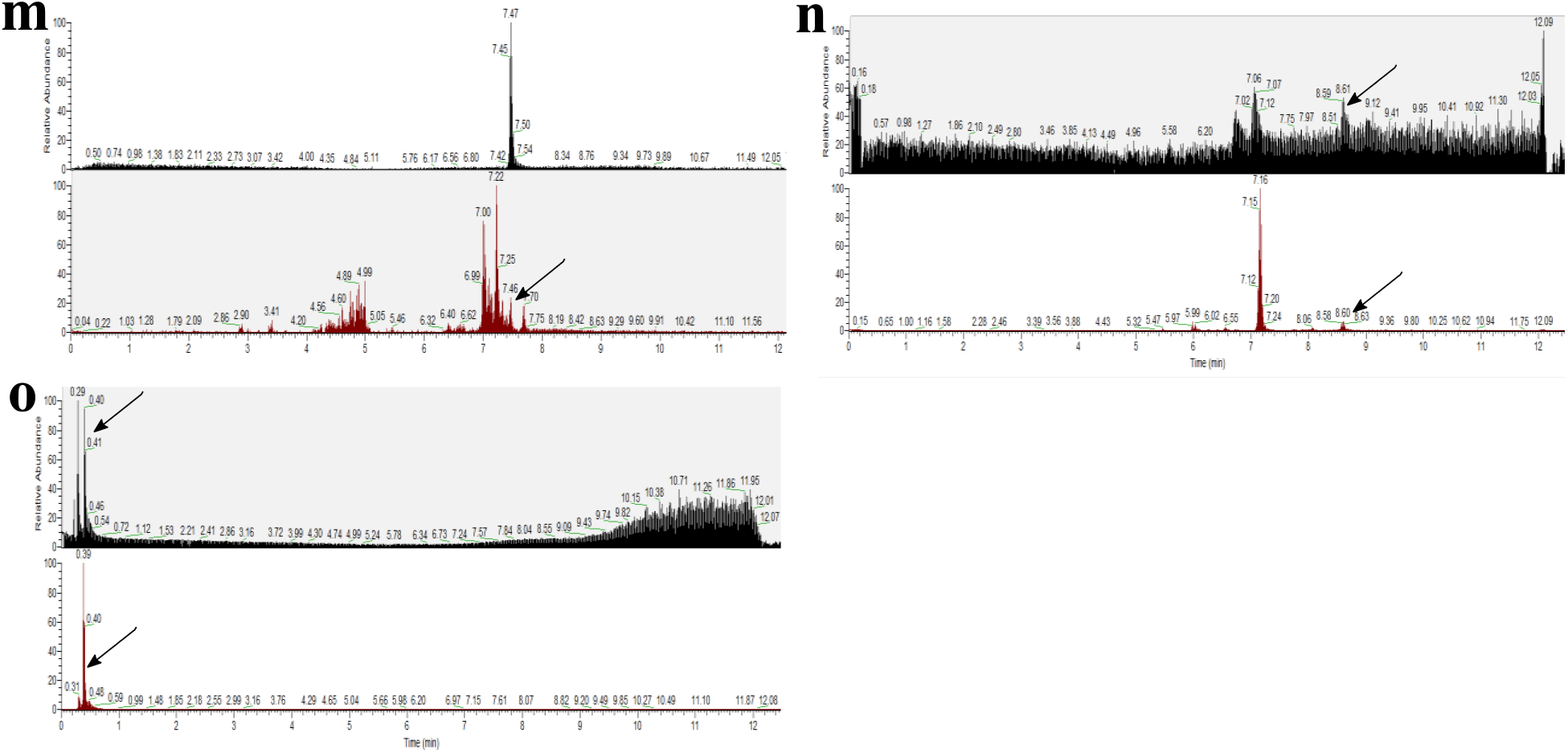
Authentic standards validating sample annotations. Standards are in black, representative samples are in red, blanks (if available) are in green. **a,** Enoxolone; **b,** N-acetyl-L-phenylalanine; **c,** trans-Ferulic acid; **d,** Lithocholic acid; **e,** Paraxanthine; **f,** L-Abrine; **g,** 13-Docosenamide, (Z)-; **h,** Hyocholic acid; **i,** Lenticin; **j,** Bilirubin. Included a blank (filtered to same *m/z*) to demonstrate standard has a distinct peak not seen in negative controls; **k,** Leucine enkephalin; **l,** L-Saccharopine; **m,** N-palmitoylglycine; **n,** Octadecanamide; **o,** Nicotinamide N-oxide.

**Supplementary Table 1.** Metadata for individual samples.

| SampleID | Population | Industrialization Score | Sex | Age |
| --- | --- | --- | --- | --- |
| NO01 | Norman | 1 | M | 23 |
| NO02 | Norman | 1 | F | 37 |
| NO03 | Norman | 1 | M | 40 |
| NO04 | Norman | 1 | M | 26 |
| NO05 | Norman | 1 | M | 28 |
| NO06 | Norman | 1 | M | 28 |
| NO07 | Norman | 1 | F | 32 |
| NO08 | Norman | 1 | F | 32 |
| NO09 | Norman | 1 | F | 34 |
| NO10 | Norman | 1 | M | 41 |
| NO11 | Norman | 1 | M | 26 |
| NO12 | Norman | 1 | F | 27 |
| NO13 | Norman | 1 | M | 35 |
| NO19 | Norman | 1 | F | 32 |
| NO20 | Norman | 1 | M | 26 |
| NO21 | Norman | 1 | M | 23 |
| NO22 | Norman | 1 | M | 26 |
| NO23 | Norman | 1 | F | 26 |
| GU1 | Guayabo | 2 | F | 52 |
| GU2 | Guayabo | 2 | F | 19 |
| GU4 | Guayabo | 2 | F | 40 |
| GU6 | Guayabo | 2 | NA | 6 |
| GU7 | Guayabo | 2 | NA | 9 |
| GU10 | Guayabo | 2 | F | 58 |
| GU11 | Guayabo | 2 | NA | 7 |
| GU12 | Guayabo | 2 | NA | 16 |
| GU13 | Guayabo | 2 | F | 41 |
| GU16 | Guayabo | 2 | NA | 4 |
| GU17 | Guayabo | 2 | F | 51 |
| GU19 | Guayabo | 2 | F | 24 |
| GU20 | Guayabo | 2 | F | 63 |
| 1TM | Tambo de Mora | 2 | F | 61 |
| 2TM | Tambo de Mora | 2 | F | 40 |
| 3TM | Tambo de Mora | 2 | NA | 5 |
| 4TM | Tambo de Mora | 2 | F | 40 |
| 6TM | Tambo de Mora | 2 | F | 31 |
| 10TM | Tambo de Mora | 2 | NA | 8 |
| 11TM | Tambo de Mora | 2 | M | 39 |
| 14TM | Tambo de Mora | 2 | F | 77 |
| 16TM | Tambo de Mora | 2 | F | 38 |
| 17TM | Tambo de Mora | 2 | NA | 7 |
| 18TM | Tambo de Mora | 2 | NA | 13 |
| 23TM | Tambo de Mora | 2 | NA | 1 |
| 24TM | Tambo de Mora | 2 | NA | 1 |
| 26TM | Tambo de Mora | 2 | F | 36 |
| 27TM | Tambo de Mora | 2 | NA | 13 |
| 28TM | Tambo de Mora | 2 | NA | NA |
| 31TM | Tambo de Mora | 2 | NA | 28 |
| TM10_01 | Burkina Faso | 3 | M | 55 |
| TM13_01 | Burkina Faso | 3 | M | 53 |
| TM23_02 | Burkina Faso | 3 | F | 51 |
| TM01_01 | Burkina Faso | 3 | M | 55 |
| TM09_02 | Burkina Faso | 3 | F | 32 |
| TM11_04 | Burkina Faso | 3 | F | 40 |
| TM17_02 | Burkina Faso | 3 | F | 35 |
| TM20_03 | Burkina Faso | 3 | F | 37 |
| TM22_03 | Burkina Faso | 3 | M | 29 |
| TM25_03 | Burkina Faso | 3 | F | 38 |
| TM29_01 | Burkina Faso | 3 | M | 73 |
| HCO01 | Tunapuco | 3 | F | 36 |
| HCO03 | Tunapuco | 3 | M | 6 |
| HCO04 | Tunapuco | 3 | M | 4 |
| HCO07 | Tunapuco | 3 | F | 3 |
| HCO09 | Tunapuco | 3 | NA | 13 |
| HCO10 | Tunapuco | 3 | M | 10 |
| HCO11 | Tunapuco | 3 | M | 36 |
| HCO12 | Tunapuco | 3 | F | 35 |
| HCO13 | Tunapuco | 3 | F | 9 |
| HCO14 | Tunapuco | 3 | F | 34 |
| HCO15 | Tunapuco | 3 | F | 63 |
| HCO16 | Tunapuco | 3 | NA | 11 |
| HCO17 | Tunapuco | 3 | F | 7 |
| HCO18 | Tunapuco | 3 | M | 11 |
| HCO21 | Tunapuco | 3 | M | 10 |
| HCO41 | Tunapuco | 3 | NA | 54 |
| HCO53 | Tunapuco | 3 | F | 44 |
| HCO61 | Tunapuco | 3 | F | 20 |
| HCO62 | Tunapuco | 3 | M | NA |
| HCO63 | Tunapuco | 3 | F | 6 |
| HCO64 | Tunapuco | 3 | F | NA |
| HCO65 | Tunapuco | 3 | F | NA |
| HCO66 | Tunapuco | 3 | M | 11 |
| HCO67 | Tunapuco | 3 | F | 26 |
| HCO68 | Tunapuco | 3 | M | 7 |
| HCO69 | Tunapuco | 3 | NA | 9 |
| HCO70 | Tunapuco | 3 | F | 40 |
| HCO72 | Tunapuco | 3 | F | 5 |
| HCO73 | Tunapuco | 3 | F | NA |
| HCO74 | Tunapuco | 3 | F | 36 |
| SM01 | Matses | 4 | M | 30 |
| SM02 | Matses | 4 | F | 25 |
| SM03 | Matses | 4 | M | 10 |
| SM05 | Matses | 4 | M | 1 |
| SM10 | Matses | 4 | F | 6 |
| SM11 | Matses | 4 | F | 4 |
| SM23 | Matses | 4 | M | 7 |
| SM25 | Matses | 4 | F | 2 |
| SM28 | Matses | 4 | F | 52 |
| SM29 | Matses | 4 | F | 50 |
| SM30 | Matses | 4 | M | 4 |
| SM33 | Matses | 4 | F | 5 |
| SM34 | Matses | 4 | M | 4 |
| SM37 | Matses | 4 | M | 12 |
| SM39 | Matses | 4 | F | 40 |
| SM41 | Matses | 4 | M | 6 |

**Supplementary Table 2.**
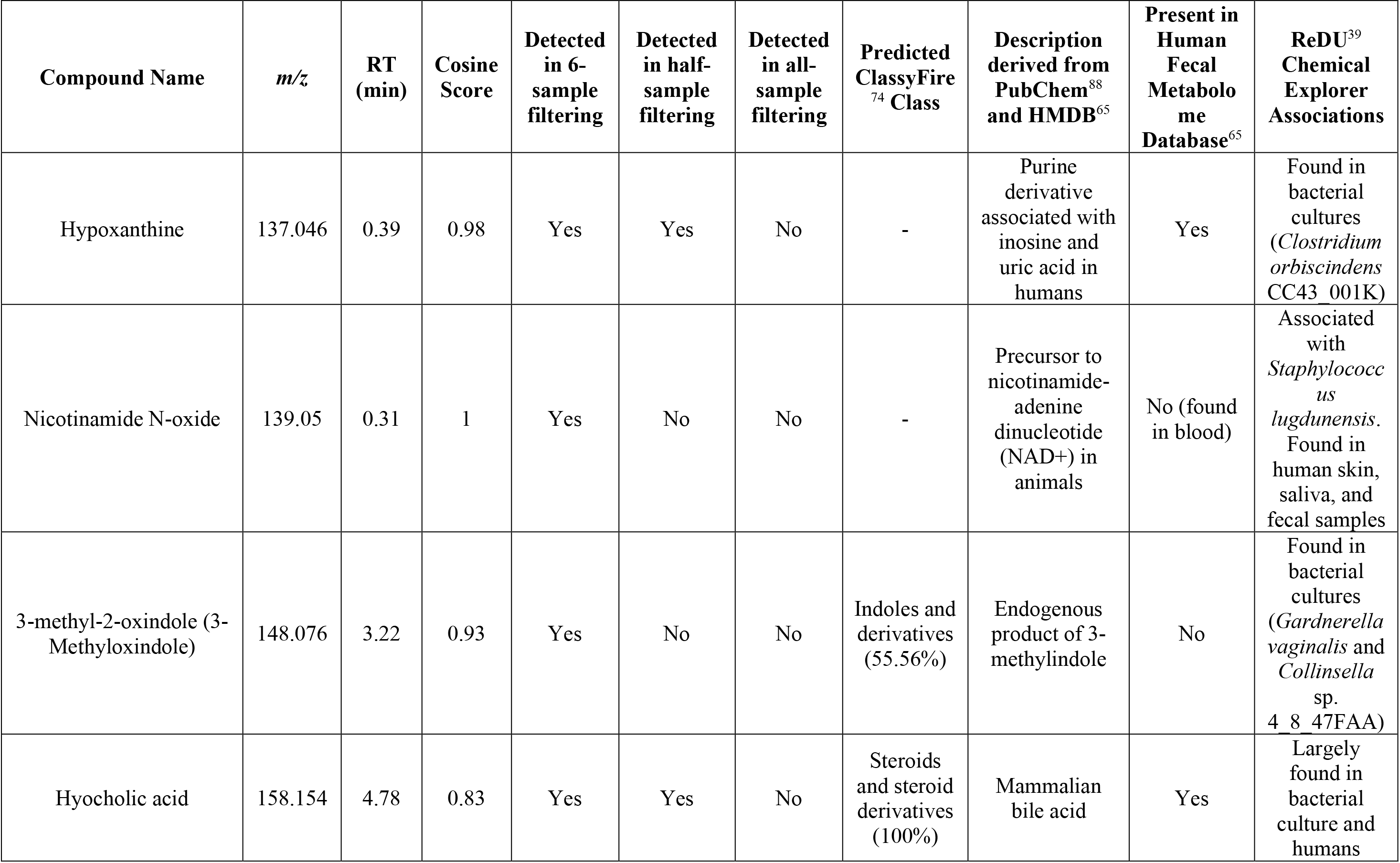
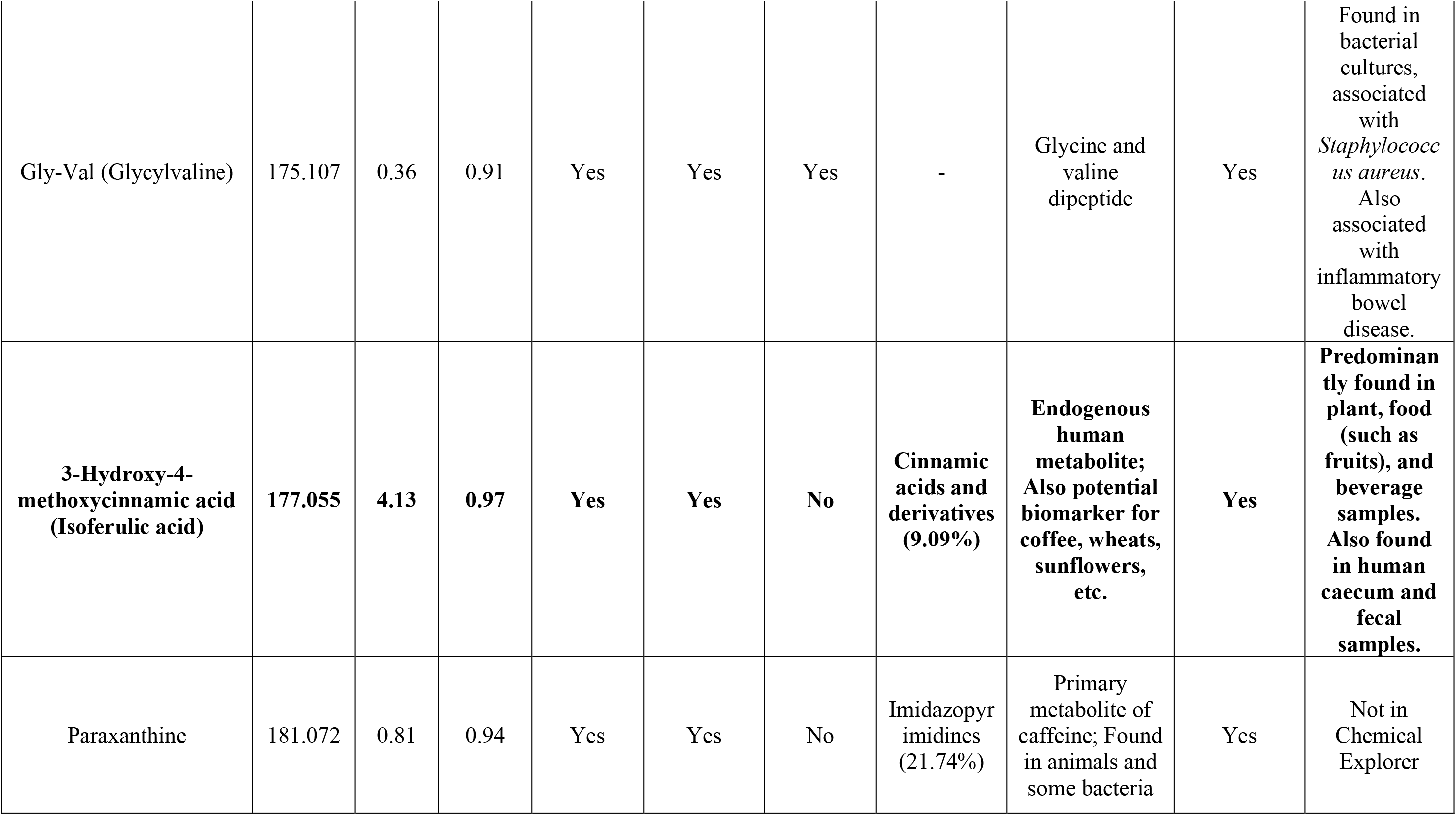
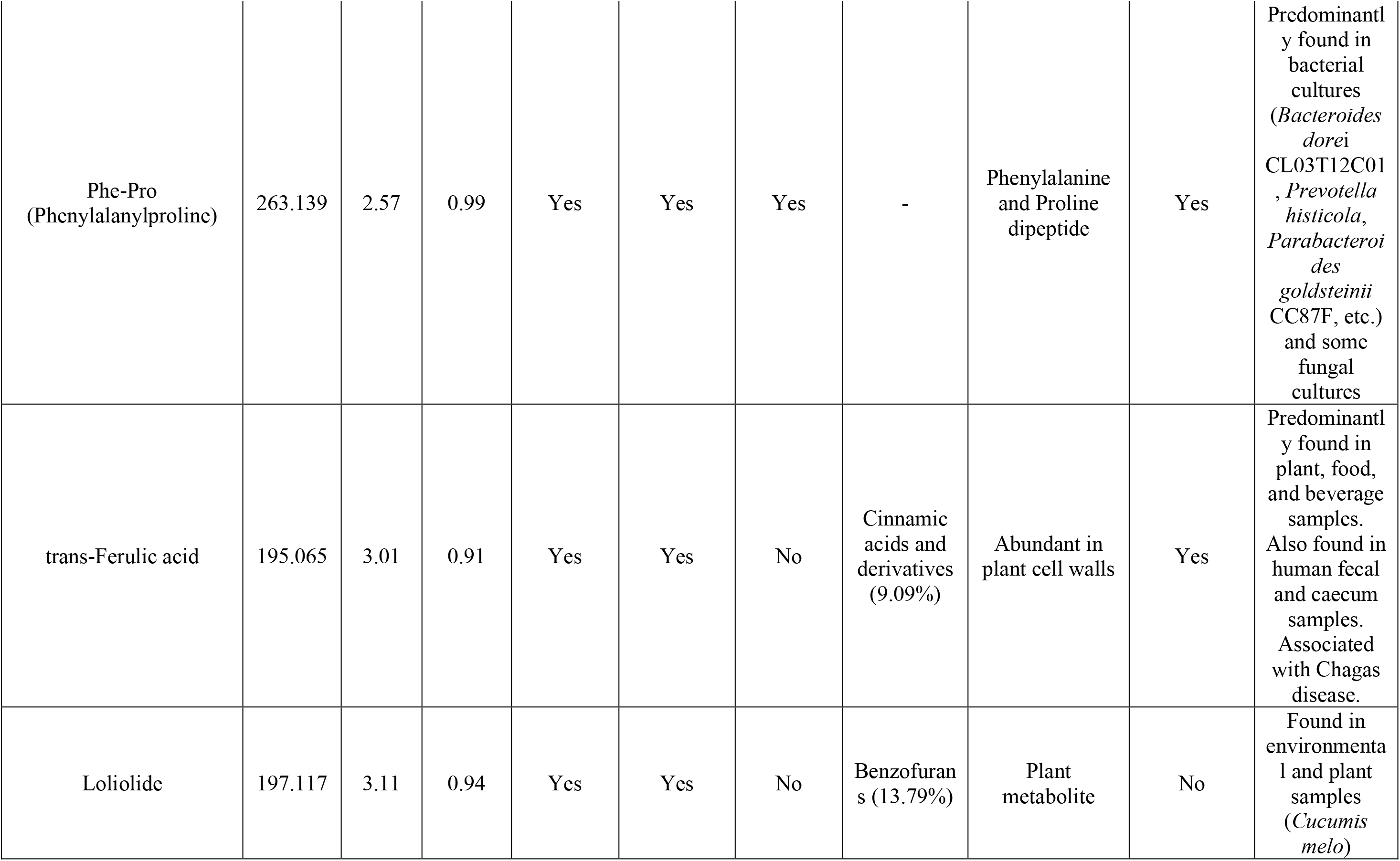
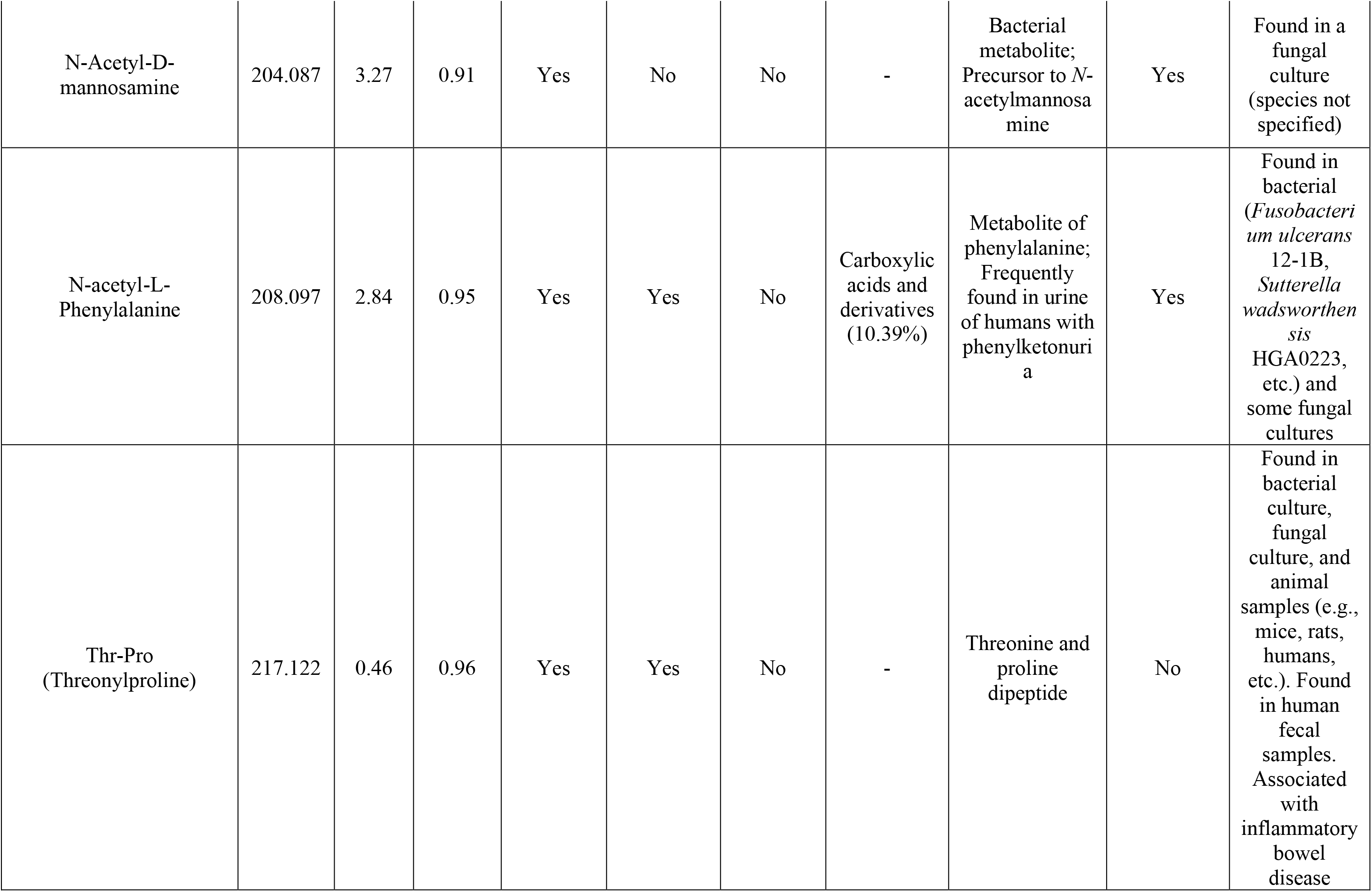
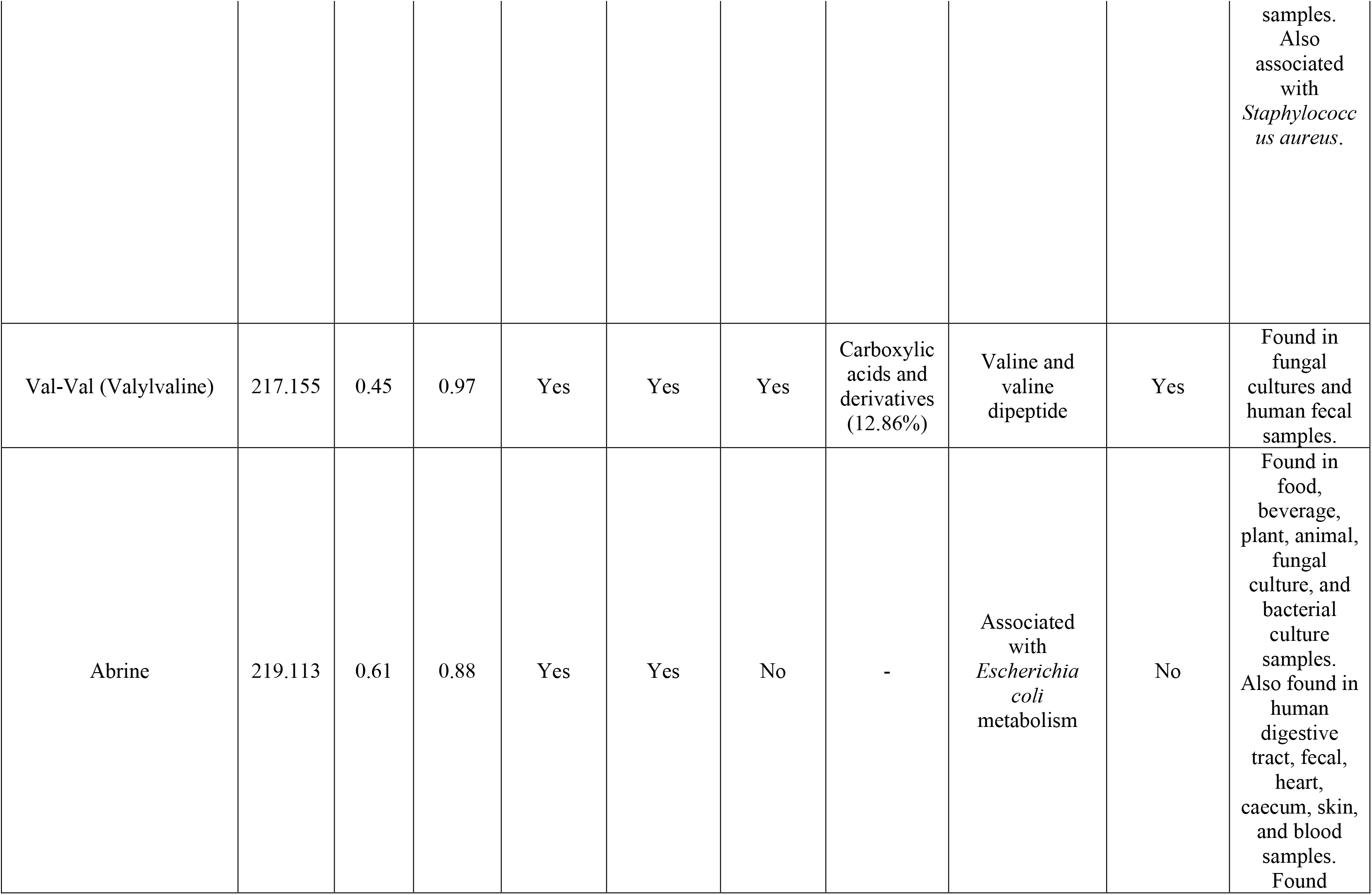
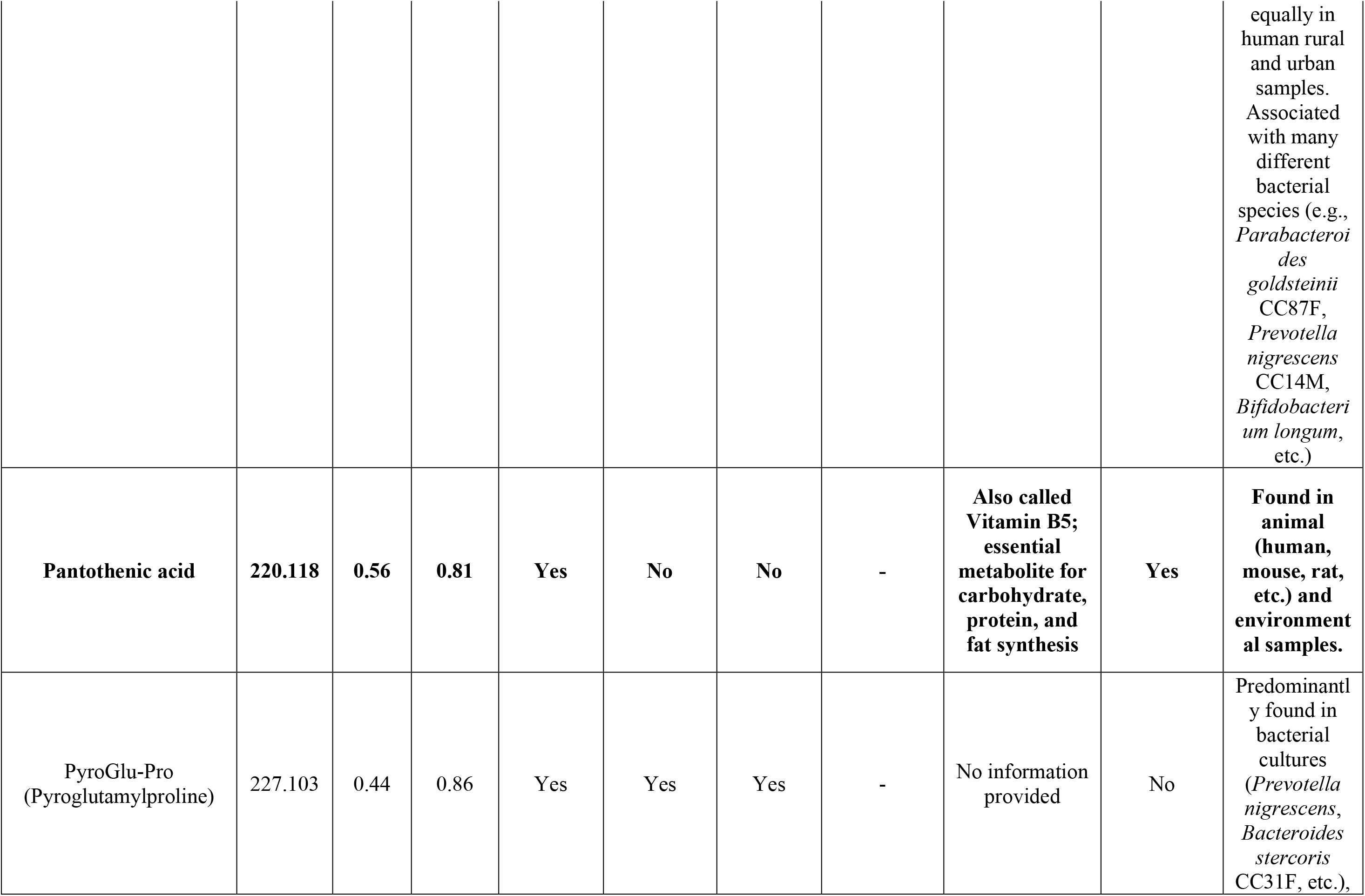
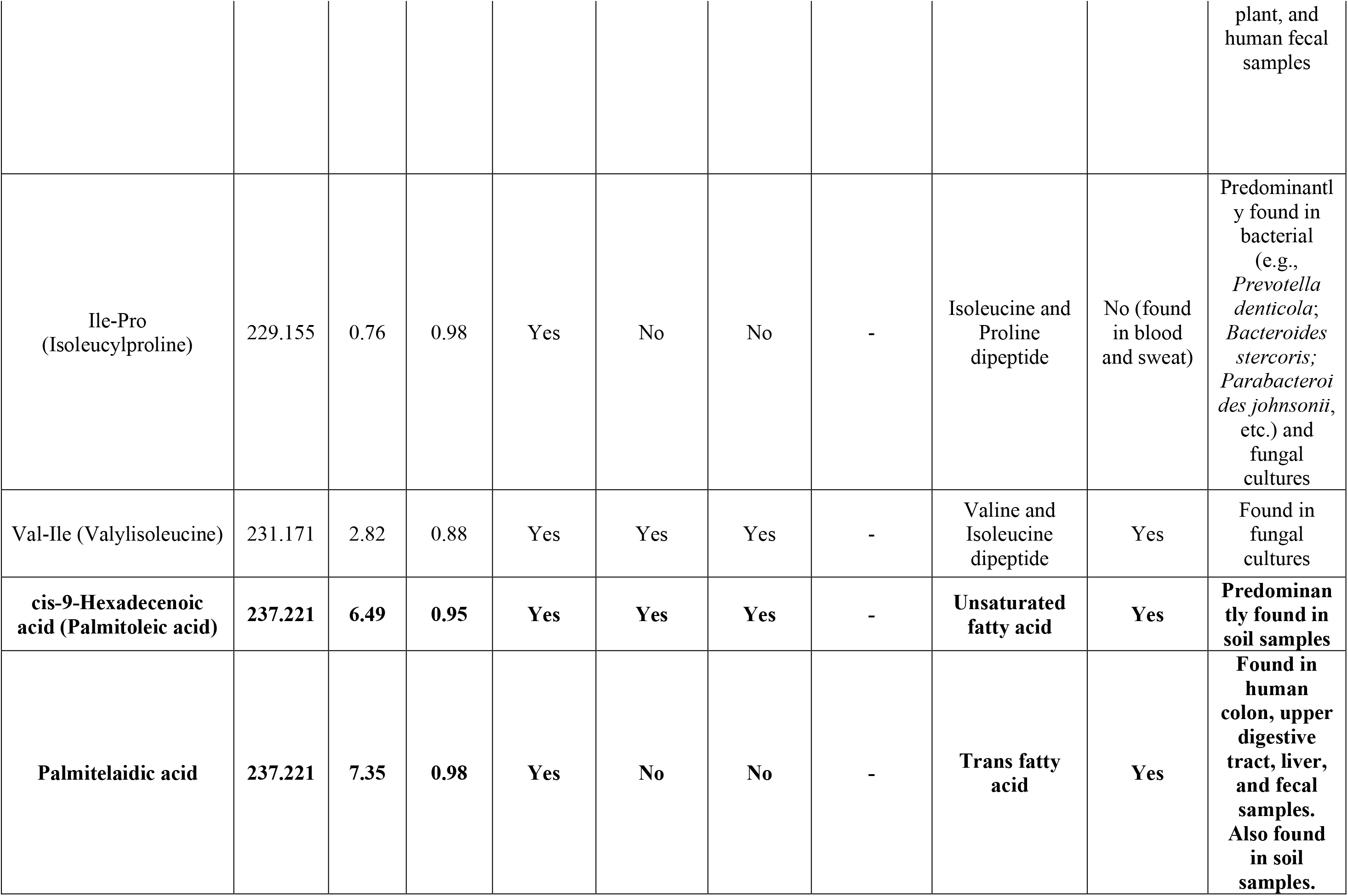
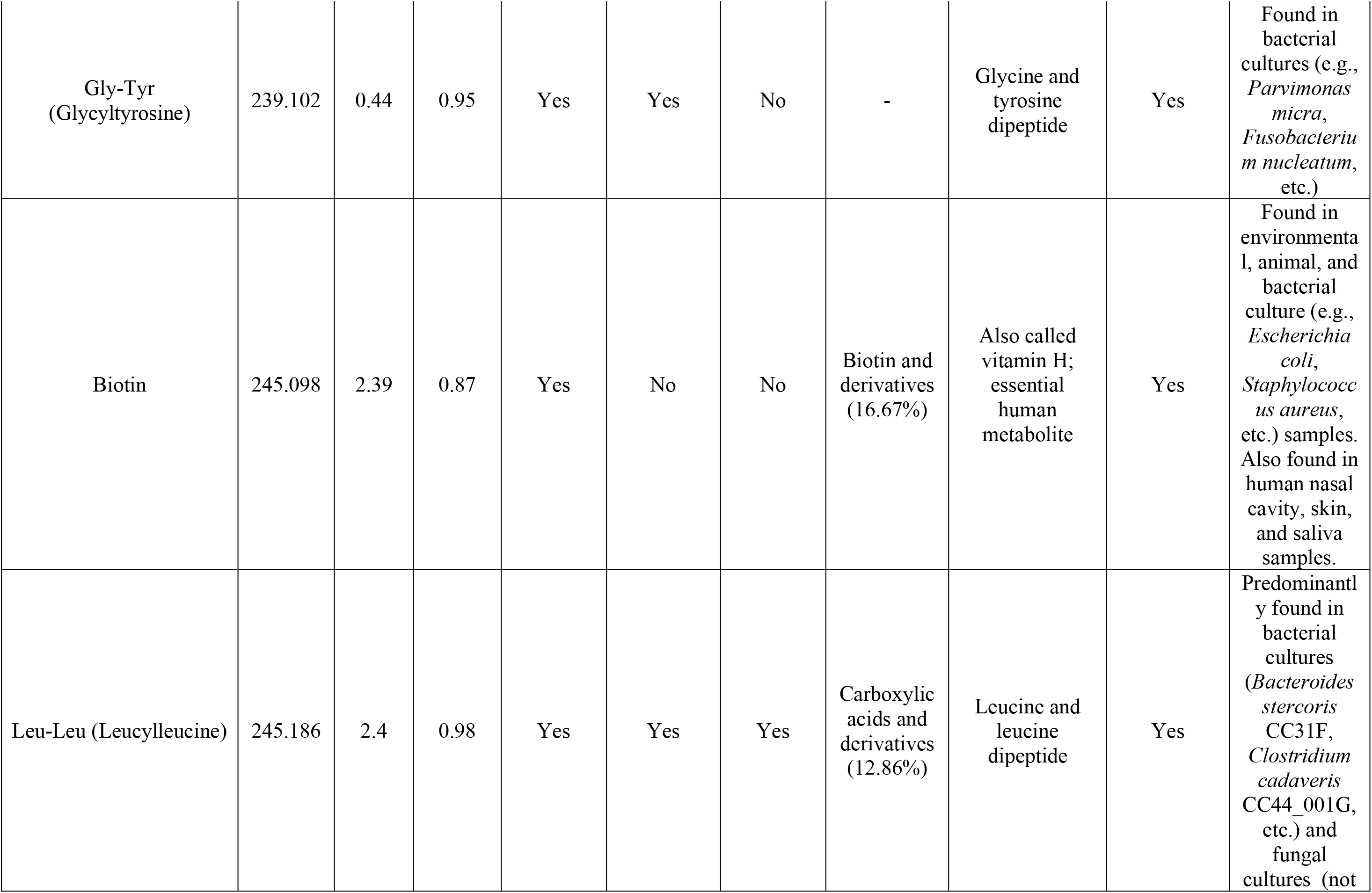
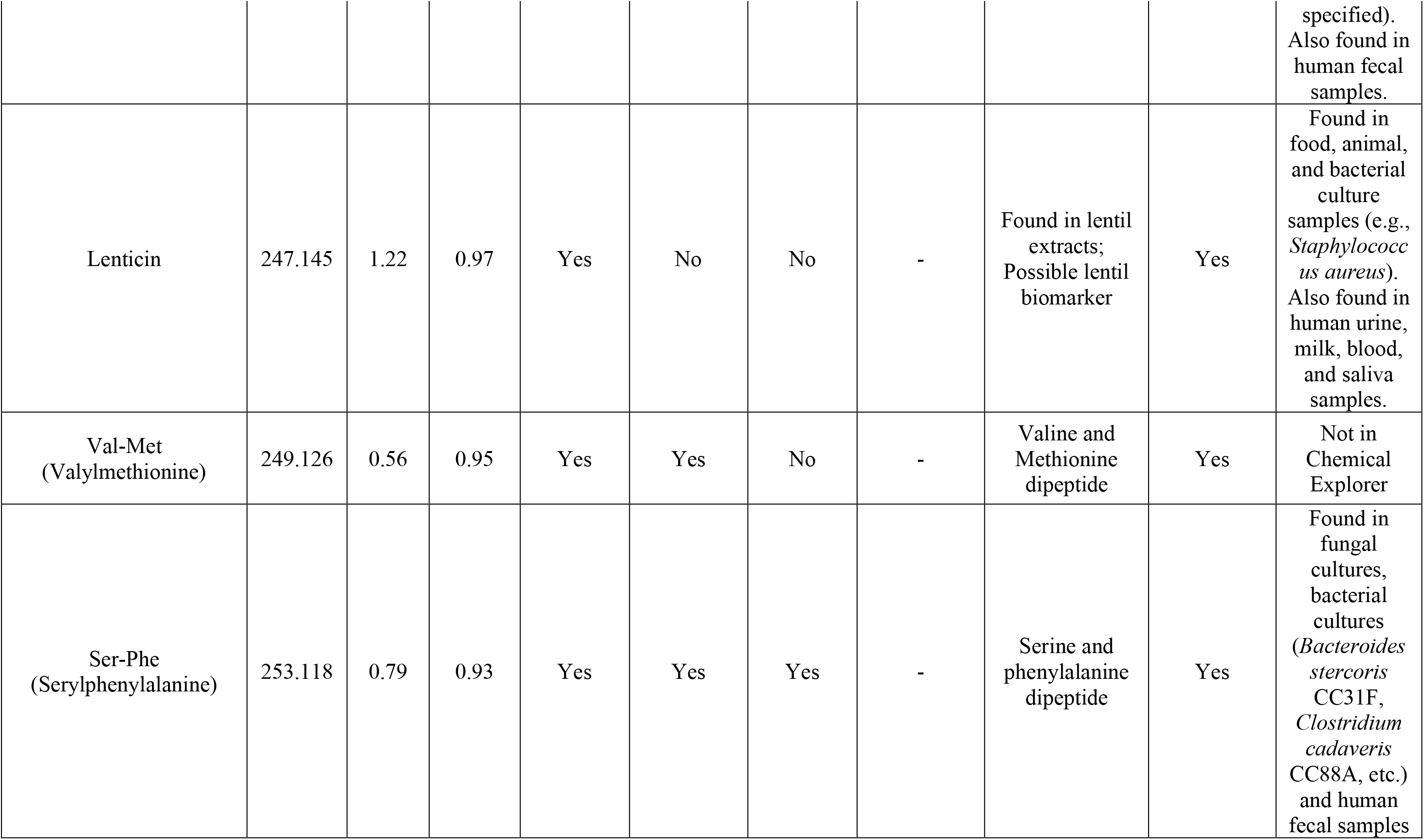
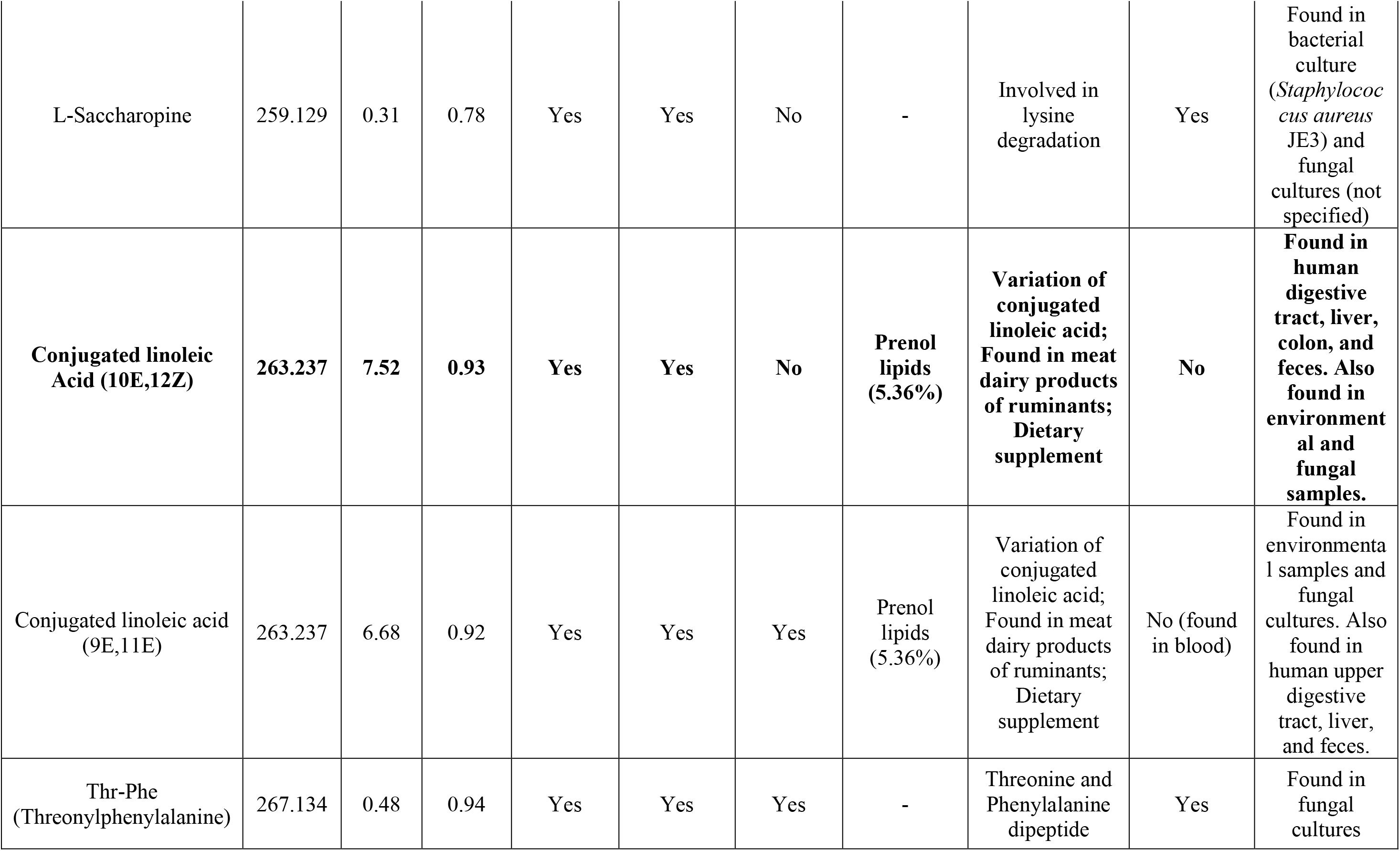
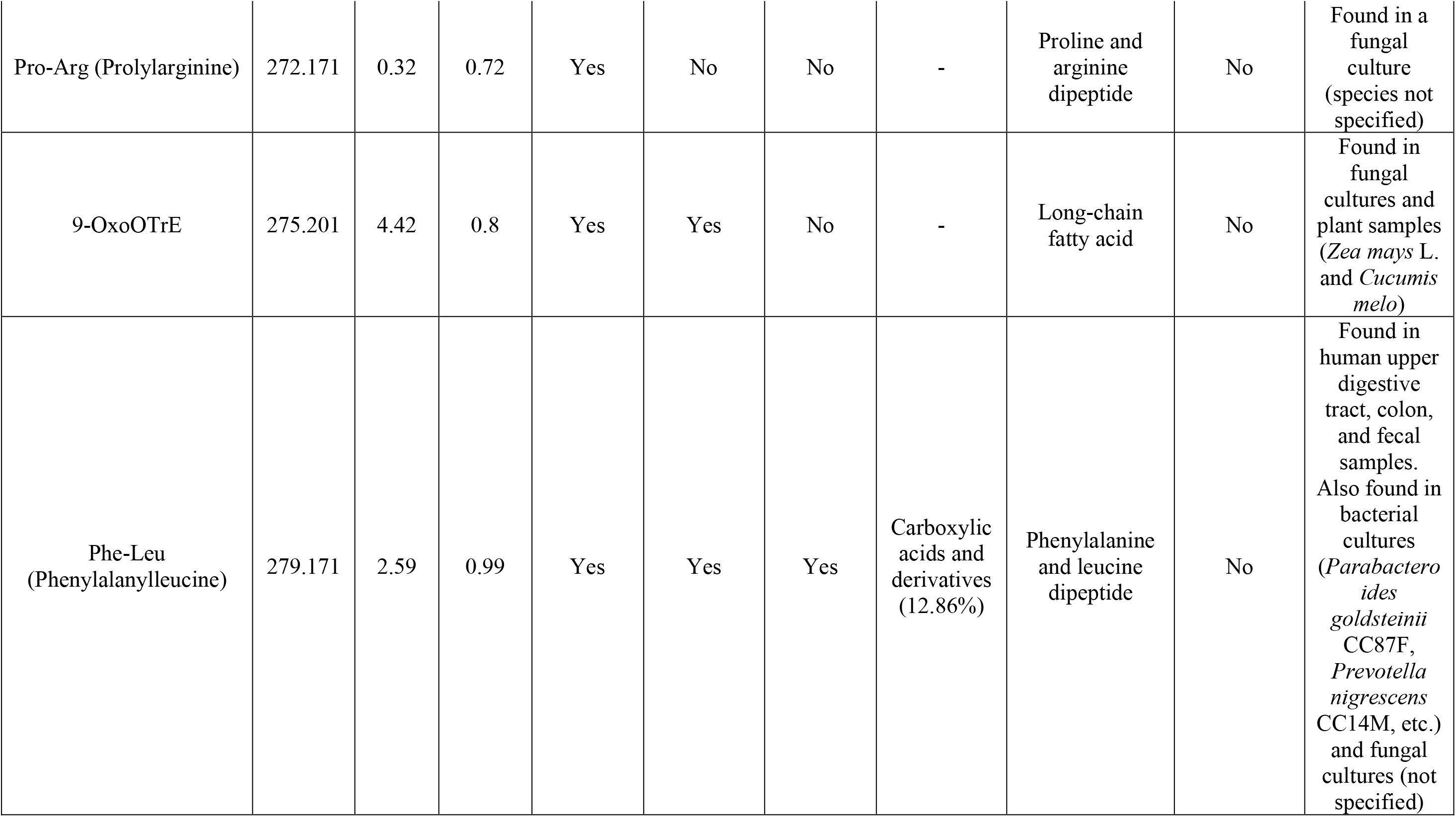
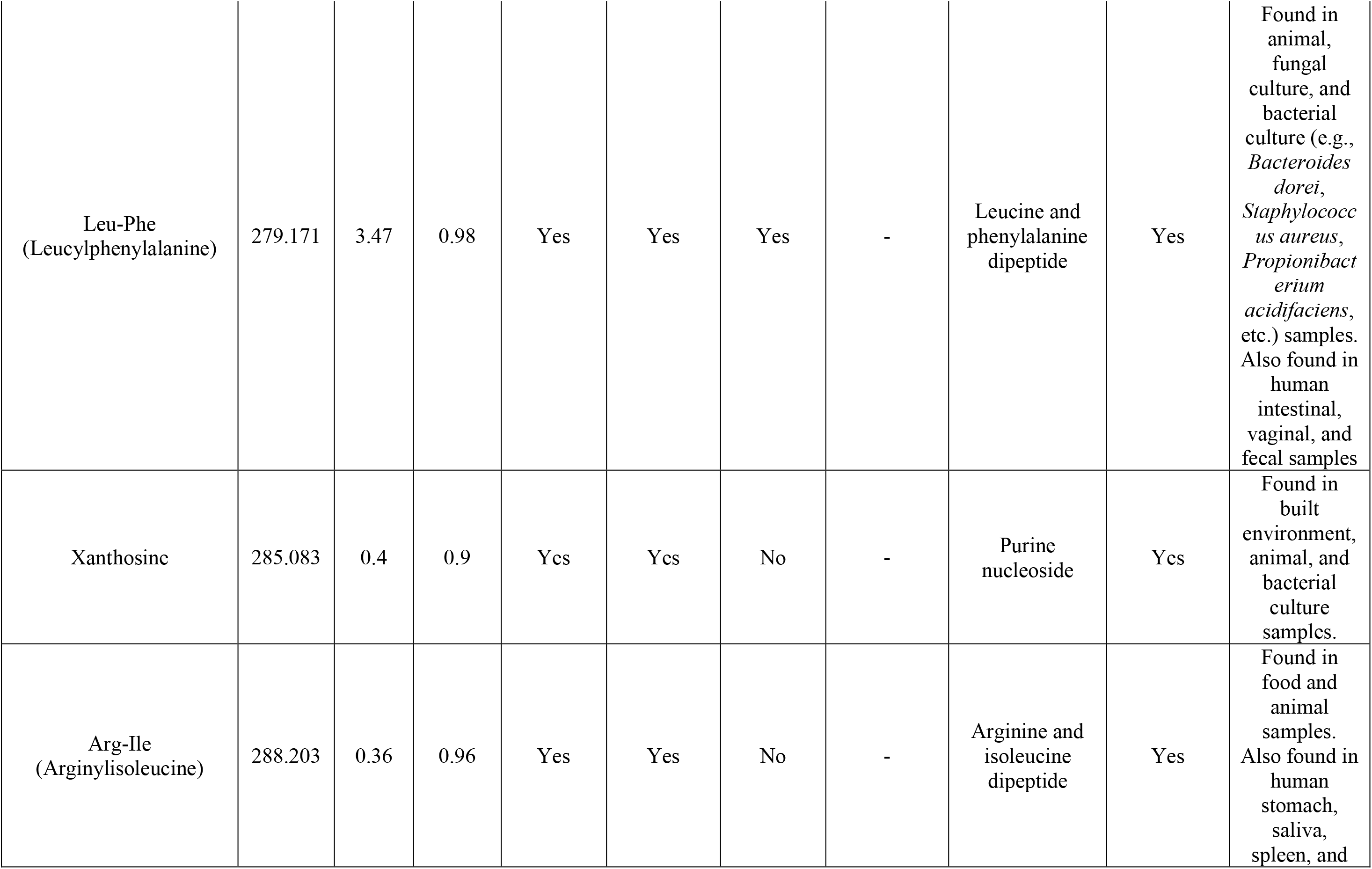
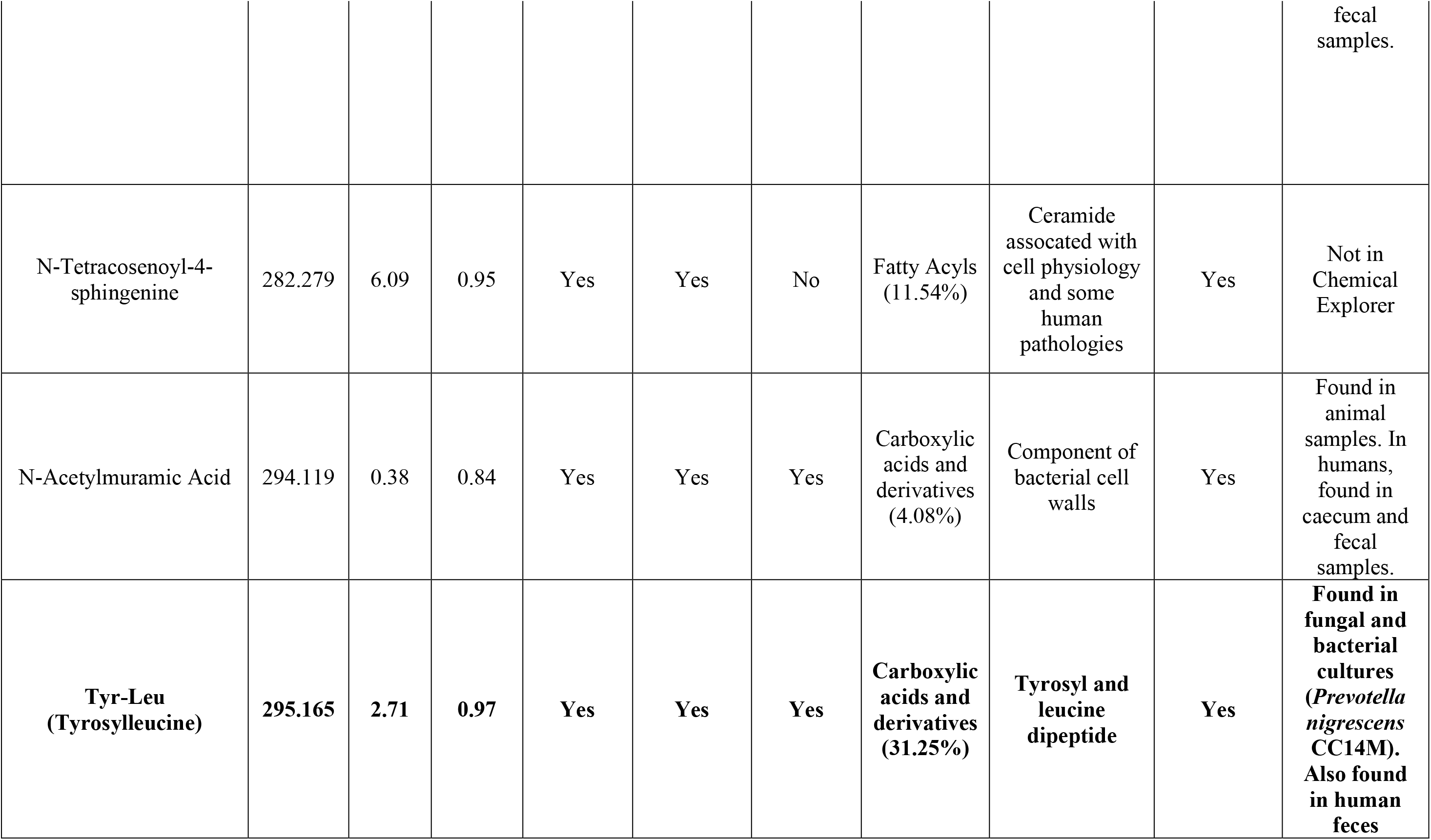
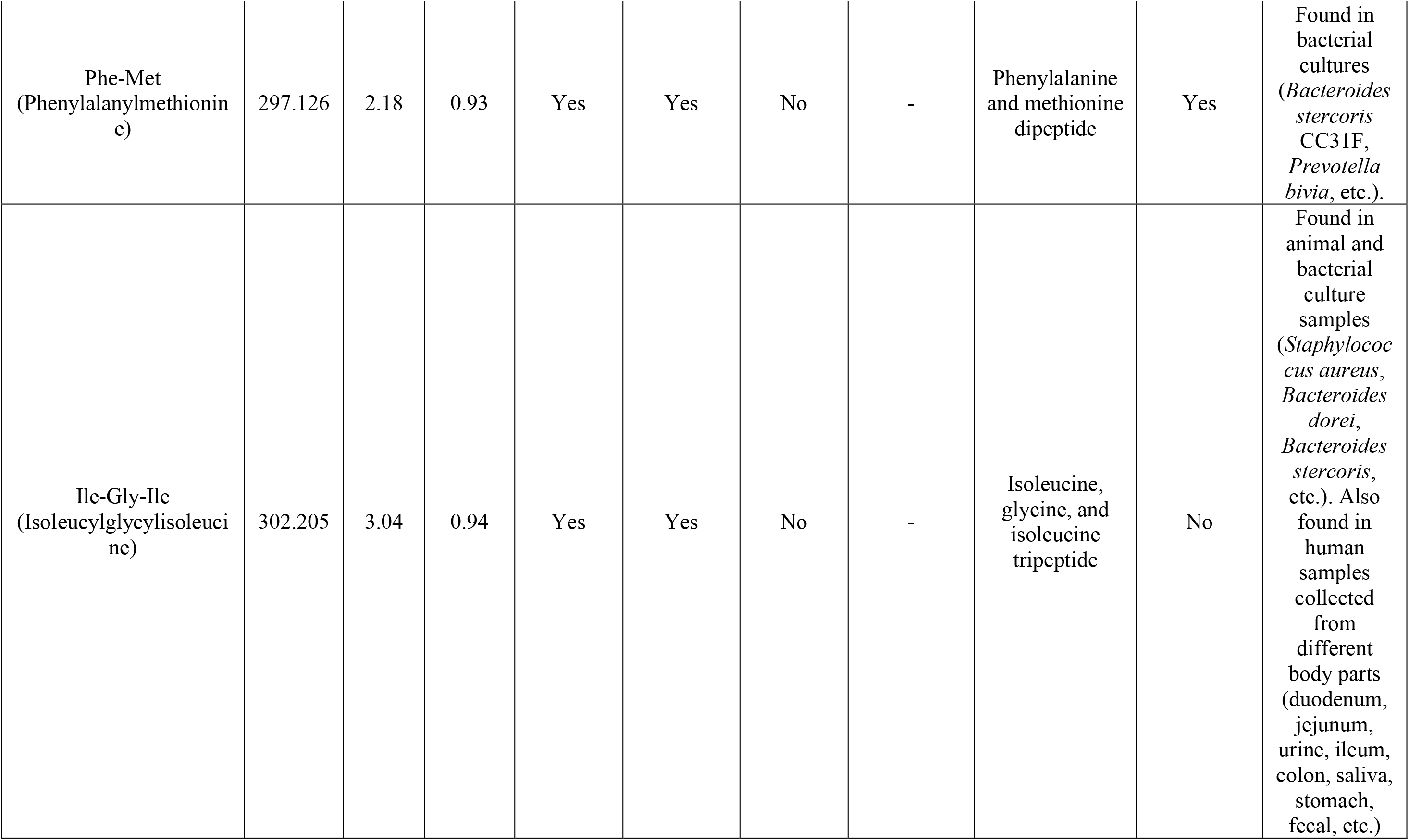
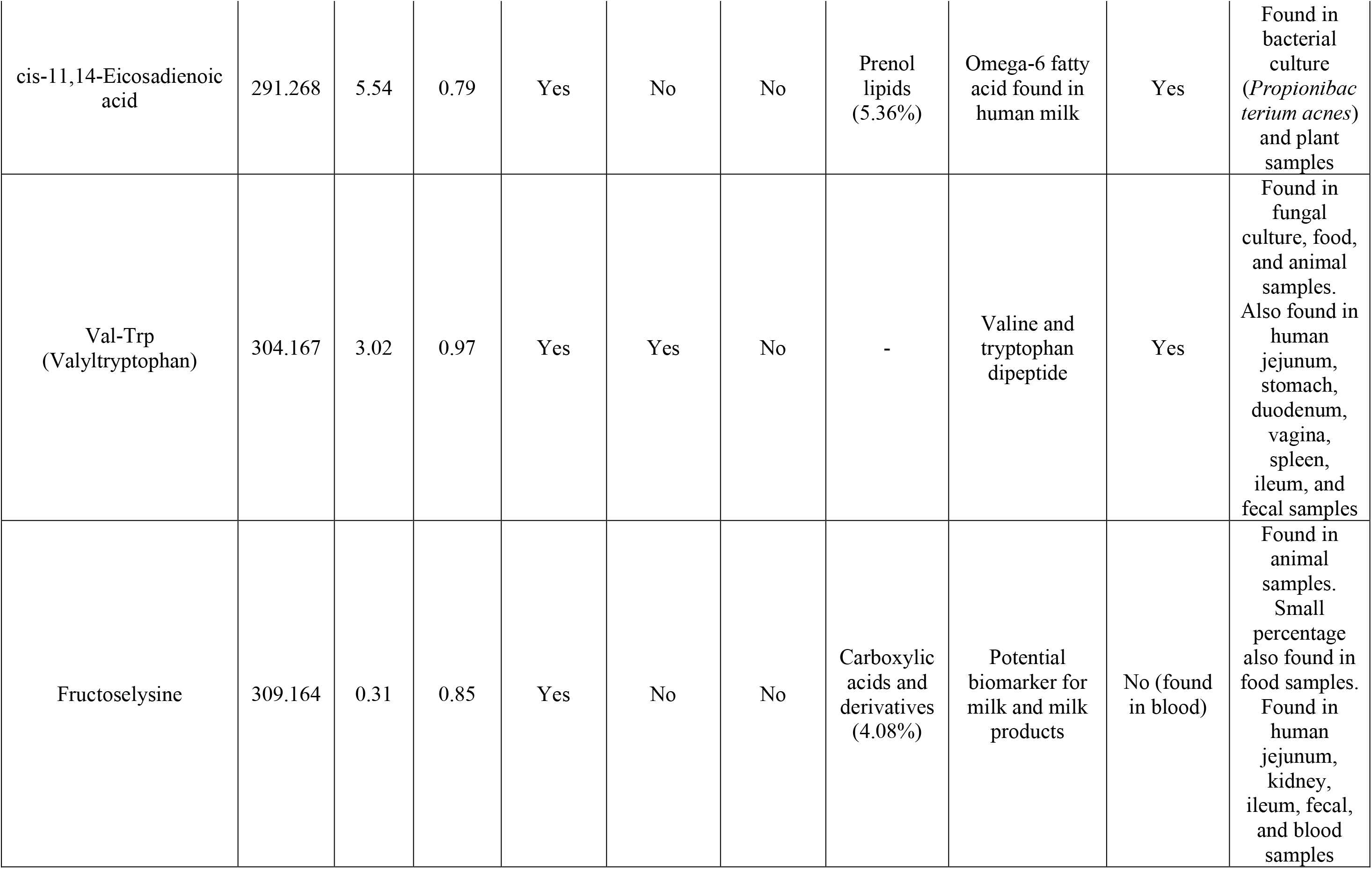
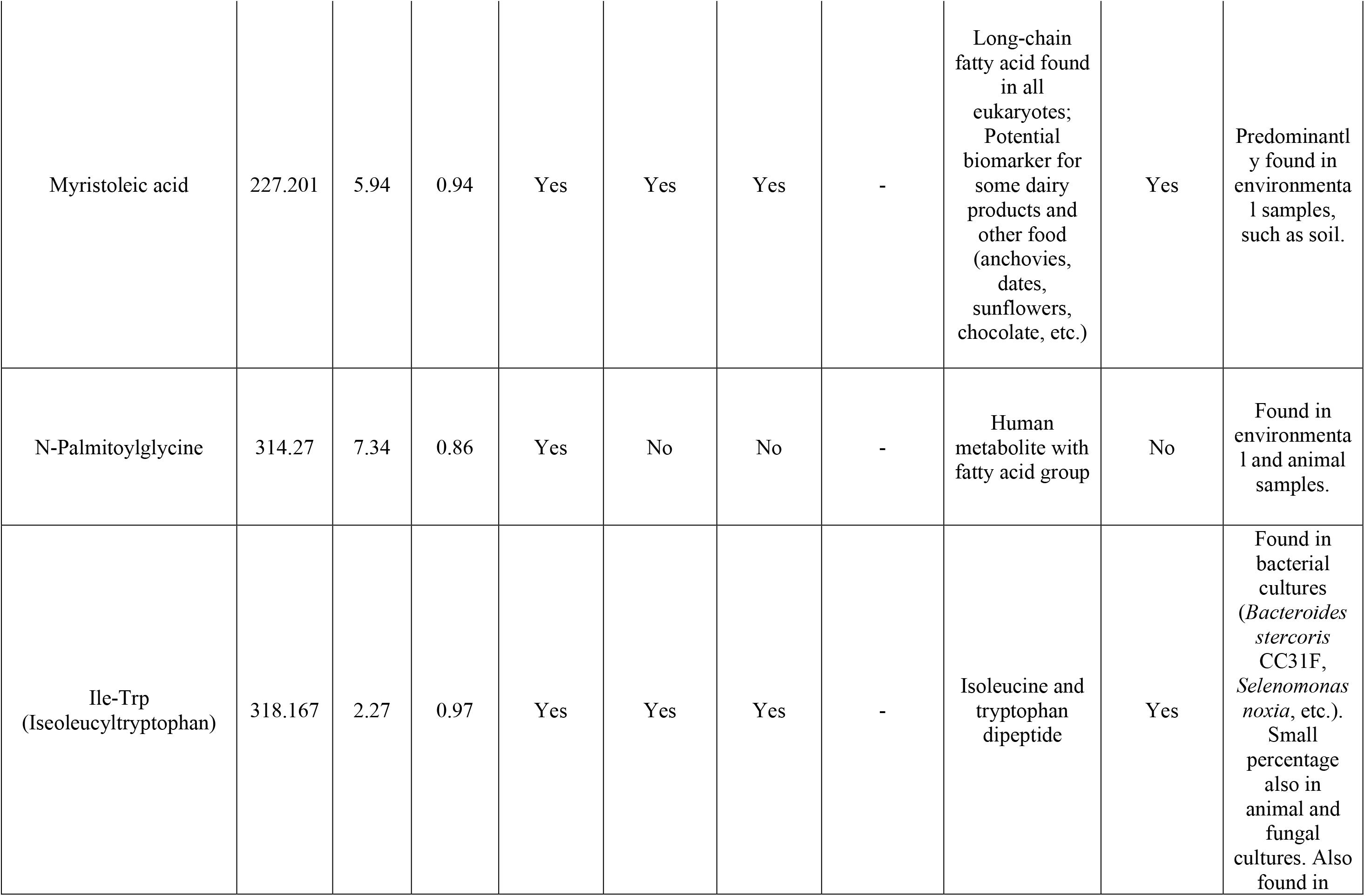
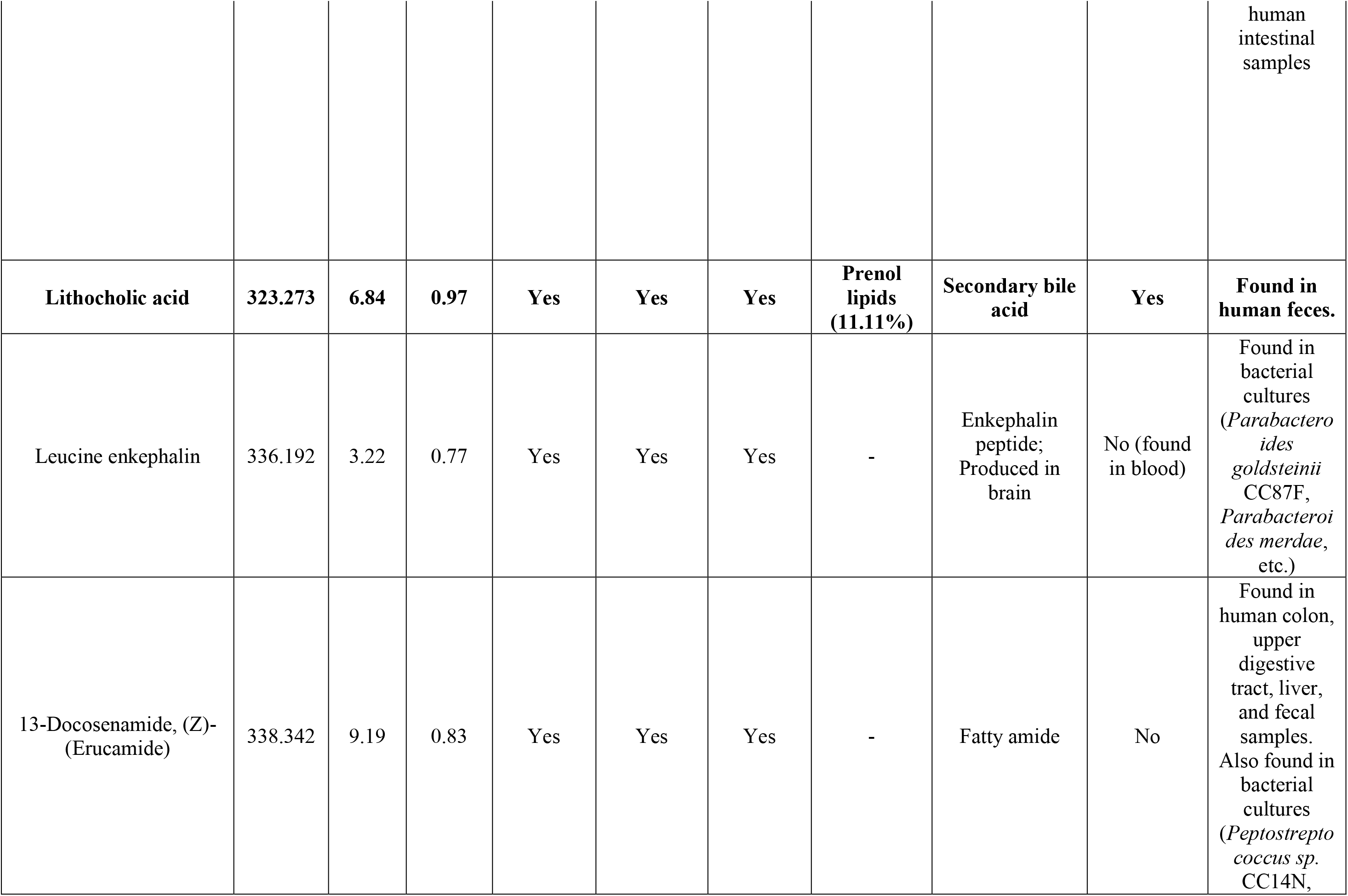
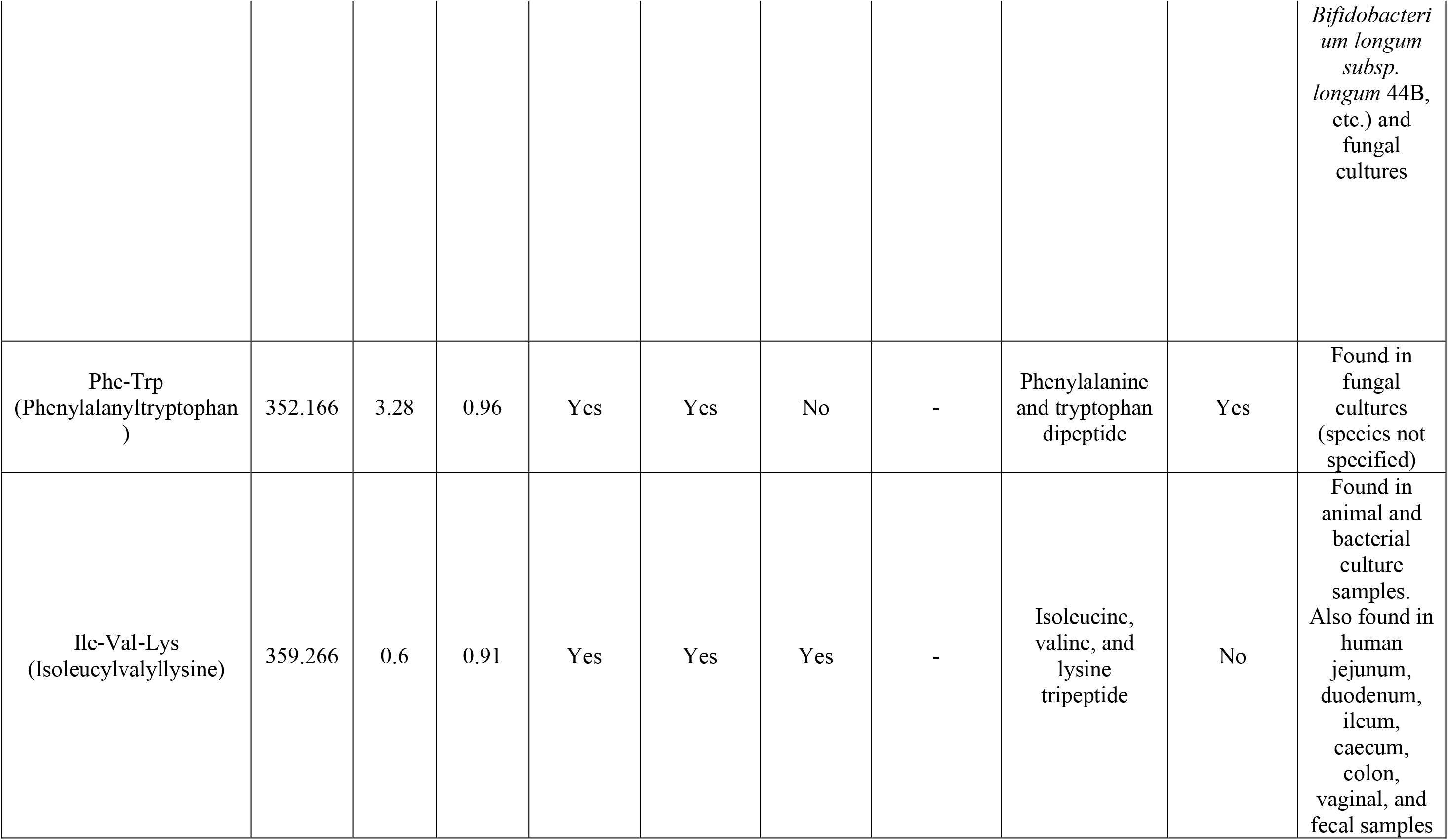
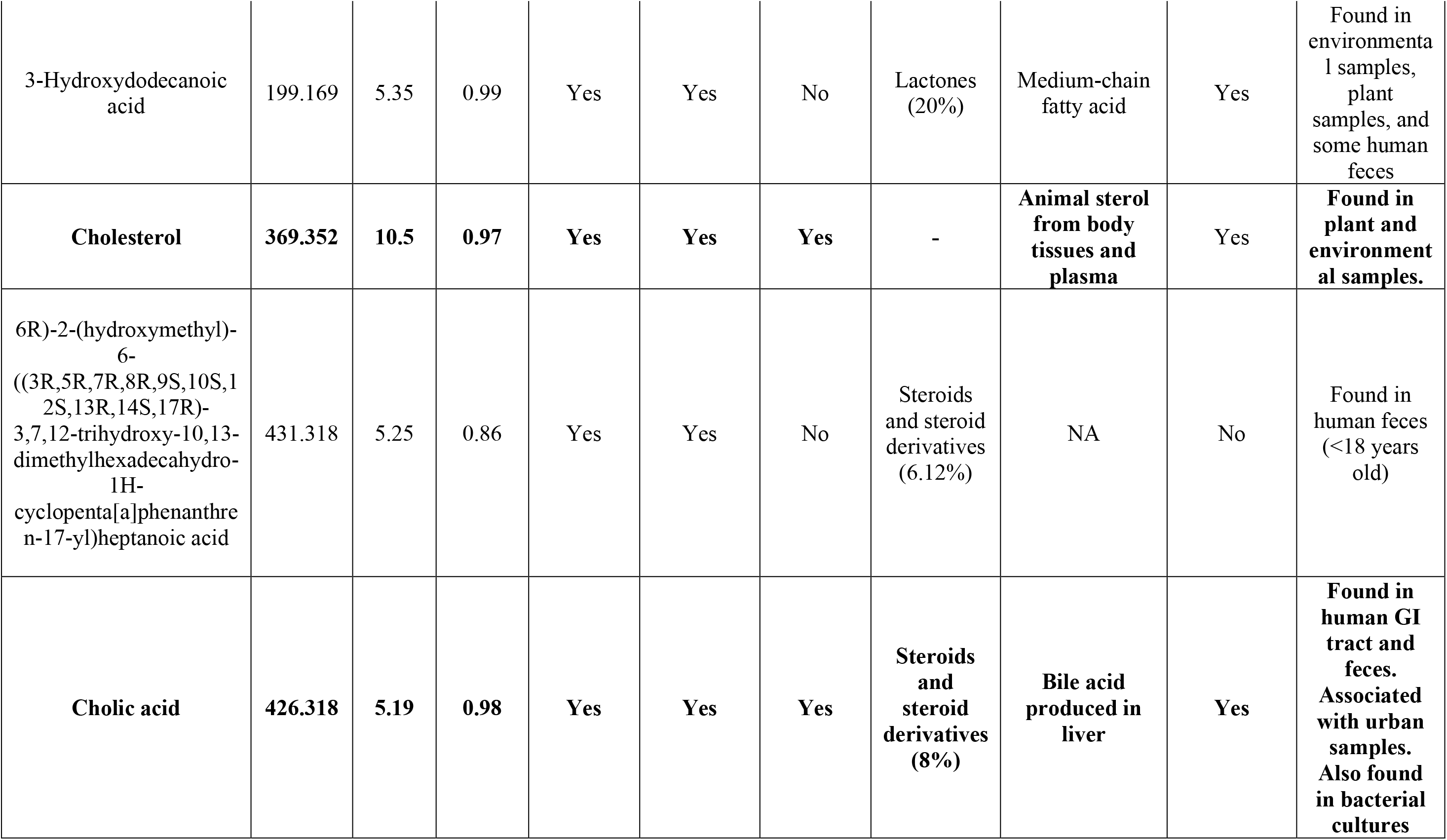
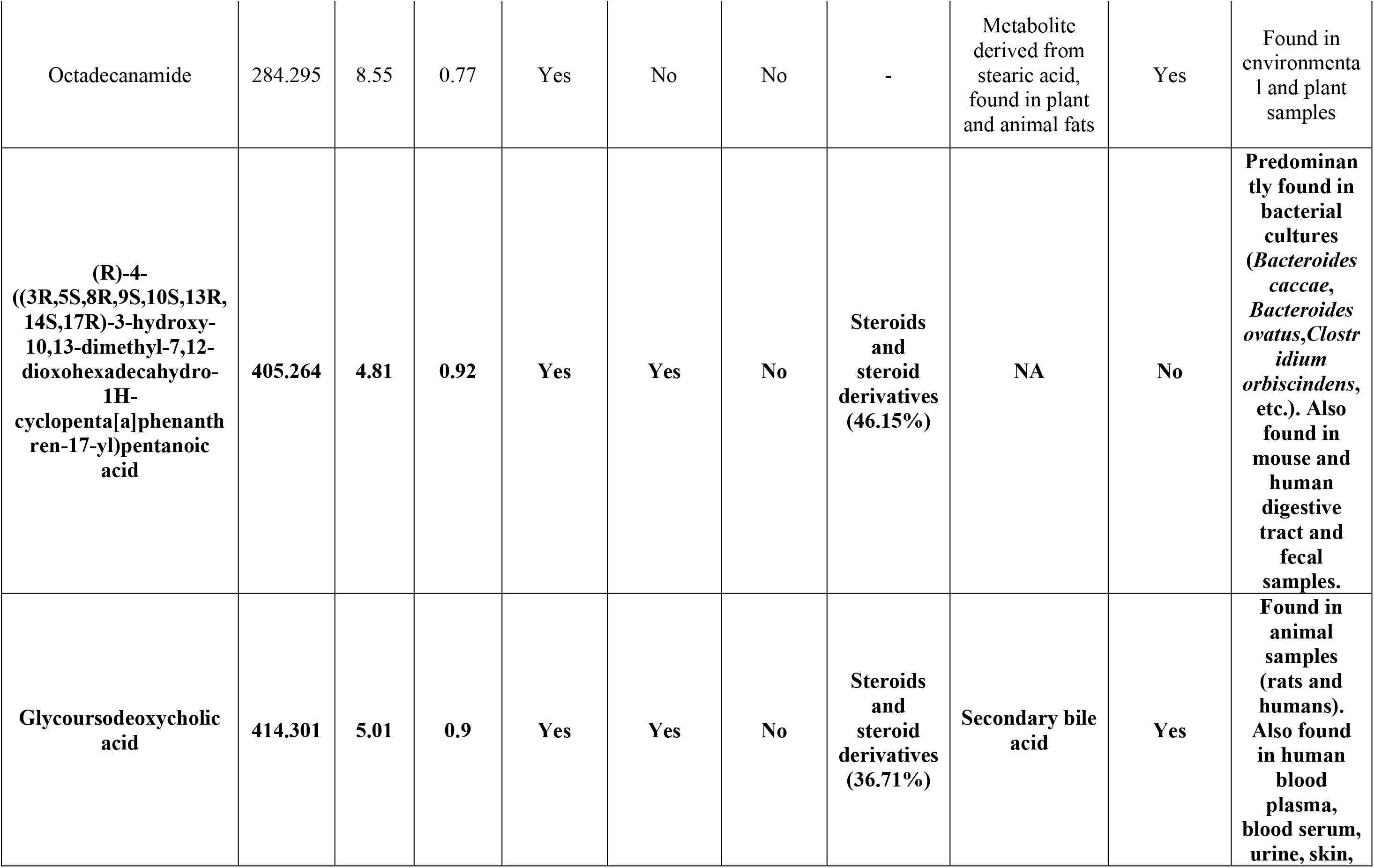
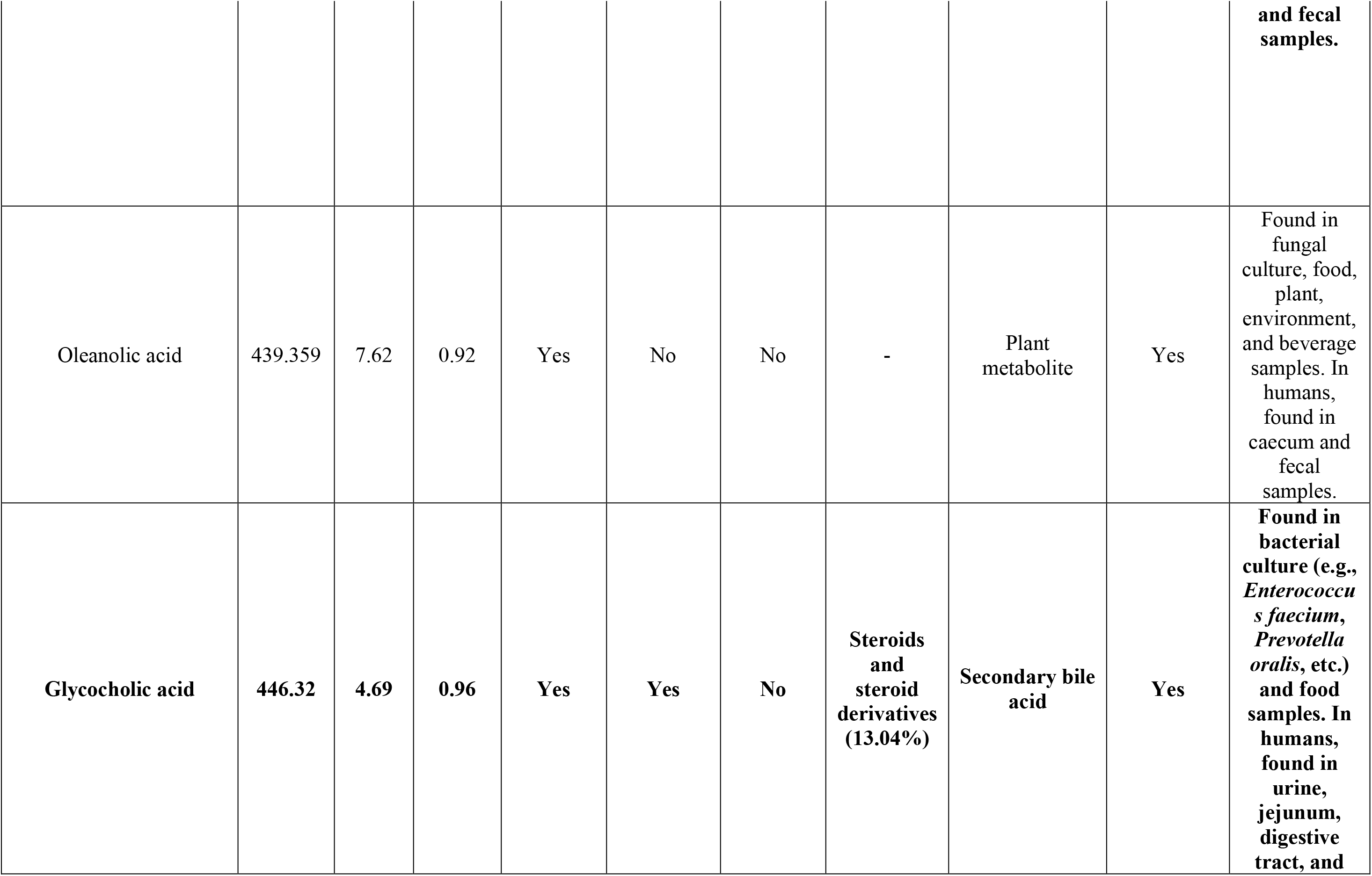
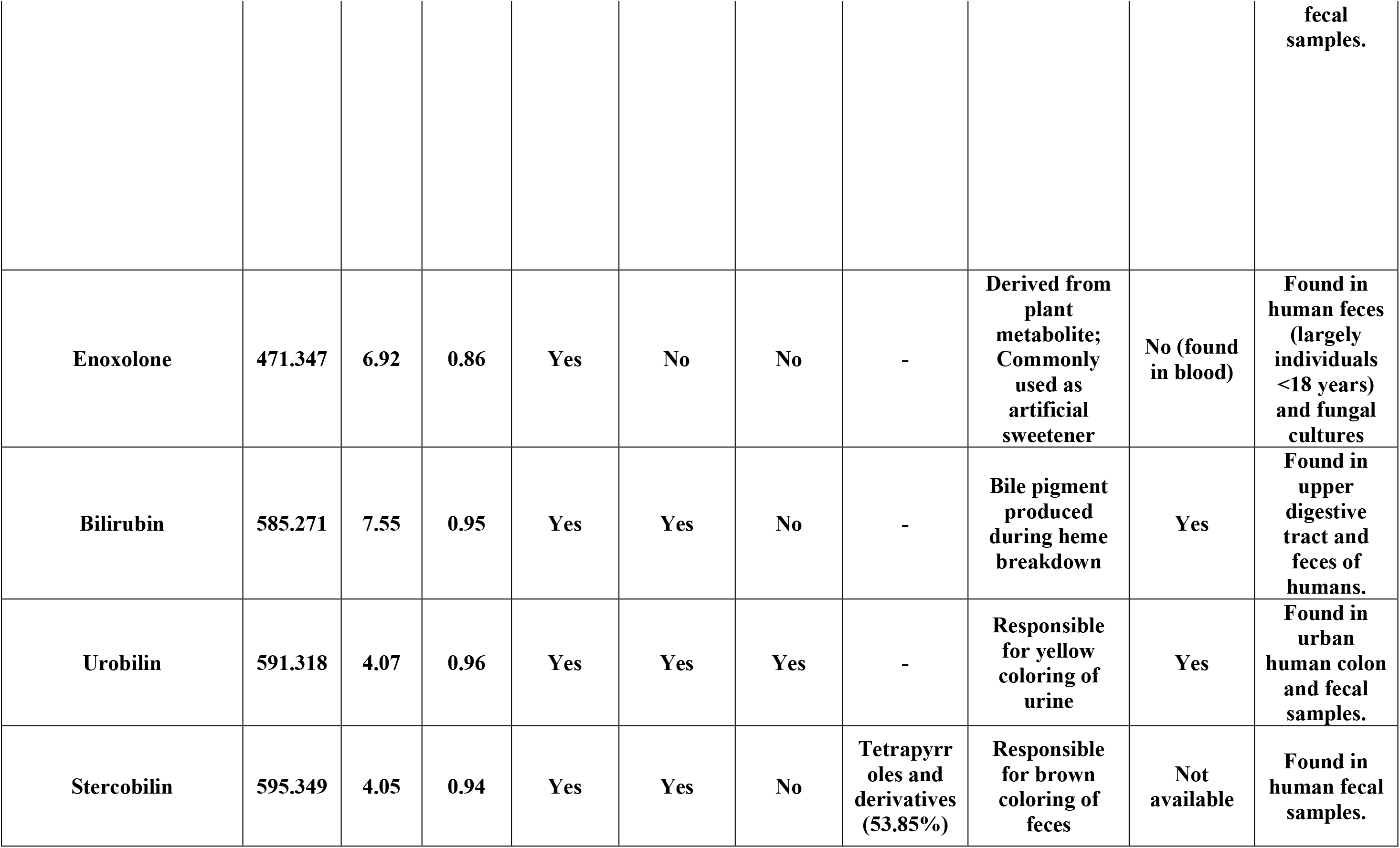
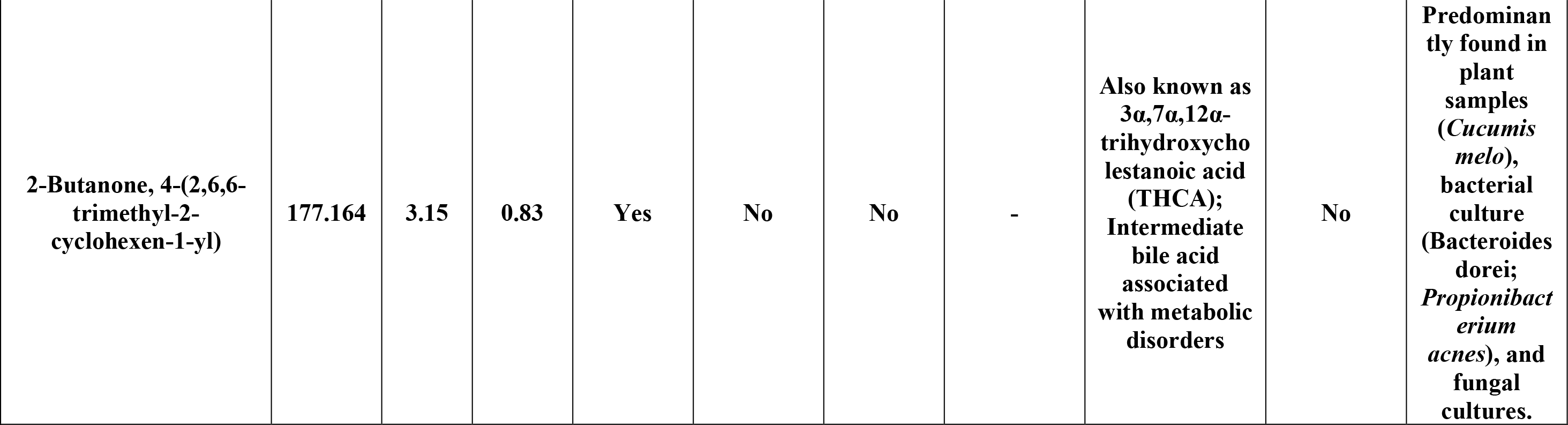
The core human fecal metabolome. All annotations had a mass difference of 0 to library reference. Bolded rows represent features detected in all ReDU datasets.

**Supplementary Table 3.** Correlated metabolite abundances for industrialization score groups.

| Metabolite Node *m/z* | Metabolite Node RT | Metabolite Annotation | Summed Metabolite Abundance at Industrialization Score 1 | Summed Metabolite Abundance at Industrialization Score 2 | Summed Metabolite Abundance at Industrialization Score 3 | Summed Metabolite Abundance at Industrialization Score 4 |
| --- | --- | --- | --- | --- | --- | --- |
| 137.046 | 0.3805 | Hypoxanthine | 2.49E+10 | 5.61E+10 | 1.01E+11 | 43640000000 |
| 139.0503 | 0.3108 | Nicotinamide N-oxide | 9.71E+09 | 1.62E+10 | 2.26E+10 | 8532100000 |
| 148.0759 | 3.2191 | 3-methyl-2-oxindole | 7.41E+08 | 2.14E+09 | 4.58E+09 | 1499345262 |
| 158.154 | 4.7767 | Hyochoic acid | 3.18E+08 | 6.46E+08 | 7.86E+08 | 398800000 |
| 177.0547 | 4.1326 | 3-Hydroxy-4-methoxycinnamic acid | 8.01E+07 | 1.13E+08 | 4.33E+08 | 226904586.4 |
| 177.1638 | 5.9084 | 2-Butanone, 4-(2,6,6-trimethyl-2-cyclohexen-1-yl)- | 4.17E+07 | 1.93E+07 | 2.78E+07 | 5170839.765 |
| 181.0721 | 0.807 | Paraxanthine | 1.09E+09 | 1.51E+08 | 3.68E+08 | 6218745.074 |
| 195.0651 | 3.0126 | trans-Ferulic acid | 3.59E+08 | 2.30E+08 | 1.40E+09 | 1096021152 |
| 197.1165 | 3.0981 | Loliolide | 1.76E+07 | 1.11E+08 | 3.75E+08 | 80086416.57 |
| 199.169 | 5.3476 | 3-Hydroxydodecanoic acid | 8.72E+07 | 3.33E+07 | 3.68E+07 | 29025942.95 |
| 204.0868 | 3.5276 | N-Acetyl-D-mannosamine | 1.78E+08 | 6.88E+08 | 9.54E+08 | 960951142.4 |
| 208.097 | 2.8411 | L-Phenylalanine, N-acetyl- | 3.20E+08 | 1.40E+09 | 2.04E+09 | 950700000 |
| 217.1224 | 0.4618 | Thr-Pro | 3.33E+07 | 8.05E+07 | 1.42E+08 | 63401131.12 |
| 217.155 | 0.4485 | Val-Val | 8.18E+09 | 6.45E+09 | 2.96E+10 | 16681356835 |
| 220.1183 | 0.5571 | Pantothenic acid | 4.84E+09 | 8.72E+09 | 1.52E+10 | 5026000000 |
| 227.2005 | 5.943 | Myristoleic acid | 4.25E+07 | 3.65E+07 | 8.52E+07 | 62295962.54 |
| 229.155 | 0.7773 | Ile-Pro | 4.69E+10 | 5.93E+10 | 1.27E+11 | 65216000000 |
| 231.171 | 2.8229 | Val-Ile | 1.28E+09 | 1.96E+09 | 8.75E+09 | 5716297868 |
| 237.2214 | 6.4871 | cis-9-Hexadecenoic acid | 1.40E+08 | 7.36E+07 | 9.26E+07 | 203983532.6 |
| 239.1021 | 0.4404 | Gly-Tyr | 2.61E+08 | 5.24E+08 | 1.40E+09 | 1028935149 |
| 245.0983 | 2.3864 | Biotin | 6.21E+07 | 4.93E+07 | 1.84E+08 | 42932706.06 |
| 245.1862 | 2.3952 | Leu-Leu | 2.52E+09 | 3.45E+09 | 1.35E+10 | 9520700000 |
| 247.1448 | 1.2228 | Lenticin | 1.52E+08 | 8.26E+07 | 2.27E+08 | 25378613.53 |
| 249.1264 | 0.5559 | Val-Met | 1.07E+09 | 7.64E+08 | 2.35E+09 | 948787911.6 |
| 263.2369 | 6.6799 | Conjugated linoleic acid (9E,11E) | 3.87E+08 | 4.46E+08 | 1.19E+09 | 324224225.4 |
| 263.237 | 7.5172 | Conjugated linoleic acid<br>(10E,12Z) | 1.54E+08 | 1.25E+08 | 4.39E+08 | 128896538.2 |
| 267.1342 | 0.4803 | Thr-Phe | 5.25E+08 | 6.05E+08 | 1.57E+09 | 482260542.5 |
| 272.1711 | 0.3211 | Pro-Arg | 2.32E+08 | 2.82E+08 | 6.76E+08 | 418814748.6 |
| 275.201 | 4.4249 | 9-OxoOTrE | 3.78E+07 | 5.82E+07 | 9.97E+07 | 69862085.1 |
| 279.1708 | 3.4667 | Leu-Phe | 4.94E+08 | 5.45E+08 | 1.97E+09 | 1723342465 |
| 282.2791 | 6.0892 | N-Tetracosenoyl-4-sphingenine | 6.14E+07 | 5.07E+07 | 5.29E+07 | 20546244.6 |
| 285.0831 | 0.4016 | Xanthosine | 1.32E+08 | 3.20E+08 | 1.15E+09 | 274911286.4 |
| 288.2028 | 0.3645 | Arg-Ile | 5.46E+08 | 1.20E+09 | 5.44E+09 | 3715539067 |
| 291.268 | 5.5386 | cis-11,14-Eicosadienoic acid | 4.96E+07 | 3.97E+07 | 2.07E+07 | 805186.4409 |
| 294.1191 | 0.3764 | N-Acetylmuramic acid | 5.36E+08 | 4.07E+09 | 6.46E+09 | 1877551752 |
| 297.1261 | 3.1402 | Phe-Met | 9.42E+06 | 3.49E+07 | 7.51E+07 | 23615664.4 |
| 302.2048 | 3.0495 | Ile-Gly-Ile | 2.11E+07 | 2.41E+07 | 6.02E+07 | 12623338.92 |
| 304.1663 | 3.017 | Val-Trp | 9.20E+07 | 2.01E+08 | 5.42E+08 | 370977408.1 |
| 309.1635 | 0.3129 | Fructoselysine | 8.26E+07 | 1.01E+08 | 2.48E+08 | 229202252.1 |
| 318.1674 | 2.2699 | Ile-Trp | 4.22E+06 | 4.16E+06 | 2.50E+06 | 5691574.455 |
| 323.2734 | 6.8392 | Lithocholic acid | 7.89E+07 | 1.12E+08 | 6.91E+07 | 43717480.88 |
| 336.1923 | 3.2198 | Leucine Enkephalin | 8.38E+08 | 5.57E+08 | 9.79E+08 | 756900000 |
| 338.3419 | 9.1869 | 13-Docosenamide, (Z)- | 1.21E+08 | 1.87E+08 | 3.05E+08 | 163546639 |
| 359.2661 | 0.5962 | Ile-Val-Lys | 6.59E+08 | 5.47E+08 | 1.27E+09 | 796505678 |
| 369.3517 | 10.5045 | Cholesterol | 9.83E+08 | 2.74E+08 | 4.91E+08 | 256617172 |
| 405.2639 | 4.8148 | (R)-4-<br>((3R,5S,8R,9S,10S,13R,14S,17R)-<br>3-hydroxy-10,13-dimethyl-7,12-dioxohexadecahydro-1H-cyclopenta[a]phenanthren-17-yl)pentanoic acid | 1.08E+08 | 2.56E+08 | 1.66E+08 | 1487753381 |
| 439.3585 | 7.6154 | Oleanolic acid | 4.07E+08 | 4.28E+08 | 1.89E+08 | 24997439.71 |
| 471.347 | 6.9183 | Enoxolone | 7.79E+08 | 1.08E+08 | 4.68E+07 | 9346458.261 |
| 585.2723 | 8.8966 | Bilirubin | 1.51E+08 | 8.39E+07 | 1.11E+07 | 2115169.671 |
| 591.3181 | 4.0748 | Urobilin | 7.68E+10 | 5.65E+09 | 1.70E+09 | 3131747944 |
| 595.3486 | 4.0491 | Stercobilin | 1.36E+11 | 1.23E+11 | 2.19E+11 | 47140614012 |
| 839.5646 | 5.1216 | Cholic acid | 6.37E+08 | 2.53E+09 | 4.27E+08 | 258646224.3 |

**Supplementary Table 4.**
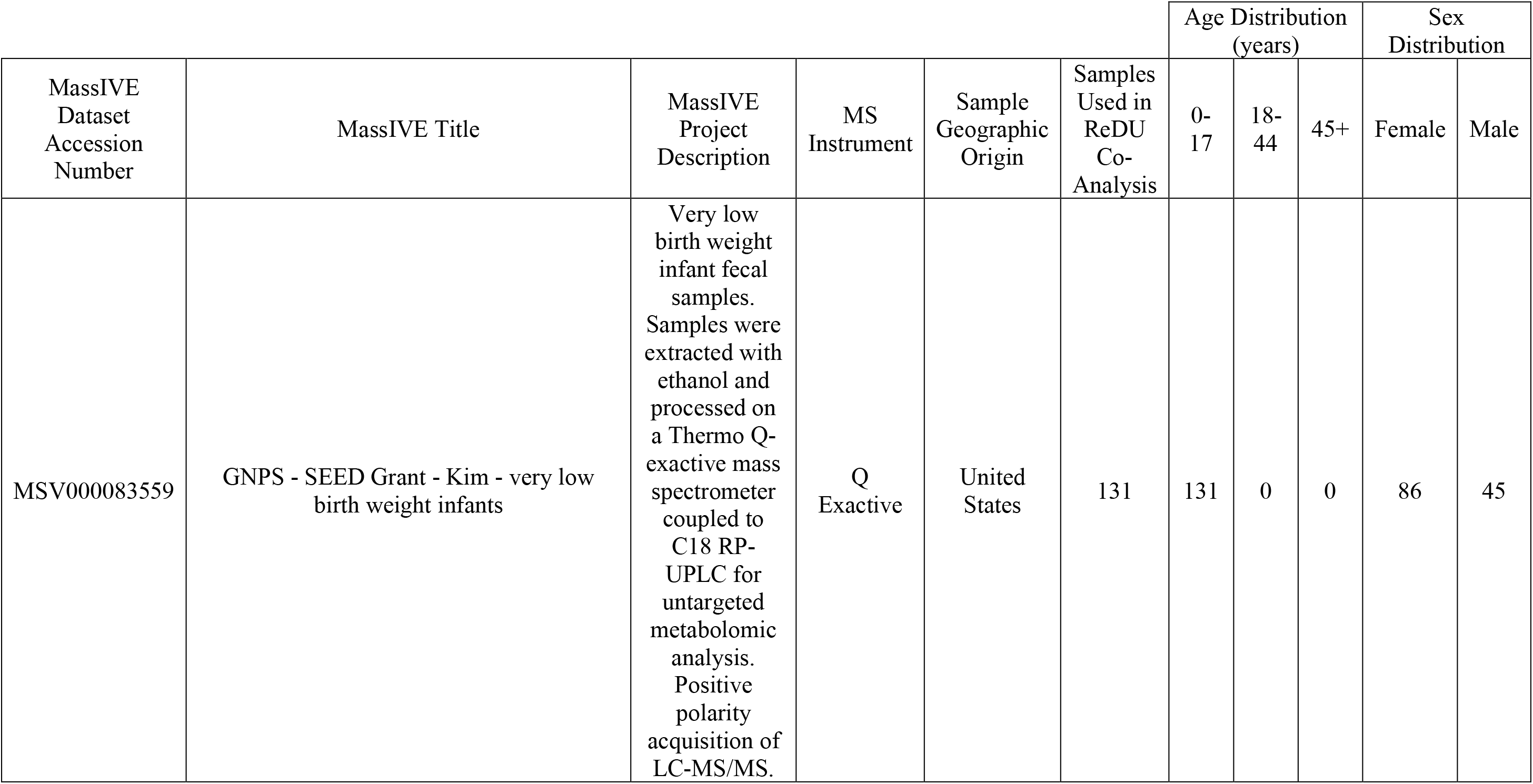
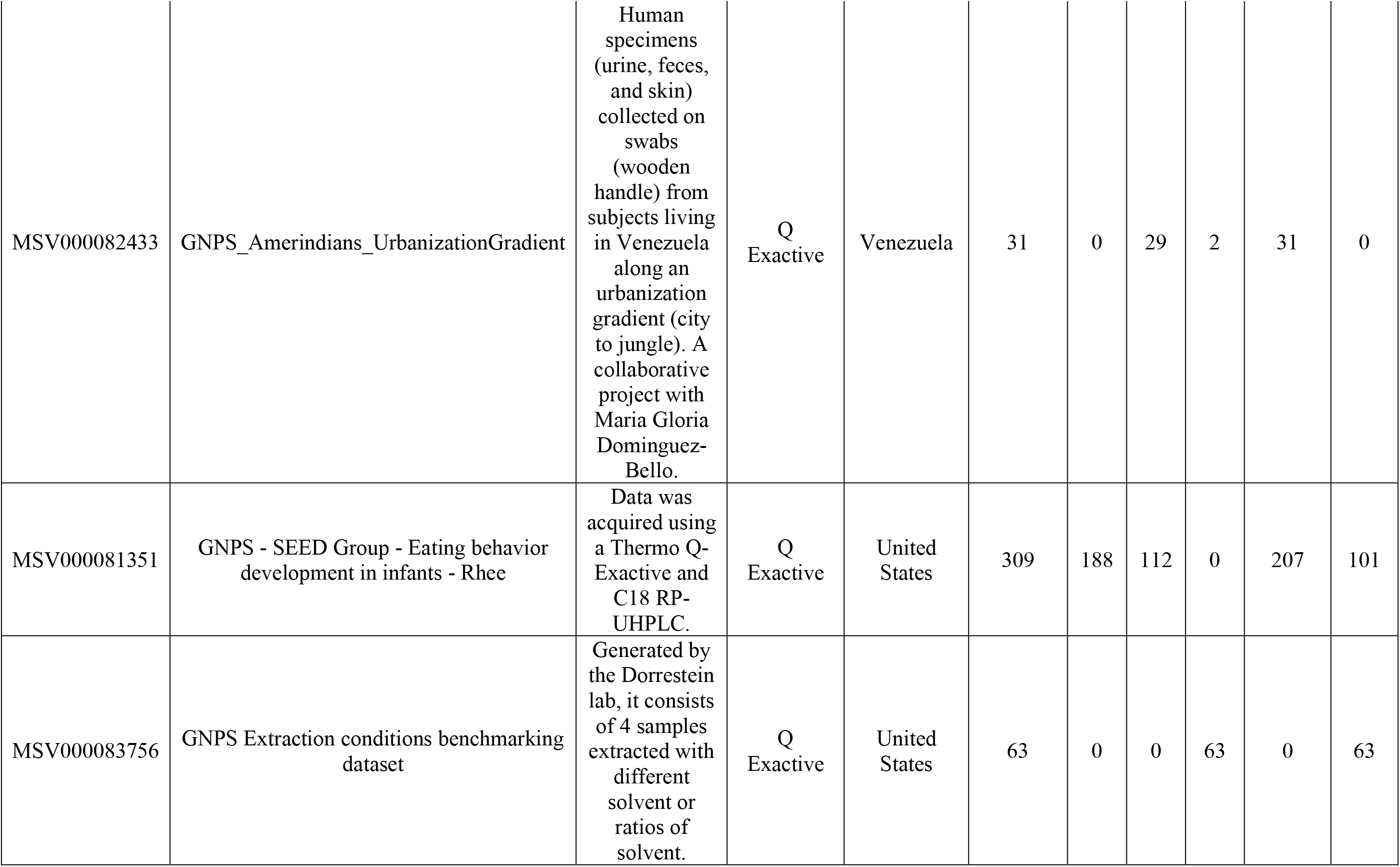
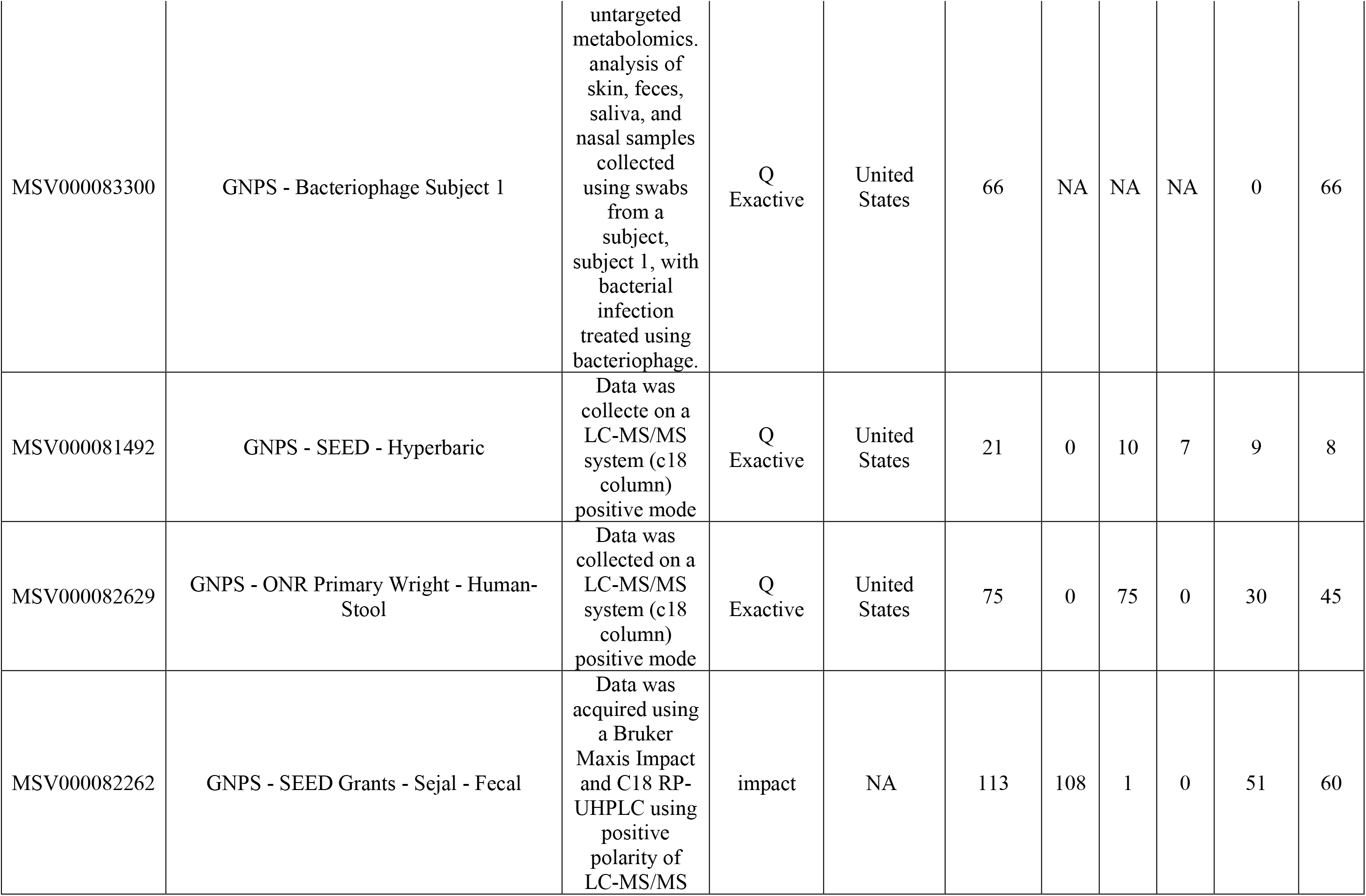
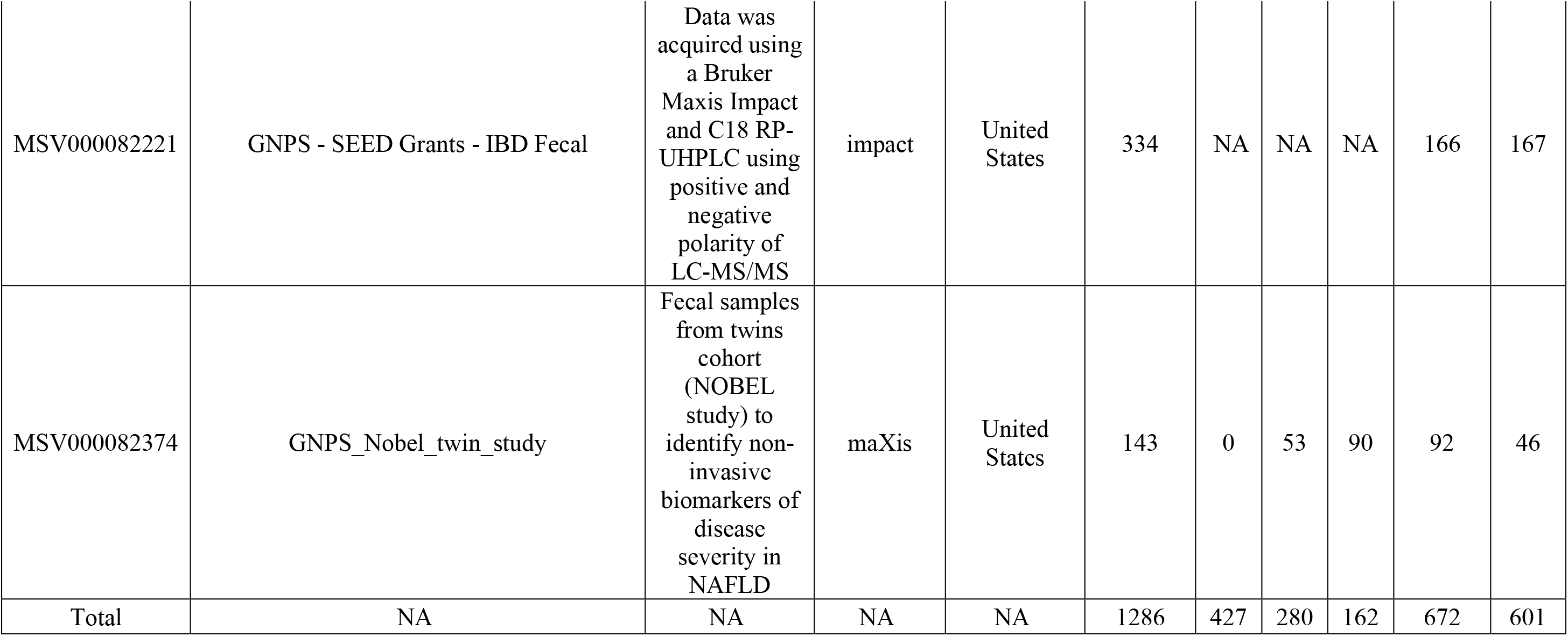
Public MassIVE datasets used for ReDU co-analysis. NA values represent data labeled “not collected”.

| MassIVE<br>Dataset<br>Accession<br>Number | MassIVE Title | MassIVE<br>Project<br>Description | MS<br>Instrument | Sample<br>Geographic<br>Origin | Samples<br>Used in<br>ReDU<br>Co-Analysis | Age Distribution<br>(years) |  |  | Sex<br>Distribution |  |
| --- | --- | --- | --- | --- | --- | --- | --- | --- | --- | --- |
|  |  |  |  |  |  | 0-17 | 18-44 | 45+ | Female | Male |
| MSV000083559 | GNPS - SEED Grant - Kim - very low birth weight infants | Very low birth weight infant fecal samples. Samples were extracted with ethanol and processed on a Thermo Q-exactive mass spectrometer coupled to C18 RP-UPLC for untargeted metabolomic analysis. Positive polarity acquisition of LC-MS/MS. | Q<br>Exactive | United States | 131 | 131 | 0 | 0 | 86 | 45 |
| MSV000082433 | GNPS_Amerindians_UrbanizationGradient | Human specimens (urine, feces, and skin) collected on swabs (wooden handle) from subjects living in Venezuela along an urbanization gradient (city to jungle). A collaborative project with Maria Gloria Dominguez-Bello. | Q<br>Exactive | Venezuela | 31 | 0 | 29 | 2 | 31 | 0 |
| MSV000081351 | GNPS - SEED Group - Eating behavior development in infants - Rhee | Data was acquired using a Thermo Q-Exactive and C18 RP-UHPLC. | Q<br>Exactive | United States | 309 | 188 | 112 | 0 | 207 | 101 |
| MSV000083756 | GNPS Extraction conditions benchmarking dataset | Generated by the Dorrestein lab, it consists of 4 samples extracted with different solvent or ratios of solvent. | Q<br>Exactive | United States | 63 | 0 | 0 | 63 | 0 | 63 |
| MSV000083300 | GNPS - Bacteriophage Subject 1 | untargeted metabolomics. analysis of skin, feces, saliva, and nasal samples collected using swabs from a subject, subject 1, with bacterial infection treated using bacteriophage. | Q<br>Exactive | United States | 66 | NA | NA | NA | 0 | 66 |
| MSV000081492 | GNPS - SEED - Hyperbaric | Data was collecte on a LC-MS/MS system (c18 column) positive mode | Q<br>Exactive | United States | 21 | 0 | 10 | 7 | 9 | 8 |
| MSV000082629 | GNPS - ONR Primary Wright - Human-Stool | Data was collected on a LC-MS/MS system (c18 column) positive mode | Q<br>Exactive | United States | 75 | 0 | 75 | 0 | 30 | 45 |
| MSV000082262 | GNPS - SEED Grants - Sejal - Fecal | Data was acquired using a Bruker Maxis Impact and C18 RP-UHPLC using positive polarity of LC-MS/MS | impact | NA | 113 | 108 | 1 | 0 | 51 | 60 |
| MSV000082221 | GNPS - SEED Grants - IBD Fecal | Data was acquired using a Bruker Maxis Impact and C18 RP-UHPLC using positive and negative polarity of LC-MS/MS | impact | United States | 334 | NA | NA | NA | 166 | 167 |
| MSV000082374 | GNPS_Nobel_twin_study | Fecal samples from twins cohort (NOBEL study) to identify non-invasive biomarkers of disease severity in NAFLD | maXis | United States | 143 | 0 | 53 | 90 | 92 | 46 |
| Total | NA | NA | NA | NA | 1286 | 427 | 280 | 162 | 672 | 601 |

**Supplementary Table 5.** MZmine parameters for feature-based molecular networking.

**MZmine v 2.33**
| Method | Parameter | Value |
| --- | --- | --- |
| **Mass Detection** | MS1 | 1.80E+05 |
|  | MS2 | 1.00E+03 |
| **Chromatogram Builder** | Min time span (min) | 0.01 |
|  | Min height | 5.40E+05 |
|  | *m/z* tolerance (ppm) | 1.00E+01 |
|  | Filter | MS1 |
| **Chromatogram Deconvolution - Local Minimum Search** | Chromatographic Threshold | 5% |
|  | Search minimum in RT range | 0.10 min |
|  | Min relative height | 15% |
|  | Min absolute height | 3.00E+05 |
|  | Min ratio of peak top/edge | 2 |
|  | Peak duration range | 0.01-1.5min |
|  | *m/z* range for MS2 scan pairing | 0.01 |
|  | RT range for MS2 scan pairing | 0.1min |
| **Isotopic Peaks Grouper** | *m/z* tolerance (ppm) | 1.00E-02 |
|  | Retention time tolerance (min) | 0.05 |
|  | Max charge | 3 |
|  | Representative isotope | Lowest *m/z* |
|  | Monotonic shape | Yes |
| **Join Aligner** | *m/z* tolerance (ppm) | 10 |
|  | Weight for *m/z* | 5 |
|  | Weight for RT | 1 |
|  | Retention time tolerance (min) | 0.5 |
| **Peaks List Row Filter** | Retention time (min) | 0.2-12.51 |
|  | Min peaks per row | empty (leave unchecked) |
|  | Keep only peaks with MS2 scan (GNPS) | yes |
| **Gap Filling Peak Finder** | Intensity tolerance (%) | 75 |
|  | *m/z* tolerance (ppm) | 1.00E+01 |
|  | *m/z* tolerance (*m/z*) | 1.00E-06 |
|  | Retention time tolerance (min) | 0.3 |
|  | RT correction | yes |

## Acknowledgements

We thank our collaborators with the Communidad Native Matses Anexo San Mateo, Caserío de Tunapuco, Centre MURAZ Research Institute, and the Ministry of Health in Burkina Faso for their collaboration and for opening their communities to our research. We thank Dr. Marielle Hoefnagels and students of the OU BioWriting class for their assistance with editing and reviewing the manuscript.

## Author Contributions

C.M.L., L.-I.M., and K.S. conceived and designed the study. C.M.L., A.J.O.-T., R.Y.T., L.M.-R., E.G.-P., and L.T.-C. led Peruvian sample collection and developed ethical guidelines for community engagement. T.S.K. led fieldwork, metadata curation, and sample processing in Burkina Faso and contributed to lab work in the U.S. D.J. assisted with fieldwork in Burkina Faso, metadata curation, and conducted data analysis. L.-I.M. directed all LC-MS/MS experimentation and data analyses. J.J.H., E.H., and L.-I.M. acquired LC-MS/MS data. J.J.H. and L-I.M. performed LC-MS/MS data analysis with contributions from M.K., A.R.P., and K.F. J.J.H. wrote the manuscript with contributions from L.-I.M. and C.M.L. All authors reviewed the final manuscript.

## Competing Interests

The authors declare no conflicts of interest.

## Funding

This study was supported by grants from the National Institutes of Health (NIH R01 GM089886) and National Science Foundation (Doctoral Dissertation Improvement Grant 1925579).

## Notes

### Competing Interest Statement

The authors have declared no competing interest.

